# Lineage-specific genomic imprinting in the *ZNF791* locus

**DOI:** 10.1101/2024.07.13.603355

**Authors:** Jinsoo Ahn, In-Sul Hwang, Mi-Ryung Park, Milca Rosa-Velazquez, In-Cheol Cho, Alejandro E. Relling, Seongsoo Hwang, Kichoon Lee

## Abstract

**Background:** Genomic imprinting is the primary epigenetic phenomenon that results in parent-of-origin effects on mammalian development and growth. Research on genomic imprinting in domesticated animals has lagged due to a primary focus on orthologous imprinted genes. This emphasis has limited the discovery of imprinted genes specific to livestock. To identify genomic imprinting in pigs, we generated parthenogenetic porcine embryos alongside biparental normal embryos, and then performed whole-genome bisulfite sequencing and RNA sequencing on these samples.

**Results:** In our analyses, we discovered a maternally methylated differentially methylated region within the orthologous *ZNF791* locus in pigs. Additionally, we identified both a major imprinted isoform of the *ZNF791-like* gene and an unannotated antisense transcript that has not been previously annotated. Importantly, our comparative analyses of the orthologous *ZNF791* gene in various eutherian mammals, including humans, non-human primates, rodents, artiodactyls, and dogs, revealed that this gene is subjected to genomic imprinting exclusively in domesticated animals, thereby highlighting lineage-specific imprinting. Furthermore, we explored the potential mechanisms behind the establishment of maternal DNA methylation imprints in porcine and bovine oocytes, supporting the notion that integration of transposable elements, active transcription, and histone modification may collectively contribute to the methylation of embedded intragenic CpG island promoters.

**Conclusions:** Our findings convey fundamental insights into molecular and evolutionary aspects of livestock species-specific genomic imprinting and provide critical agricultural implications.

## Background

Genomic imprinting is the primary epigenetic phenomenon that results in parent-of-origin-dependent gene expression in offspring. A subset of mammalian genes is affected by genomic imprinting, which fundamentally contributes to normal development and growth [1]. Epigenetic imprints, such as DNA methylation, are differentially established in the parental germ cells, i.e., oocytes and sperm. After fertilization, if these primary germline differentially methylated regions (gDMRs) resist genome-wide demethylation during pre-implantation development, they can serve as imprinting control regions (ICRs) [2]. Subsequently, ICRs dictate monoallelic expression depending on whether the allele is inherited from the mother or the father, leading to the formation of imprinted gene clusters in the majority of cases involving multiple coding and noncoding genes [3] or micro-imprinted domains in fewer cases in which surrounding genes escape imprinting [4].

The DMRs show spatially distinctive patterns. Sperm-derived gDMRs/ICRs are predominantly found in intergenic regions. In contrast, oocyte-derived gDMRs/ICRs, often located at CpG island (CGI) promoters, are associated with transcriptional repression that drives the silencing of the maternal allele and the expression of the paternal allele of corresponding genes [5]. Therefore, in this case of direct silencing, the paternal allele-specific expression is accompanied by a maternally methylated gDMR (hereafter, maternal gDMR) that originates from oocytes and is maintained throughout zygotic and post-implantation embryonic development. Since the discovery of genomic imprinting, molecular biology approaches have been applied to identify these gDMRs and the allelic expression of putative imprinted genes, utilizing single nucleotide polymorphisms (SNPs). It has been found that genomic imprinting occurs in therian mammals, including eutherians and marsupials, while there is no evidence of genomic imprinting in prototherians (monotremes), avians (birds), and lower vertebrates such as reptiles, amphibians, and fish [6].

Research on genomic imprinting in domesticated mammals has primarily targeted orthologous genes known to be imprinted in mice and humans [7]. This focus may restrict the identification of livestock-specific imprinted genes that do not have imprinted orthologs in mice and humans. Importantly, the generation of diploid parthenogenetic or gynogenetic embryos, which contain two maternal genomes in place of the paternal genome, leads to early lethality in mouse embryos, underscoring the necessity of both paternal and maternal genomes for normal embryonic development [8]. However, these types of uniparental embryos tend to survive longer in domesticated animals [9]. Based on this, our research focused on generating parthenogenetic porcine embryos and comparing them with biparental embryos to identify maternal imprints that have not been previously reported in mice and humans [10]. Utilizing these embryos, we aim to investigate a maternal DMR located at a CGI promoter, alongside assessing concurrent differential gene expression patterns. Specifically, analyzing imprinting states at the transcript level will allow us to gain detailed insights into the imprinting status in a locus. Further comparative analyses in other mammalian species, including other domesticated mammals, humans, non-human primates, and rodents, will serve to highlight the evolutionary conservation or non-conservation of imprinting, as well as explore potential mechanisms of *de novo* DNA methylation.

In this study, we analyzed porcine zinc finger protein genes, which have been reported to play roles in maintenance methylation [11], but whose imprinting statuses have yet to be comprehensively studied at the transcript level. We highlight the zinc finger protein 791-like (*ZNF791-like*) gene as an imprinted gene in pigs, which has not been extensively studied. Using whole-genome bisulfite sequencing (WGBS) and RNA sequencing (RNA-seq) on parthenogenetic and normal biparental porcine embryos, we identified a maternal DMR, an imprinted isoform, and a previously unannotated antisense transcript in pigs. Importantly, our comprehensive analyses of the orthologous genes across eutherian mammalian species revealed that the imprinting patterns were exclusive to domesticated mammals and absent in primates and rodents. Furthermore, we explored the potential mechanisms underlying the establishment of maternal imprints in oocytes, supporting the notion that a long terminal repeat (LTR) integration, upstream transcription initiation, and histone modification may act in concert to DNA methylation at embedded intragenic CpG island promoters. These findings describe lineage-specific *ZNF791* imprinting, offering insights into molecular and evolutionary aspects of epigenetic regulation among eutherian mammals.

## Results

### Detection of Differentially Methylated Regions and Imprinted Expression

Following the generation of parthenogenetically activated (PA) and control (CN) porcine embryos as detailed in the Materials and Methods section (Supplementary Fig. 1), we conducted WGBS (Supplementary Table 1). Subsequently, we analyzed DNA methylation levels in these pig embryos within genomic regions (between –10 kb from the TSS and +10 kb from the TES) and putative promoter regions (defined as ± 2 kb from the TSS) of all 2,900 porcine zinc finger transcripts from 642 genes. Our analysis revealed that mean DNA methylation levels in the putative promoter regions were generally low (Fig. 1A), indicative of a condition that is conducive to unbiased detection of DMRs in the promoter regions of these embryos. Moreover, methylation levels were higher in gene bodies (from TSS to TES) compared to flanking regions across embryos, aligning with the gene-body methylation pattern in mammals (Fig. 1A). Based on the methylation ratios between PA and CN embryos, the top 88 porcine zinc finger transcripts with orthologous loci in humans were identified (Supplementary Table 2). Further sorting according to the PA1 methylation levels revealed that the putative promoter regions of 21 *PLAGL1* transcripts and two *ZNF791-like* transcripts exhibited high methylation in PA embryos and moderate methylation in CN embryos (Fig. 1B). This suggests a maternal methylation pattern, where PA embryos with only maternal alleles tended to have higher methylation levels than the control embryos.

**Figure 1.**
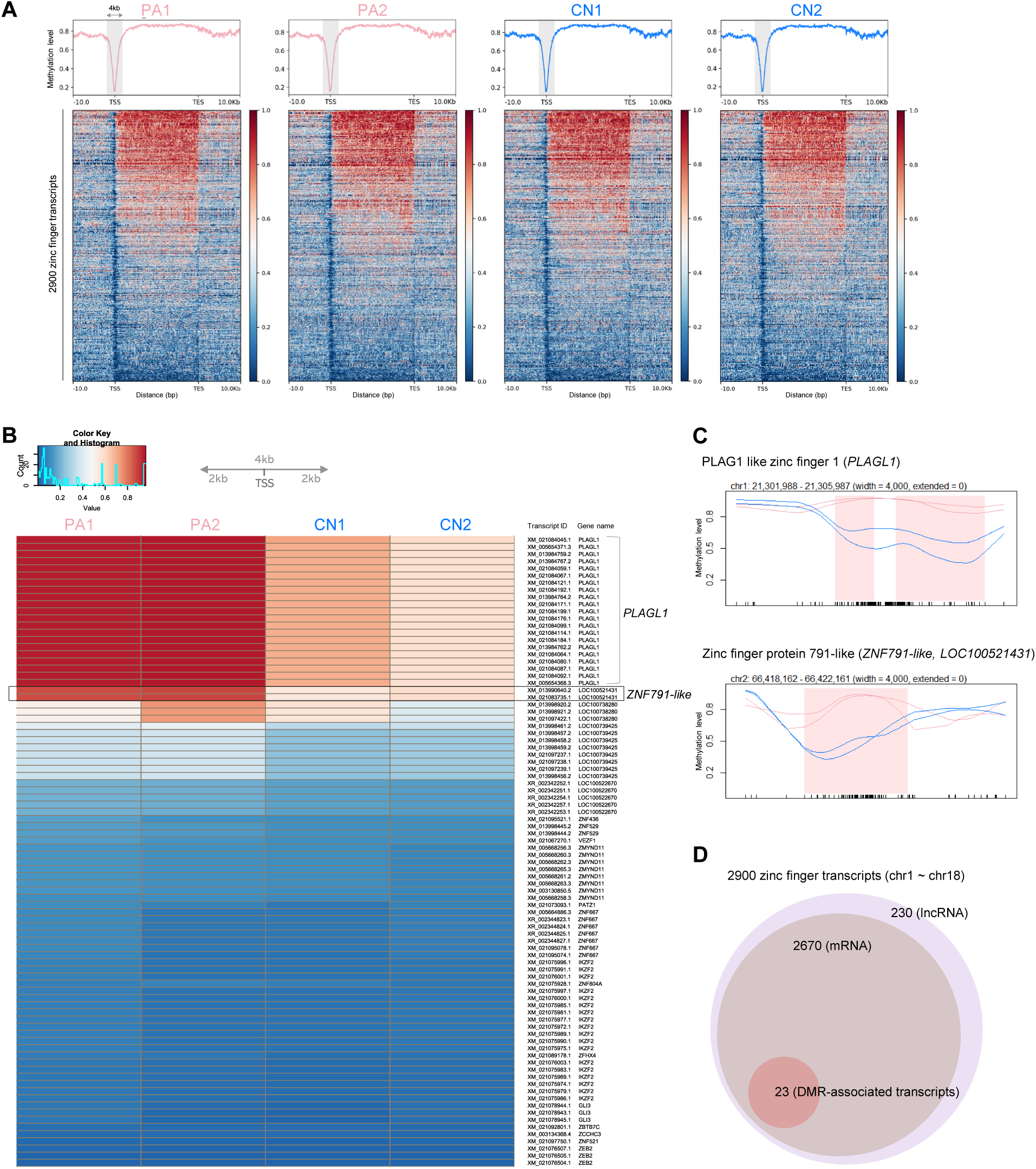
Overview of imprinted transcript discovery. **(A)** Mean DNA methylation levels of genomic regions in porcine embryos. The analyzed genomic regions include transcript bodies (TSS – TES) and ± 10 kb (–10 kb from TSS and +10 kb from TES). The ± 2 kb region from TSS is highlighted with grey. DNA methylation levels of all 2,900 porcine zinc finger transcripts from autosomes (chr1 ∼ chr18) are presented with heatmaps in descending order. TSS, transcription start site; TES, transcription end site. **(B)** A heatmap of mean DNA methylation levels at CpG sites within ± 2 kb of the TSS of 88 selected transcripts. Based on methylation ratios between PA and CN embryos, the top 88 porcine transcripts having human orthologous loci were identified and re-sorted by methylation levels in PA1 embryo. **(C)** Smoothed methylation profiles for PA replicates (pink lines) and CN replicates (blue lines). Identified DMRs (pink-colored regions) for two genes using the BSmooth method from bsseq are displayed. CpG sites are denoted at the bottom of the plots. **(D)** Numbers of analyzed porcine zinc finger transcripts. Out of 2,900 porcine zinc finger transcripts, 23 mRNA transcripts from the *PLAGL1* and *ZNF791-like* genes were identified as DMR-associated transcripts based on our criteria.

We then employed a DMR detection software to identify differentially methylated regions between the PA and CN embryos within the putative promoter regions. Using the BSmooth method with default parameters, candidate DMRs were identified in the promoter regions (TSS± 2 kb) with dense CpG sites of two zinc finger protein genes, *PLAGL1* and *ZNF791-like* (Fig. 1C). Among 2,900 porcine zinc finger transcripts, there are 2,670 mRNAs and 230 long-noncoding RNAs (lncRNA), and 23 mRNA transcripts of *PLAGL1* and *ZNF791-like* were identified to be associated with DMRs in their putative promoter regions (Fig. 1D). The known imprinted locus of *PLAGL1* was utilized as a positive control for the maternally methylated DMR. This DMR was further validated using the metilene software (FDR < 0.05) at the promoter region of expressed transcripts, and the corresponding paternal expression pattern was exclusively observed in the control embryos (Supplementary Fig. 2). Another DMR identified upstream by metilene was more than 3kb apart from the TSS of XM_021084156.1, falling outside the TSS ± 2 kb range (Supplementary Fig. 2). In summary, differentially methylated regions were identified within the putative promoter regions of zinc finger transcripts at the porcine *ZNF791-like* locus, as well as at the *PLAGL1* locus, which served as a positive control.

### Locus-Specific Paternal Expression of *ZNF791-like* was Attributed to a Major Transcript Isoform

Following the current annotation of pig reference genome assembly (Sscrofa11.1/susScr11), the *LOC100521431* (*ZNF791-like*) locus was explored to reveal the imprinting status. In the downstream (3′) of *MAN2B1* gene, short transcripts (XM_031990640.2 and XM_021083735.1) of *ZNF791-like* appeared to be expressed from the CN embryos exclusively that are initiated from a CpG island (CGI) (Fig. 2A,B). In addition to this NCBI genome annotation, we integrated the *ENSSSCG00000033388* gene from the Ensembl genome annotation in the upstream (5′) of long transcripts of *ZNF791-like*. This Ensembl transcript is initiated from a CGI and appeared to be expressed from both PA and CN embryos. There also appeared to be an expression of previously unannotated antisense (AS) transcript overlapping the short transcripts of *ZNF791-like* (Fig. 2A,B). Our profiling of DNA methylation status revealed a DMR spanning the CGI that regulates transcription of the short *ZNF791-like* transcripts (Fig. 2C). This DMR resulted from full methylation in PA embryos and partial methylation in CN embryos, indicating maternal DNA methylation (i.e., maternal imprint) that could lead to silencing of the maternal allele of short *ZNF791-like* transcripts and their subsequent paternal expression. In contrast, the CGI regulates transcription of the *ENSSSCG00000033388* gene was unmethylated in both PA and CN embryos (Fig. 2C).

**Figure 2.**
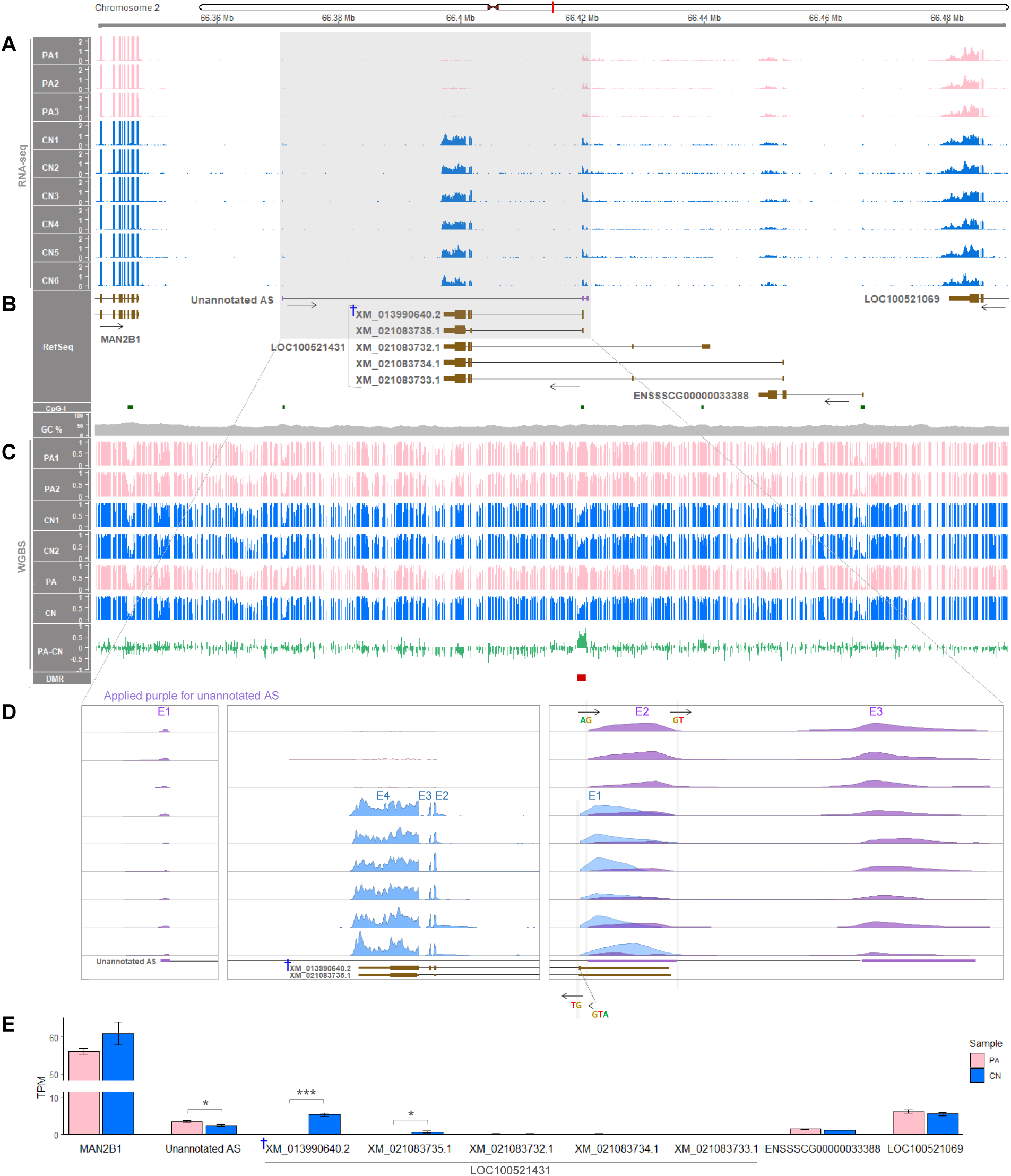
Regional view of the LOC100521431 (ZNF791-like) locus in pigs. **(A)** RNA-seq read coverages of PA (pink, n = 3) and CN (blue, n = 6) embryos. The 15-Kb region between 66.34 Mb (66,340,000) and 66.49 Mb (66,490,000) is depicted. Read coverages were normalized to TPM values. **(B)** The NCBI RefSeq annotation is displayed along with an Ensembl-derived transcript (ENSSSCG00000033388) and an unannotated transcript [unannotated antisense (AS) depicted in purple]. For protein-coding transcripts which are depicted in brown, tall boxes represent translated regions and short boxes represent untranslated regions. Transcriptional direction is indicated with black arrows. CpG-I, CpG island; GC%, GC content. **(C)** WGBS DNA methylation levels (n = 2 for each of PA embryos and CN embryos). Mean methylation ratios (PA and CN) are followed by mean methylation differences (PA-CN). A DMR (FDR < 0.05) is marked in red. **(D)** A close view of expressed transcripts with exon (E) numbers. Splicing donor (GT) and acceptor (AG) sites are denoted with directional arrows to indicate their orientation. The start codon (ATG) marks the beginning of a protein-coding open reading frame (ORF). **(E)** Quantification of expressed transcripts and genes. TPM values are represented as mean ± SEM. *, *P* < 0.05; ***, *P* < 0.001; †, a major predominant *ZNF791-like* transcript.

A zoomed visualization of the short *ZNF791-like* transcripts showed prominent exon 3 (E3) usage, instead of E3 skipping, as seen in the transcript XM_031990640.2 (Fig. 2D). A combined close view revealed that exon 2 (E2) of the unannotated AS overlapped with exon 1 (E1) of the short *ZNF791-like* transcript (Fig. 2D). The splicing of the unannotated AS was supported by the presence of splicing donor (GT) and accepter (AG) sites, allowing the inference of its transcriptional direction based on the orientation of the GT and AG sequences (Fig. 1D and Supplementary Fig. 3). Of note, expression of the unannotated AS tended to be higher in PA embryos than in CN embryos. Further quantification of transcript expression revealed that the mean expression level of the major short *ZNF791-like* transcript (XM_031990640.2) in the CN embryos was approximately 8-fold higher than that of another short transcript (XM_021083735.1): 5.36 ± 0.40 TPM for XM_031990640.2 and 0.66 ± 0.23 TPM for XM_021083735.1 (Fig. 2E). Notably, expression of the unannotated AS was significantly 1.43-fold higher in the PA embryos than in the CN embryos (*P* < 0.05): 2.43 ± 0.22 TPM in CN and 3.49 ± 0.25 in PA embryos (Fig. 2E), suggesting a potential maternally biased expression. This could further support the *ZNF791-like* imprinting and paternal expression, which might affect the expression of a nearby overlapping gene. In short, the paternal expression of *ZNF791-like* was greatly due to one of the short transcripts (XM_031990640.2), that could be regulated by the DMR, and potentially led to greater expression of the overlapping unannotated AS transcript in the PA embryos than in the CN embryos.

### DNA Methylation Imprint in the *ZNF791* Locus was Species-Specific

To examine conservation of *ZNF791* imprinting in mammalian species, we analyzed publicly available WGBS data (listed in Supplementary Table 3) from gametes, tissues, and cells of 13 species. In humans, mice, and rats, the *ZNF791* or *Zfp791* promoter regions were unmethylated or significantly hypomethylated in both oocytes and sperm, indicating biallelic transcriptional initiation (Fig. 3A). On the other hand, in pigs and cows, the promoter CGIs were fully methylated in oocytes and unmethylated in sperm, indicating that maternal silencing and paternal monoallelic transcriptional initiation of *ZNF791* are conserved in porcine and bovine gametes (Fig. 3A). To further investigate maintenance of methylation patterns, various tissues and cells were analyzed. In humans, non-human primates (crab-eating macaque, chimpanzee, rhesus monkey, and gibbon), and rodents (mice and rats), all the CGI promoters or the promoter regions were unmethylated or significantly hypomethylated in various fetal and adult tissues (Fig. 3B).

**Figure 3.**
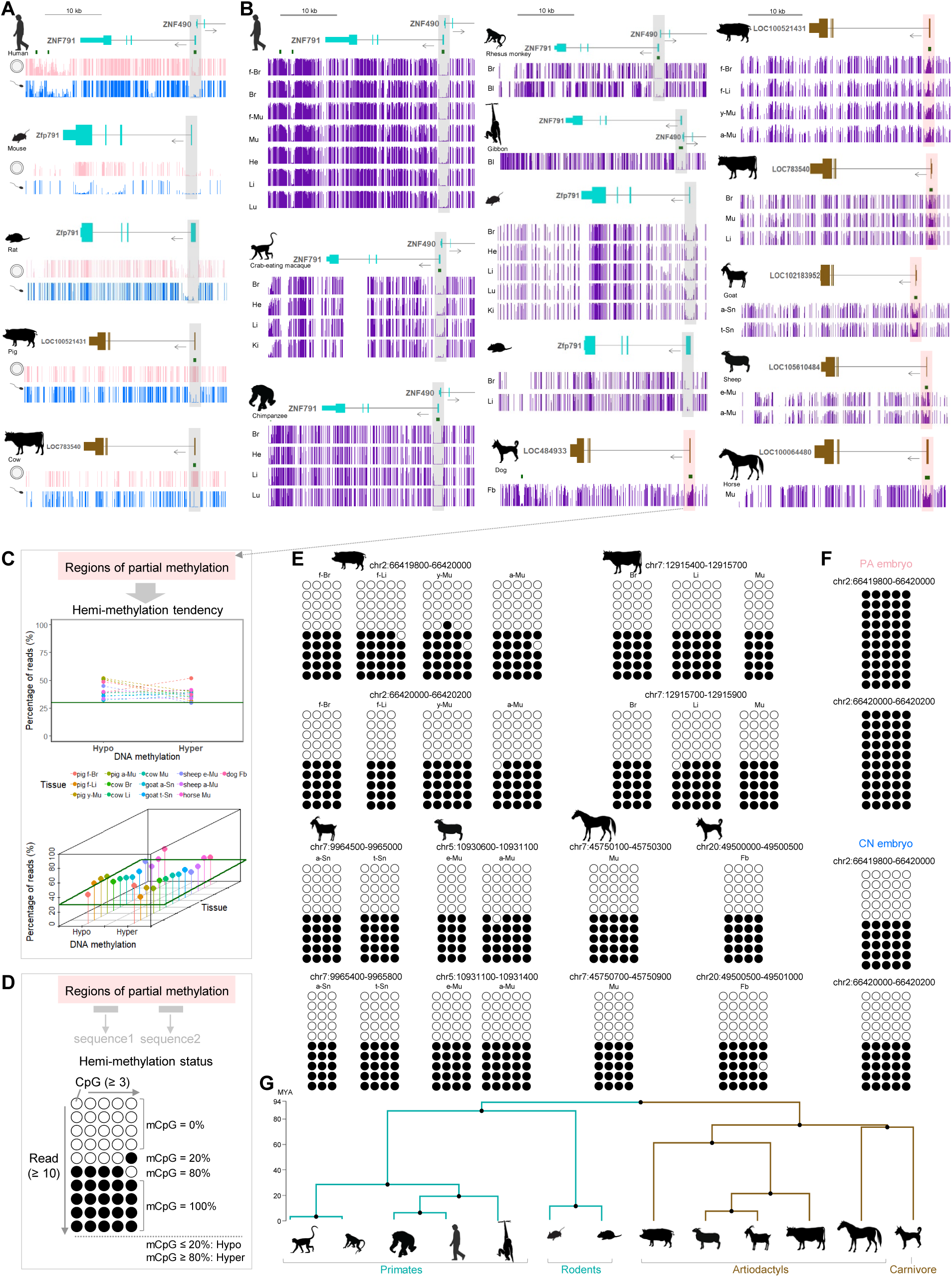
Comparison of DNA methylation status of the ZNF791 locus in mammalian cells and tissues. **(A)** DNA methylation levels, ranging from 0 (unmethylation) to 1 (full methylation), in oocytes (pink) and sperm (blue) across species: human, mouse, rat, pig, and cow. Transcripts are color-coded to distinguish between species groups: cyan for humans, mice, and rats, and brown for pigs and cows. The promoter regions of *ZNF791* (or *Zfp791*) and *ZNF791-like* are highlighted with grey shading. CpG islands are marked with green rectangles. **(B)** DNA methylation levels in various tissues or cells. Transcripts are also color-coded with cyan for primates and brown for dogs, goat, sheep, and horses. The promoter regions of *ZNF791* (or *Zfp791*) are highlighted with grey shading. The promoter regions of *ZNF791-like*, which exhibited partial methylation, are highlighted with pink shading. f-Br, fetal brain; Br, brain; f-Mu, fetal muscle; Mu, muscle; He, heart; Li, liver; Lu, lung; Ki, kidney; Bl, blood; Fb, fibroblast; f-Li, fetal liver; y-Mu, young skeletal muscle (d40); a-Mu, adult skeletal muscle (d180); a-Sn, angen stage of skin; and t-Sn, telogen stage of skin; e-Mu, embryonic muscle (embryonic day 110); a-Mu, adult skeletal muscle (2-years-old). **(C)** Analyses of the partial methylation. WGBS reads for the partial methylation region were extracted and analyzed for hemi-methylation tendency. A threshold was set as the presence of at least 30% of hypomethylated reads and hypermethylated reads. Hypomethylated or hypermethylated reads are defined in D. **(D)** Hemi-methylation status was further evaluated in two sequences from the regions of partial methylation. Criteria were set as follows: more than or equal to ten reads (Read ≥ 10) and more than or equal to three consecutive CpG dinucleotides on each read (CpG ≥ 3). The percentage of methylated CpG dinucleotides (mCpGs) was then calculated. Reads were defined as hypomethylated if the percentage of mCpG was less than 20% and as hypermethylated if it was more than 80%. Further details on the criteria are provided in the Materials and Methods section. **(E)** Evaluation of hemi-methylation status in each species. WGBS reads were extracted from indicated regions for each tissue or cell. Open and closed circles indicate unmethylated and methylated CpGs, respectively. **(F)** Full- and hemi-methylation states in PA and CN embryos, respectively. **(G)** A phylogenetic time tree showing estimated divergence time. MYA, million years ago.

To the contrary, in domesticated mammals (pigs, cows, goat, sheep, horses, and dogs), all the CGI promoters were partially methylated in various fetal and adult tissues and cells (Fig. 3B). Whether the partial methylation is originated from methylation difference between parental alleles (i.e., hemi-methylation pattern) could be analyzed based on WGBS reads. All reads overlapping the partially methylated regions tended to be either hypomethylated or hypermethylated, while the full methylated region in PA embryos showed hypermethylation only (Fig. 3C, Supplementary Figs. 4∼10, and Supplementary Table 4). Our definition of hypo- or hypermethylation is the percentage of methylated CpG dinucleotides (mCpGs) less than 20% or more than 80% and is consistent with previous approaches [12] (Fig. 3D). In at least two consecutive CpG dinucleotide sequences within reads overlapping the partially methylated regions, the percentage of mCpGs were either 0 ∼ 20% (hypomethylation) or 80 ∼ 100% (hypermethylation) in analyzed individual reads, indicating hemi-methylation status (Fig. 3E). Estimated divergence time between clade 1 (primates and rodents) and clade 2 (artiodactyls and carnivore) was *circa* 94 million years ago (MYA) (Fig. 3F). Consequently, DNA methylation imprints in the *ZNF791* or *ZNF791-like* locus appeared in one group of the analyzed mammalian species (artiodactyls and carnivore), but not in another group (primates and rodents), indicating a species-specific imprinting pattern.

### Bi or Mono-Allelic Expression of the *ZNF791* Gene Across Species Based on Individual-Matched Genomic DNA and RNA-Seq Reads

With the aim of investigating whether *ZNF791* expression is non-imprinted or imprinted, its biallelic or monoallelic expression was to be examined across mammalian species. To achieve this, we first identified single nucleotide polymorphisms (SNPs) where two different alleles exist (i.e., heterozygous SNPs), except for inbred mice. In humans, rhesus monkeys, chimpanzees, dogs, pigs, and cows, heterozygous SNPs were identified on genomic DNAs covering exons of *ZNF791* (Fig. 4 and Supplementary Table 5). Whether *ZNF791* is expressed from both alleles or from a single allele was determined by analyzing the expression of alleles at the SNP sites in various normal tissues and cells. In RNA-seq data derived from the same individual whose genomic DNA was sequenced, both alleles were expressed in humans, rhesus monkeys, and chimpanzees in analyzed transcripts (Fig. 4, Supplementary Table 5, and Supplementary Figs. 11∼12). On the other hand, in dogs, pigs, and cows, either a single alternate or reference allele was expressed in predominantly expressed isoforms in all analyzed tissues (Fig. 4, Supplementary Table 5, and Supplementary Fig. 13). Additionally, the unannotated AS was expressed in the liver of a 60-days-old pig (Supplementary Fig. 13C). In contrast, due to the presence of homozygous alleles at each SNP in inbred mice, a different approach was applied for the mouse analysis. It revealed that offspring from both initial and reciprocal crosses between CAST/EiJ and C57BL/6J strains consistently expressed both alleles in expressed transcripts (Fig. 4, Supplementary Table 5, and Supplementary Fig. 12D). In summary, *ZNF791* was found to be expressed biallelically in humans, non-human primates, and mice, indicating a non-imprinted expression in these species. Conversely, in dogs, pigs, and cows, *ZNF791* or *ZNF791-like* exhibited monoallelic expression at the heterozygous SNP sites, indicative of an imprinted expression in these mammalian species.

**Figure 4.**
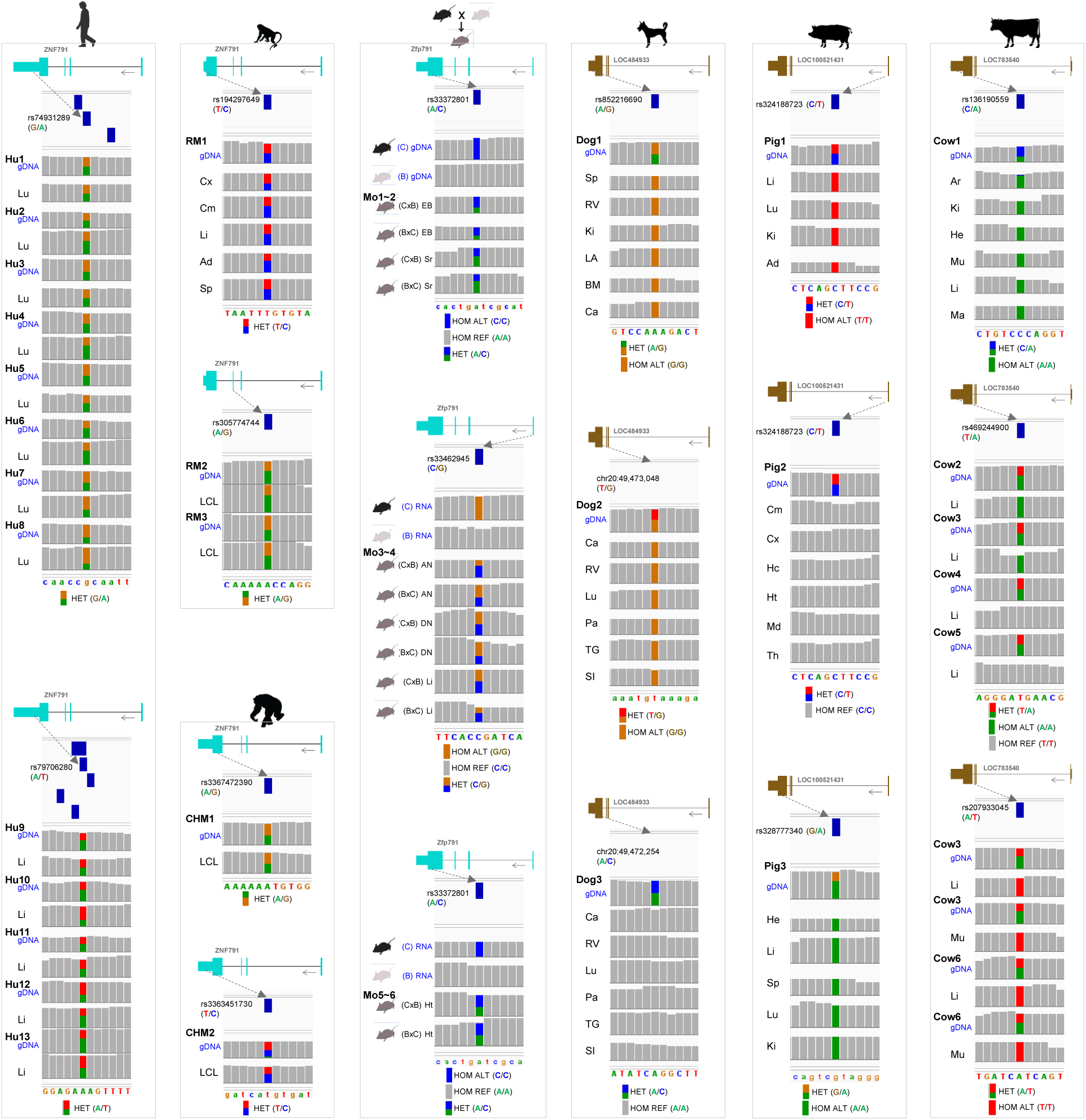
Bi- or monoallelic expression of ZNF791 or ZNF791-like across mammalian species. i) By analyzing whole genome or exome sequencing data, SNPs with two different DNA bases (i.e., heterozygous SNPs) were selected as informative SNPs to analyze corresponding bi- or monoallelic mRNA expression. For inbred mouse strains, SNPs with two identical DNA bases (i.e., homozygous SNPs) were chosen to analyze bi- or monoallelic mRNA expression in offspring resulting from reciprocal crosses. ii) In the same individuals, mRNA expressions in normal tissues or cells were analyzed to determine bi- or monoallelic expression pattern of *ZNF791* or *ZNF791-like*. The reference SNP identifiers (rs IDs) are accompanied by the reference and alternate alleles, denoted as ‘ref/alt’. For SNPs that have not been assigned rs IDs, genomic coordinates are provided, with the reference and alternate alleles specified. Hu, humans; RM, rhesus monkey; CHM, chimpanzee; Mo, mouse. For the mouse, reciprocal crosses of CAST/EiJ (C) and C58BL/6J (B) were analyzed. Tissues include: Lu, lung; Li, liver; Cx, cortex; Cm, cerebellum; Ad, adipose tissue; Sp, spleen; LCL, lymphoblastoid cell line; EB, embryonic brain; Sr, striatum; AN, arcuate nucleus; DN, dorsal raphe nucleus; Ht, hypothalamus; RV, right ventricle; Ki, kidney; LA, left atrium; BM, bone marrow; Ca, cartilage; Pa, pancreas; TG, thyroid gland; SI, small intestine; Hc, hippocampus; Md, midbrain; Th, thalamus; He, heart; Ar, adrenal cortex; Mu, skeletal muscle; Ma, mammary gland. Accession numbers of datasets, run IDs, and individual IDs are listed in Supplementary Table 5. HET, heterozygous allele; HOM REF, homozygous reference allele; HOM ALT, homozygous alternate allele.

### LTR-Mediated Transcription might Regulate *ZNF791* Imprinting in Pigs and Cattle

To investigate potential mechanisms behind the imprinting of the maternal allele of *ZNF791* gene, we analyzed transcription in oocytes and histone modifications during early development. Because the CGI promoter of *ZNF791* was maternally imprinted (Figs. 2 and 3), there could be oocyte-specific events that contribute DNA methylation. In fully grown porcine oocytes, the *ZNF791-like* transcript (NCBI accession no. XM_021083732.1) was expressed, initiating from the transcription start site (TSS) located near another putative CGI promoter and the LTR sequence, LTR52 (Fig. 5A, B, grey shading, and Supplementary Figs. 14∼15). This LTR52 was identified through a search in the Dfam database (E-value = 3.7 × 10^−5^), which uses profile hidden Markov models (HMMs) to improve the sensitivity of detection (Supplementary Fig. 16). Additionally, H3K4me3, a hallmark of promoters, was found enriched in the CGI promoter region of this transcript XM_021083732.1 (Fig. 5C, grey shading). From the site where H3K4me3 enrichment ended, both H3K36me2 and H3K36me3 tended to be enriched, supporting their role in *de novo* gene-body methylation in oocytes (Fig. 5C, D). During zygotic development (4 cell, 8 cell, and blastocyst), H3K4me3 became enriched around the intragenic CGI at the blastocyst stage, indicating transcriptional permissiveness of the transcripts XM_013990640.2 and XM_021083735.1 (Fig. 5B, † and Fig. 5C, red shading).

**Figure 5.**
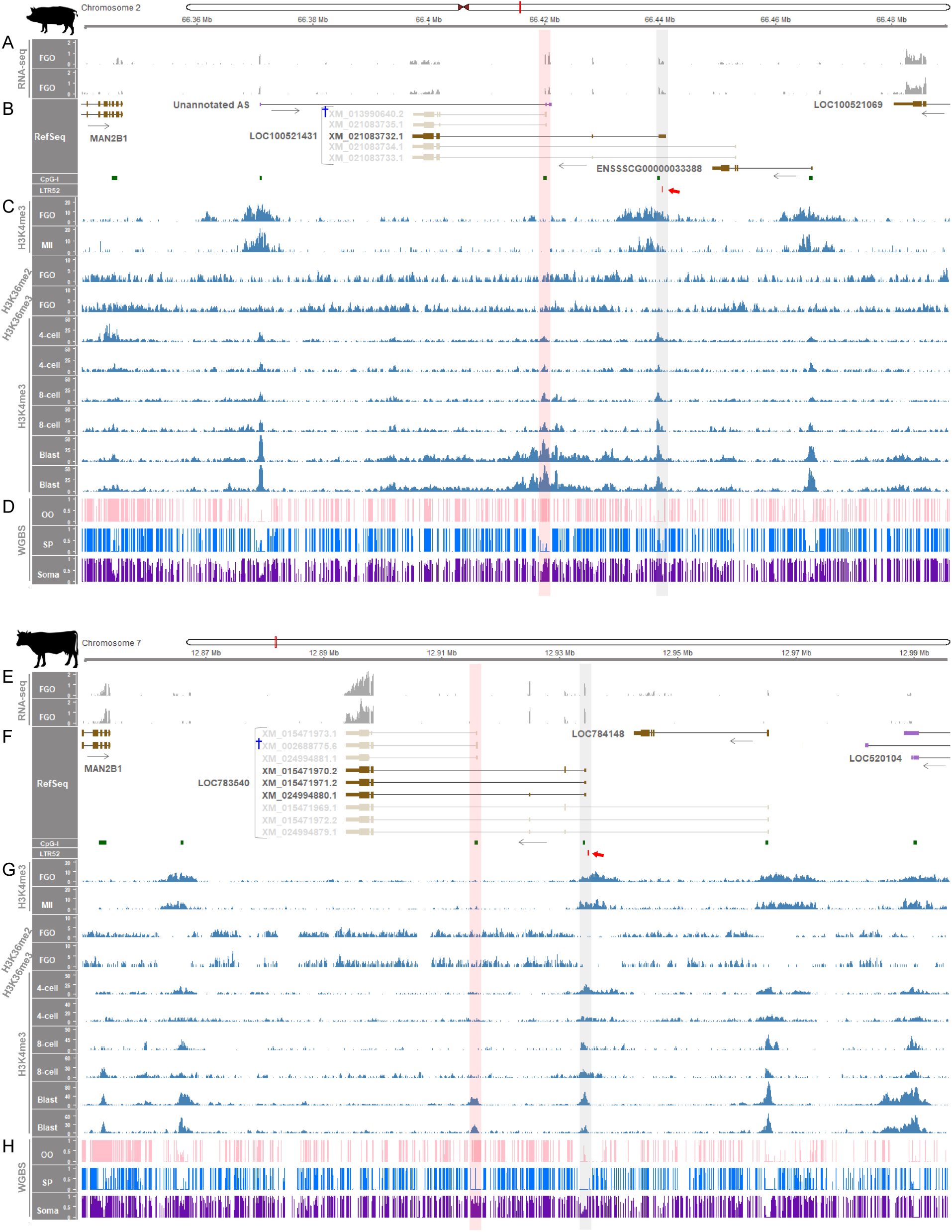
Establishment of methylation imprints in porcine and bovine oocytes. **(A, B)** RNA-seq read coverages (in TPM) from porcine fully grown oocytes (FGOs) and the expressed *ZNF791-like* transcript. The CpG-island (CpG-I) and LTR52, located around the transcription start site (TSS; chr2:66,440,912), are marked with green and red rectangles, respectively, and both are highlighted with grey shading; the LTR52 is further indicated by the red arrow. Non-expressed transcripts are faded out. **(C)** Enrichment of histone modifications, H3K4me3 and H3K36me2/me3, during porcine pre-implantation development. **(D)** WGBS methylation ratios of porcine samples. Full methylation in oocytes (OO), unmethylation in sperm (SP), and partial methylation in somatic tissue (d120 muscle) are highlighted with red shading. **(E, F)** RNA-seq coverages from bovine FGOs and expressed *ZNF791* transcripts. The CpG-I and LTR52 located around the TSS (chr7:12,934,209), are marked with green and red rectangles, respectively, and both are highlighted with grey shading; the LTR52 is further pointed out by the red arrow. Faded out are non-expressed transcripts. **(G)** Histone modifications, H3K4me3 and H3K36me2/me3, during bovine pre-implantation development. **(H)** WGBS methylation ratios of bovine samples. Full methylation, unmethylation, and partial methylation in OO, SP, and somatic tissue (muscle), respectively, are highlighted with red shading. MII, MII stage oocytes; 4-cell, 4-cell stage embryos; 8-cell, 8-cell stage embryos; Blast, blastocysts. From the red-shaded region, the major predominant *ZNF791* transcripts, marked with † in (B) and (F), are preferentially transcribed at later developmental stages.

In bovine oocytes, the *ZNF791* transcripts (XM_015471970.2, XM_015471971.2, and XM_024994880.1) were expressed, initiating from TSSs located near a putative CGI promoter and an LTR52 element (E-value = 1.2 × 10^−11^) (Fig. 5E, F, grey shading, and Supplementary Figs. 16∼17). Enrichment of H3K4me3 further supported the active transcription of those transcripts (Fig. 5G, grey shading). From the site where H3K4me3 terminated, H3K36me2/me3 enrichment began in oocytes, indicating their role in *de novo* gene-body DNA methylation (Fig. 5G, H). The demarcation of H3K36me2/me3 (i.e., the formation of a boundary) was somewhat more pronounced in bovine oocytes compared to porcine oocytes. As zygotes developed, H3K4me3 became enriched at the intragenic CGI during the blastocyst stage (Fig. 5G, red shading)

Overall, the patterns of transcription initiation, histone modification, and DNA methylation were similar between porcine and bovine oocytes and developing zygotes. Taken together, these data suggest a mechanism whereby transcription was initiated near the LTR element during oogenesis, leading to the recruitment of DNA methyltransferase, subsequent *de novo* gene-body DNA methylation at the intragenic CGI, and the establishment of the DMR.

### The *ZNF791* Locus in Humans and Mice Lacks Genomic Imprinting

The imprinting mechanism described above was previously identified for the *Impact* gene which is imprinted in mice (Supplementary Fig. 18) and rats but not in humans and pigs, indicating species-specific imprinting patterns. In contrast, the *ZNF791* (or *Zfp791*) gene exhibited no maternal imprints in both humans and mice (Fig. 3) and showed non-monoallelic expression in both species (Fig. 4), indicating a non-imprinted status. To confirm the absence of genomic imprinting at this locus, we further explored the *ZNF791* (or *Zfp791*) region. In human oocytes, the only annotated transcript of *ZNF791* was expressed, to a lesser degree than the *ZNF490* transcript, which is transcribed in the opposite direction (Fig. 6A, B). Also, H3K4me3 was consistently enriched in the CGI promoter region during both gametic and zygotic stages (Fig. 6C, grey shading). The absence of an additional CGI promoter in the downstream region suggests the lack of canonical maternal imprints in human oocytes (Fig. 6D).

**Figure 6.**
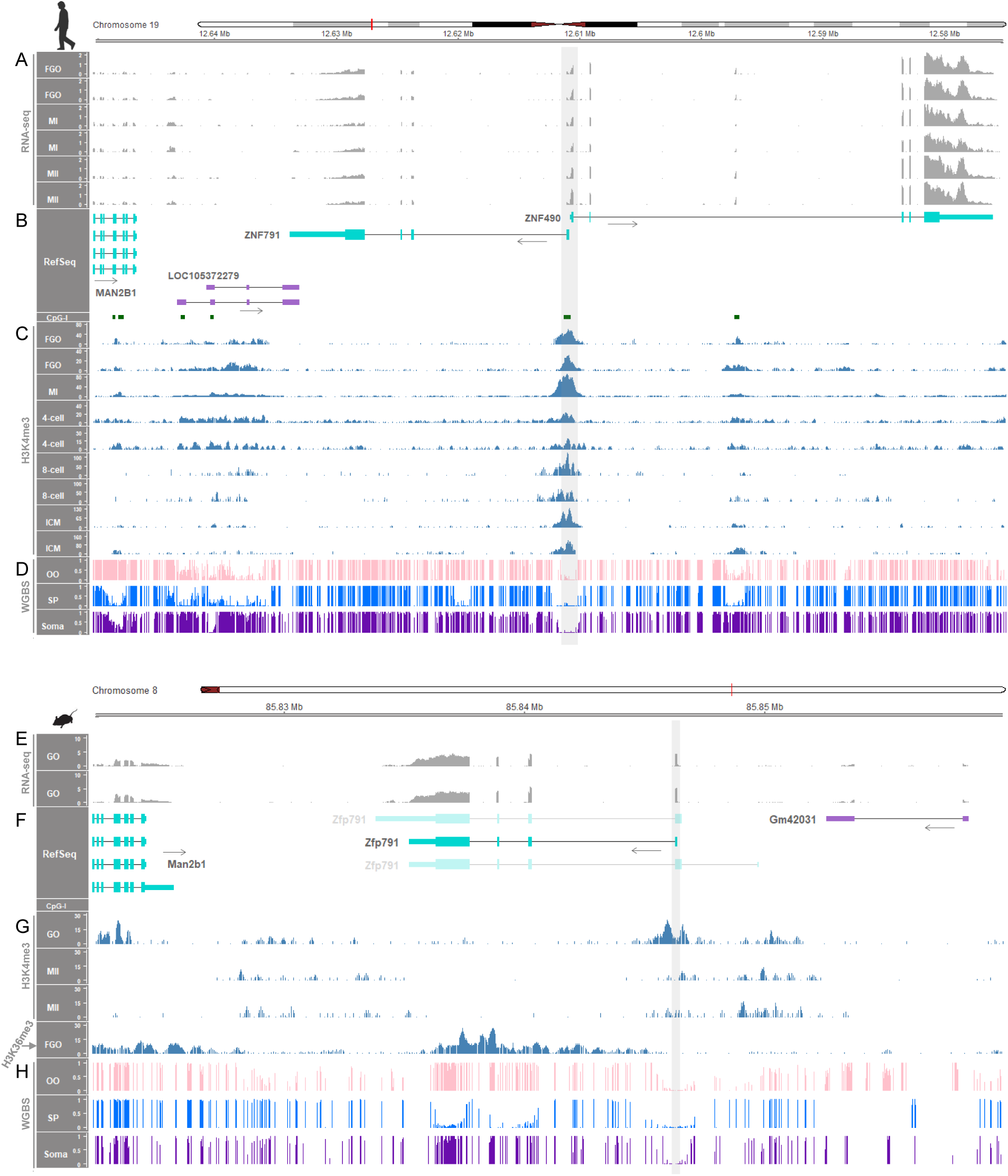
Non-establishment of methylation imprints in human and mouse oocytes. (**A, B**) RNA-seq read coverages (in TPM) from human oocytes at FGO, MI, and MII stages and the expressed *ZNF791* transcript. (**C**) H3K4me3 histone modifications during developmental stages of oocytes and zygotes. ICM, inner cell mass of the blastocyst. (**D**) WGBS methylation ratios of human samples. Unmethylation in oocytes (OO), sperm (SP), and somatic tissue (adult adipose tissue) are highlighted with grey shading. (**E, F**) RNA-seq read coverages from mouse growing oocytes (GOs) and the expressed *Zfp791* transcript. Non-expressed transcripts are presented as faded. (**G**) Histone modifications, H3K4me3 and H3K36me3, in mouse oocytes. (**H**) WGBS methylation ratios of mouse samples. Unmethylation in OO, SP, and somatic tissue (kidney) are highlighted with grey shading. All mouse samples are from the C57BL/6 strain.

In mouse oocytes, only one *Zfp791* transcript was expressed, with no other transcripts annotated to be transcribed from the downstream region (Fig. 6E, F). The promoter region was enriched with H3K4me3 histone modification (Fig. 6G, grey shading). Enrichment of H3K36me3 occurred in the gene body of *Zfp791* (Fig. 6G). The absence of a downstream CGI promoter region and additional annotated transcripts suggests a deficiency of canonical maternal imprints in mouse oocytes (Fig. 6G), in contrast to the mouse *Impact* gene, which contains the canonical maternal imprint and exhibits partial methylation in somatic tissue (Supplementary Fig. 18).

Unlike in pigs and cattle, maternal methylation imprints were not established in human and mouse oocytes, attributable to the absence of additional CGI promoters and downstream *ZNF791* transcript isoforms. This suggests that divergence between these domesticated animals and humans and mice has led to lineage-specific imprinting at the *ZNF791* locus.

### *ZNF791* Imprinting Implies Selective Forces of LTRs and Evolution of Lineage-Specific Imprinting

To further investigate orthologous imprinted *ZNF791* loci, we first compared the loci and LTRs in artiodactyls and dogs. Focusing on the *ZNF791* sequences immediately downstream (3′) of the *MAN2B1* gene in each species, we found that LTR52-derived sequences are present near the second CGI in domesticated Artiodactyla species (pigs, cows, sheep, horses, and goat), while an LTR103-derived sequence is present near the second CGI in dogs (order Carnivora) (Fig. 7A, grey shaded rectangles). E-values, or expectation values, for LTRs derived from the Dfam database were as follows: pig LTR52 (E-value = 3.7 × 10^−5^), cattle LTR52 (E-value = 1.2 × 10^−11^), sheep LTR52 (E-value = 6.8 × 10^−6^), horse LTR52 (E-value = 2.4 × 10^−6^), goat LTR52-int (E-value = 7.0 × 10^−5^), and dog LTR103_Mam (E-value = 0.051). These values indicate significant sequence similarity with the consensus sequences of LTR52 or LTR52-int (E-value < 0.05) and suggest a potential similarity with the consensus LTR103_Mam sequence in dogs. As imprinted *ZNF791* transcripts are expected to be expressed from the third CGI downstream (3′) of LTRs (Fig. 7A, red shaded rectangles) in pigs and cows (Fig. 2, Fig. 5, and Supplementary Fig. 13B,C), in dogs (Supplementary Fig. 13A), and in sheep, horses, and goats (Supplementary Fig. 19A,B), it suggests that the conservation of *ZNF791* imprinting in the analyzed species is also subject to an analogous pattern. The phylogenetic analysis of domesticated animals showed that the genetic distances between LTR52 sequences and the consensus LTR52 sequence are smaller than the distances between goat- and dog-derived sequences and the same consensus sequences (Fig. 7B and Supplementary Fig. 20). From the motif discovery/comparison and scanning, distribution of transcription factor binding sites (TFBSs) in LTR52 and its derivatives were identified (Fig. 7C). From the motif discovery and comparison, DNA binding motifs of E2F3 and E2F2 (E2F2_DBD_1 and E2F3_DBD_1) were found to be significant in LTR52 sequences from pigs, cows, sheep, and horses (E-value < 0.05, *q*-value < 0.05) (Fig. 7C,D and Supplementary Fig. 21). Also, from motif scanning, binding sites for the MYBL2 transcription factor were found in the LTR52 sequences from pigs, cows, sheep, horses, and goat_LTR52-int (*q*-value < 0.05) (Fig. 7C,D). Binding sites for the ZNF770 transcription factor were found in the LTR52 sequences from pigs, cows, sheep, horses, and dog_LTR103_Mam (*q*-value < 0.05) (Fig. 7C,D).

**Figure 7.**
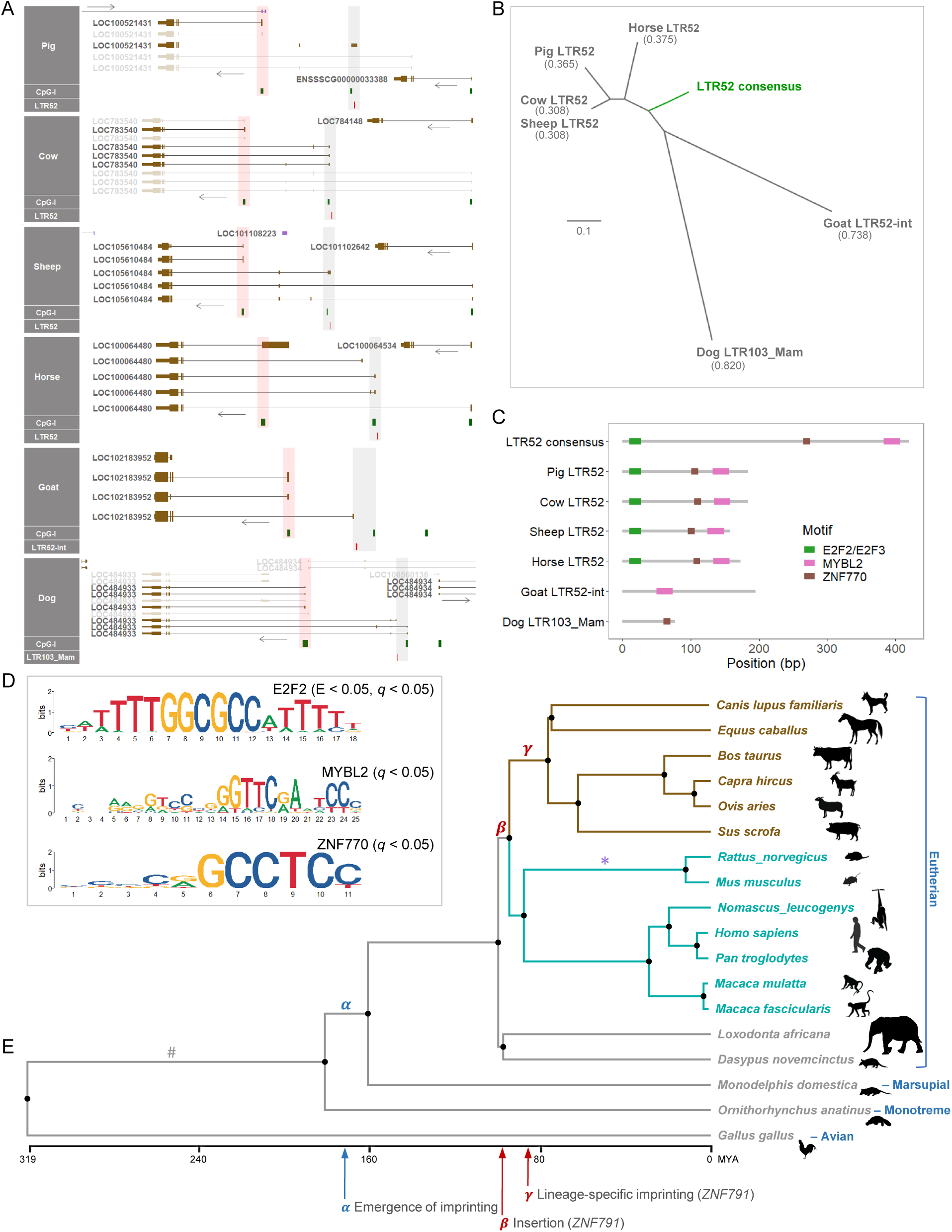
LTR52 sequence comparison across the *ZNF791* genomic regions in domesticated animals. (A) LTR locations near CpG islands are marked by red rectangles in grey shaded areas. Red shaded areas indicate the 1st exon of candidate imprinted transcripts and CpG island promoter regions. (B) Phylogenetic analyses of genetic distances between LTR sequences. The scale bar represents the evolutionary distance between sequences, with a value of 0.1 indicating a substitution per nucleotide, as estimated by the Kimura 2-parameter (K2P) model. (C) Transcription factor binding motifs detected in the analyzed LTR sequences. (D) Logos of the significant motifs. (E) Phylogenetic time tree. Species of artiodactyls and carnivores are highlighted with brown color. Primates and rodents are highlighted with cyan color. Silhouette images of these species are from phylopic.org. Potential evolutionary events are marked with letters and arrows. MYA, million years ago; #, inclusion of the *MAN2B1* gene; *, lineage-specific imprinting (*Impact*).

The phylogenetic time tree suggests that the insertion of the *ZNF791* gene likely occurred around 94 MYA (Fig. 7E, *β*), although the immediate downstream gene, *MAN2B1*, predates the emergence of genomic imprinting (Fig. 7E, #). In *Gallus gallus* (chicken, avian), the whole locus containing both the *MAN2B1* gene and the *ZNF791* gene is missing from the NCBI RefSeq database. In *Ornithorhynchus anatinus* (platypus, monotreme), *Monodelphis domestica* (gray short-tailed opossum, marsupial), *Dasypus novemcinctus* (nine-banded armadillo, edentate), and *Loxodonta africana* (African savanna elephant, proboscidean), the *MAN2B1* gene exists but the downstream *ZNF791* gene is absent. Contrary to the proposed emergence of imprinting at least 160 MYA following the divergence between monotremes and therians (marsupial and eutherian mammals) (Fig. 7E, *α*), lineage-specific imprinting of *ZNF791* might have occurred at least approximately 78 MYA in the common ancestors of artiodactyls and dogs (Fig. 7E, *γ*). This lineage-specific imprinting might have involved the acquisition of upstream LTRs, overlapping transcripts, and the maternal gDMR locus, as illustrated by the *ZNF791* locus in this study. Similarly, lineage-specific imprinting of *Impact* may have occurred relatively recently in the common ancestors of rodents, involving the acquisition of maternal gDMR processes (Fig. 7E, ∗). The *ZNF791* imprinting also leads to imprinted maternal expression of the overlapping unannotated AS in pigs (Fig. 2D), and based on the pig QTLdb, a SNP located 327 bp upstream of the 1st exon of the unannotated AS is associated with residual feed intake (RFI) at a suggestive level (*P* = 2.89E-05) (Supplementary Fig. 22), Taken together, in analyzed domesticated mammals (pigs, cows, sheep, horses, goat, and dogs), genomic imprinting at the *ZNF791* or *ZNF791-like* locus might have been selectively evolved with LTR sequence integration.

## Discussion

Herein, we conducted a comparative investigation of the imprinting statuses of *ZNF791* across 13 mammalian genomes, focusing on previously unreported DMRs, unannotated transcripts, and several key factors: CpG island promoter methylation, allele-specific expression, transcription in oocytes, integration of transposable elements, and lineage-specific imprinting. To identify candidate maternal imprints, we initially compared DNA methylation levels between bimaternal parthenogenetic embryos and biparental control embryos from pigs, utilizing two biological replicates from WGBS. This comparison facilitated the unbiased detection of maternal DMRs at CGI promoters which implicated transcriptional repression of the maternal allele [5]. Subsequently, to determine whether the maternal DMR detected in the *ZNF791-like* locus constitutes a germline DMR, we analyzed WGBS methylation levels from oocytes and sperm. Interestingly, the CGI promoters exhibited full methylation in oocytes from pigs and cattle, yet remained unmethylated in oocytes from humans, mice, and rats, indicating species-specific maternal gDMR imprints. The maintenance of allele-specific DNA methylation in somatic tissues of outbred animals were analyzed by examining methylation levels of consecutive CpG dinucleotides in individual WGBS reads (a read-based method), independent of SNPs [12]. It revealed that hemi-methylation (i.e., reflection of full methylation in one of the parental alleles and unmethylation in the other) at the CGI promoters is observed only in artiodactyls and dogs. It suggests conservation of this maternal gDMR in domestic animals and led us to examine allele-specific gene expression to identify corresponding imprinted genes.

Genomic imprinting can affect all transcript isoforms in an isoform-independent manner, however when there are multiple promoters and they are imprinted in different ways, a mixed pattern of imprinting can occur (isoform-dependent imprinting) [13]. In this study, among previously annotated transcripts in the NCBI RefSeq database, we identified the major *ZNF791* transcript isoforms that were predominantly expressed in pig embryos, as well as in adult dogs, pigs, cattle, sheep, horses, and goats, indicating that *ZNF791* undergoes isoform-independent imprinting. Accordingly, we revealed monoallelic expression of *ZNF791* in these domestic animals, contrasting with the biallelic expression observed in humans, non-human primates, and mice. Additionally, the presence of long non-coding RNAs (lncRNAs) within an imprinted locus can further complicate the imprinting landscape, especially when these lncRNAs overlap with protein-coding mRNA transcripts and run in an antisense orientation. This overlap can lead to allele-specific silencing of mRNAs and ncRNAs through two possible mechanisms: i) transcriptional interference of a promoter and ii) RNA-interference (RNAi) [3]. The former mechanism, transcriptional interference, was demonstrated by the paternal allele-specific silencing of *Igf2r*, which requires the overlap of *Airn* lncRNA transcription over the *Igf2r* promoter, thereby inhibiting RNA polymerase II recruitment [14]. In the current study, we identified a previously unannotated antisense transcript (unannotated AS). The maternally biased expression of this unannotated AS might occur through the latter mechanism, RNAi, mediated by the paternally expressed *ZNF791* transcript. This process occurred without overlapping the promoter region of unannotated AS, indicating a potential RNAi-mediated silencing of the unannotated AS. During oogenesis, RNA transcription, initiating from an upstream transcription start site and extending across intragenic CpG island promoters, is associated with the establishment of DNA methylation imprints at these CGI promoters, which are situated within actively transcribed regions [15]. Accordingly, in our analyses on pigs and cattle, we discovered that the transcription of long *ZNF791* mRNA in oocytes overlaps with intragenic CGI-spanning gDMRs, indicating acquisition of the upstream transcript over the existing CGI that resulted in establishing maternal imprints. Additionally, we identified upstream alternative promoters that are active in porcine and bovine oocytes, supporting the transcription of the long overlapping *ZNF791* transcript. These upstream promoters overlapped with transposable elements (TEs) as observed in the mouse *Impact* locus. It has been suggested that the mechanisms underlying genomic imprinting in therian genomes may have evolved from an epigenetic defense system against integrated TEs [16]. Furthermore, transcripts initiated from species-specific LTRs have been implicated in the establishment of DNA methylation in oocytes [17]. The LTR52, identified in the *ZNF791* locus, may have undergone mutations during evolution but might act as artiodactyl-specific alternative promoters that induce transcription and CGI methylation. Moreover, in mouse oocytes, transcribed regions are enriched with H3K36me3, which mediates *de novo* DNA methylation, thereby leading to the establishment of maternal imprints. Conversely, H3K36me2 is broadly deposited in intergenic regions, mediating moderate levels of DNA methylation [18, 19]. However, in porcine and bovine oocytes, both H3K36me2 and H3K36me3 modifications have been observed to be enriched within transcribed regions and showed a high correlation with DNA methylation [20]. Although these modifications were somewhat more widespread in porcine oocytes than in bovine, their distribution appeared to overlap supporting their conserved role in methylation processes. During post-fertilization, the *ZNF791* DMRs/ICRs might be selectively preserved by being protected against the genome-wide demethylation occurring before embryo implantation.

Over the past three decades, more than 200 imprinted genes have been identified in both mice and humans, with 63 of these genes being common to both species and the majority being species-specific, underscoring both evolutionary conservation and divergence [21]. In domestic pigs, approximately 40 genes have been identified as imprinted, with about 30 of these being common when compared to mice and humans and about a dozen being unique to pigs, according to a previous publication [7] and http://geneimprint.com (accessed Feb 2024). This indicates that the current catalog of porcine imprinted genes may be insufficient to fully understand the complexity and variability of the pig epigenome. Recently, the number of imprinted genes in marsupial mammals was estimated to be 60 [22] suggesting divergence of the imprinting processes across Theria. This divergence can be attributed by molecular factors, as the establishment of the maternal gDMRs for *ZNF791* and *Impact* did not coincide with the onset of genomic imprinting at least 160 MYA. Instead, these gDMRs were established at irregular time points during mammalian evolution. In domestic animals, parent-of-origin effects on body size, body weight, and feed efficiency, which are transmitted either paternally or maternally, have been reported in cross-bred species [7]. According to the parental conflict theory, paternally expressed genes (PEGs) are thought to promote growth [23]. As mentioned, a SNP is located at a putative promoter of the unannotated AS, which may regulate expression of unannotated AS and consequently affect the overlapping *ZNF791-like* expression through RNAi-mediated silencing mechanisms. Because this SNP was associated with residual feed intake (RFI), according to the pig QTLdb, based on a recent GWAS [24] and imprinted genes are dosage-sensitive [25], further research on RFI and genomic imprinting in this locus is warranted. Overall, our findings convey molecular and evolutionary insights into species-specific genomic imprinting in the *ZFN791* locus and may benefit future investigations on genetic selection of livestock.

## Conclusions

In this study, our results reveal that the *ZNF791* locus is imprinted in a lineage-specific manner, shedding light on epigenetic mechanisms uniquely evolved in livestock species. This highlights critical agricultural implications and indicates a deficiency of this imprinting status in humans, non-human primates, and mice.

## Methods

### Animal Ethics and Sample Preparation

All animal procedures were conducted in accordance with guidelines approved by the Institutional Animal Care and Use Committee (IACUC) of the National Institute of Animal Science, Rural Development Administration (RDA) of Korea (NIAS2015-670). The method of parthenogenesis has been described in our previous report [26]. Briefly, ovaries of Landrace × Yorkshire × Duroc (LYD) pigs were obtained from a slaughterhouse. Cumulus-oocyte complexes (COCs) were collected and washed in Tyrode’s lactate-HEPES-polyvinyl alcohol. Oocytes with several layers of cumulus cells were selected and washed three times in TCM-199 based medium (GIBCO, Grand Island, NY, USA). For *in vitro* maturation (IVM), 50 COCs were transferred into 500 µL of maturation medium in four-well dishes and matured for 40 h at 38.5°C in an incubator containing 5% CO_2_.

The cumulus cells were removed, and oocytes having the first polar body were selected and placed in a fusion chamber with 250 µm diameter wire electrodes (BLS, Budapest, Hungary) covered with 0.3 mol/L mannitol solution containing 0.1 mM MgSO_4_, 1.0 mM CaCl_2_, and 0.5 mM HEPES. To achieve fusion, two DC pulses (1 sec interval) of 1.2 kV/cm were applied for 30 µs using an LF101 Electro Cell Fusion Generator (Nepa Gene Co., Ltd., Chiba, Japan). After a 2-h stabilization period, the parthenogenetically activated (PA) embryos were transferred into the oviducts of two LYD surrogate gilts, each aged 12 months on the estrus phase, for further development.

To generate fertilized control (CN) embryos, two LYD gilts were naturally mated with boars twice with a 6-h interval during their natural heat period at the onset of estrus. Both parthenogenetically activated embryos and control embryos were recovered from the euthanized gilts 21 days after the onset of estrus. This timing prevented the abnormal morphological changes in parthenogenetic embryos that typically appear around day 30, while the sizes of parthenogenetic embryos were smaller than that of controls prior to day 30 [27] (Supplementary Fig. 1). The recovery process involved sectioning the reproductive tracts, isolating the placenta from the uterus, and separating embryos from the surrounding placenta, and subsequently the samples were stored in liquid nitrogen until further use.

### Whole-Genome Bisulfite Sequencing and Downstream Analyses

Genomic DNA was isolated from the whole embryos (two biological replicates) and fragmented. The fragmented DNA (200 ng) underwent bisulfite conversion using the EZ DNA Methylation-Gold Kit (Zymo Research, Irvine, CA, USA). To construct the DNA library, 1 ng of DNA was processed with the Accel-NGS Methyl-Seq DNA Library Kit (Swift Biosciences, Inc. Ann Arbor, MI, USA). PCR was conducted with adapter primers and Diastar™ EF-Taq DNA polymerase (Solgent, Daejeon, Korea) with thermal cycling conditions of a 3 m denaturation at 95°C, followed by 10 cycles of 30 s at 95°C, 30 s at 60°C, and 30 s at 72°C, and a final 5 m extension at 72°C. Following bead-based purification, the DNA library was sequenced to produce paired-end reads of 151 nucleotides using the HiSeqX system operated by Macrogen Inc. (Seoul, Korea).

Read quality was evaluated using FastQC (v0.12.1). The raw WGBS reads were trimmed and filtered, using the default parameters of Trim Galore (v0.12.1) to remove adapters and short reads less than 20bp, except for additional trimming to eliminate up to 18 bp of low complexity sequence tags, introduced during the library preparation, from the 3′ end of R1 (–three_prime_clip_R1 18) and from the 5′ end of R2 (– clip_R2 18). The resulting cleaned reads (approximately 420 million pairs for PA1, 425 million pairs for PA2, 431 million pairs for CN1, and 433 million pairs for CN2), were mapped to the pig reference genome (Sscrofa11.1/susScr11) using the Bismark aligner (v0.22.3) with default parameters [28]. Mapping efficiency ranged from 79.6% to 81.3% (Supplementary Table 1). Deduplication was performed with deduplicate_bismark, removing PCR-based duplications that comprised between 13.00% and 15.21% of the mapped pairs. Methylation percentages for cytosines in the CpG context were determined using bismark_methylation_extractor, and CpG sites that were covered at least three reads were further analyzed.

Genomic coordinates of all 2,900 porcine zinc finger transcripts from autosomes (chr1 ∼ chr18) were extracted from the GFF file for the pig reference genome, excluding unplaced scaffolds (chrUn), sex chromosomes (chrX and chrY), and the mitochondrial chromosome. Profile plots and heatmaps for the transcript bodies (TSS ∼ TES) and ± 10 kb regions (–10 kb from TSS and +10 kb from TES) were generated using computeMatrix, plotProfile, and plotHeatmap from deepTools (v3.5.5) [29]. A heatmap for mean DNA methylation levels of all CpG sites within the putative promoter regions (±2 kb from TSS) of selected transcripts was generated using heatmap.2 of the gplots package (v3.1.3.1). For this transcript selection, all 2,900 zinc finger transcripts were initially sorted based on their methylation ratios, defined as the mean methylation level in PA embryos divided by the mean methylation level in CN embryos. From this sorted list, the top 88 transcripts having orthologous loci in humans were identified and re-sorted based on their mean methylation levels in the first PA embryo.

Using the BSmooth method from bsseq (v1.38.0) with default parameters [30], we initially obtained methylation profile curves and candidate DMRs within the putative promoter regions. To avoid possible false positives, we further applied additional procedures using the metilene software (v0.2-8) [31]. For the metilene analysis, we employed default parameters, which include a maximum distance between CpG dinucleotides (-M) of 300, a minimum number of CpGs (-m) of 10, and a mode of operation (-f) of *de novo*. Moreover, we specified a more stringent minimum mean methylation difference (-d) of 0.2 and applied multiple testing correction (-c) with a false discovery rate (FDR) adjustment. We designated regions with an FDR < 0.05 as DMRs. DNA methylation ratios were plotted as histograms using the R package Gviz (v1.44.0) [32].

### Analyses of Hemi-Methylation

For the analysis of WGBS reads overlapping with partially methylated regions, we utilized MethylDackel v.0.6.1 [33] to extract the names and locations of individual reads, count the CpG sites on each read, and calculate the methylation percentage (Supplementary Table 3). We retained only those WGBS reads containing at least three CpGs sites, referred to as ‘qualified reads’, and partially methylated regions supported by more than 30 such qualified reads for further analyses, based on a method previously described [12]. From the percentage of methylated CpGs (mCpGs) for each qualified read, hypomethylated reads (0-20%) and hypermethylated reads (80-100%) were determined. After calculating the percentages of these hypo- and hypermethylated reads among all reads in the partially methylated regions, we plotted the hemi-methylation tendency of these regions, defined as regions with more than or equal to 30% of hypomethylated reads and more than or equal to 30% of hypermethylated reads. For consecutive CpG sites within the partially methylated regions, lollipop plots were drawn using a web-based tool, QUMA (quantification tool for methylation analysis) [34].

### RNA Sequencing and Data Processing

Total RNA was isolated from the whole embryos (two biological replicates) using TRIzol reagent (Sigma-Aldrich, USA). The RNA samples were treated with DNase I to avoid genomic DNA contamination and electrophoresed in 1.2% agarose gels to evaluate RNA integrity, which was confirmed by a 28S/18S rRNA ratio greater than 2 and an RNA integrity number (RIN) greater than 7, using an Agilent 2100 BioAnalyzer. The concentration of RNA was assessed using the ratios of A260/A280 and A260/A230 (1.8–2.0). To select poly-A tails and construct cDNA libraries, 1 µg of total RNA was used with the TruSeq RNA Sample Prep Kit v.2 (Illumina, San Diego, CA, USA). The cDNA libraries were quantified by quantitative Real-Time PCR (qPCR). The Illumina HiSeq2500 RNA-seq platform was used to sequence the library products to produce 101 nucleotide paired-end reads.

Read quality was assessed using FastQC (v0.12.1). To remove adapters and low-quality reads from the raw RNA-seq data, we utilized Trimmomatic (v0.39), applying parameters LEADING:3 TRAILING:3 SLIDINGWINDOW:4:15 MINLEN:36 [35]. This process yielded approximately 27 to 43 million read pairs for the PA and CN embryo samples. Using the STAR aligner (v2.7.9a), the cleaned reads were aligned to the reference genome (Sscrofa11.1/susScr11) with default parameters [36], resulting in 96.0% to 96.5% of the reads being uniquely mapped (Supplementary Table 1). These reads were retrieved with SAMtools [37] and 20.0% to 25.0% of them were subsequently deduplicated using Picard MarkDuplicates for further analyses. BAM files were normalized to generate BigWig files using bamCoverage from deepTools (v3.5.5) [29], with options --binSize 10 and --smoothLength 15. These files were then visualized using Gviz (v1.44.0) [32]. Transcript quantification was performed using Salmon (v1.3.0) in mapping-based mode [38], calculating transcript per million (TPM) values.

### Data Mining and Processing

Our generated data, along with publicly available data downloaded from the NCBI Gene Expression Omnibus (GEO) and other resources, are listed in Supplementary Table 2. WGBS and RNA-seq data were processed using the procedures described earlier in this report, while ChIP-seq, whole genome sequencing (WGS), and whole exome sequencing (WES) data were processed as previously outlined in our report [39]. Reference genomes used in this study include GRCh38.p14/hg38 (human), macFas5 (crab-eating macaque), panTro6 (chimpanzee), rheMac8 (rhesus monkey), Asia_NLE_v1 (gibbon), GRCm39/mm39 (mouse), rat (rn7), equCab3 (horse), canFam3 (dog), susScr11 (pig), bosTau9 (cow), oriAri4 (sheep) and CHIR_1.0 (goat). The VCF files used for SNP analyses are listed in Supplementary Table 2. For VCF files based on a previous genome, the Picard LiftoverVcf (v2.23.8) was used to convert them to match the reference genome mentioned above (Supplementary Table 2).

### LTR Search

When there are multiple *ZNF791* orthologs, we focused on the *ZNF791* gene immediately downstream (3′) of the *MAN2B1* gene across species. We extracted sequences of the promoter regions of the *ZNF791* gene and conducted searches for transposable elements (TEs) in the Dfam database [40] (accessed on December 28, 2023), which utilizes profile hidden Markov models (HMMs), as detailed in Supplementary Fig. 16. For each species analyzed, the search was parameterized with the --species option, while employing the default gathering (GA) threshold via --cut_ga, the Dfam curated threshold, to ensure specificity to repetitive element families. For dogs, however, which did not yield significant matches under the GA threshold, we adapted our approach by utilizing an E-value threshold (-E 0.1) to identify potential LTRs.

### Phylogenetic Analyses

For phylogenetic analyses, LTR-derived sequences from domesticated animal species and the consensus LTR52 sequence (DF000000543.4) from the Dfam database [40] (accessed December 28, 2023) were aligned using MUSCLE for multiple sequence alignment, as implemented in the MEGA11 software [41]. Using MEGA11, the genetic distances were calculated by the Kimura 2-parameter substitution model with uniform rates among sites [42] and maximum-likelihood phylogenetic trees (500 bootstrap replications) were constructed. For another construction of phylogenetic trees with divergence times estimated by TimeTree 5, resulting Newick files were downloaded from http://www.timetree.org (accessed Feb 5, 2024) [43]. The phylogenetic trees were edited using FigTree (v.1.4.4) [44].

### Motif Analyses

*De novo* motif discovery was conducted using the MEME tool from the MEME Suite [45], and the discovered motifs were compared with the JASPAR2022 CORE vertebrates non-redundant database using the TOMTOM tool. Also, motif scanning was performed using the FIMO tool with transcription factor binding sites (TFBSs) data downloaded from the AnimalTFDB v4.0 database [45]. Motif logos were drawn using the R package universalmotif (v1.20.0).

### Statistical Analyses

For DEG analysis, the output files generated by Salmon were imported into R and analyzed using the DESeq2 package (v.1.28.1) [46]. DEGs were identified based on two criteria: an absolute log2-fold change greater than 1 and an FDR below 0.05. To compare mean expression levels at the transcript level between two unrelated groups, specifically PA and CN embryos, the Welch two sample t-test, implemented in R, was used.

## Abbreviations

CN: Control
DEG: Differentially expression gene
DMR: Differentially methylated region
FDR: False discovery rate
ICR: Imprinting control region
LTR: Long terminal repeat
MYA: Million years ago
PA: Parthenogenetically activated
QTLdb: Quantitative trait loci database
TES: Transcription end site
TSS: Transcription start site
WGBS: Whole-genome bisulfite sequencing

## Supplementary Information

Supplementary Figure 1. Measurement of porcine embryos.

Supplementary Figure 2. Positive controls of imprinted expression in pig embryos.

Supplementary Figure 3. Identification of expressed transcripts from RNA-seq data of porcine PA and CN embryos. Supplementary Figure 4. Partial DNA methylation at the *ZNF791* locus in pigs downstream of the *MAN2B1* gene.

Supplementary Figure 5. Partial DNA methylation at the *ZNF791* locus in cattle downstream of the *MAN2B1* gene.

Supplementary Figure 6. Partial DNA methylation at the *ZNF791* locus in sheep downstream of the *MAN2B1* gene.

Supplementary Figure 7. Partial DNA methylation at the *ZNF791* locus in horses downstream of the *MAN2B1* gene.

Supplementary Figure 8. Partial DNA methylation at the *ZNF791* locus in goat downstream of the *MAN2B1* gene.

Supplementary Figure 9. Partial DNA methylation at the *ZNF791* locus in dogs downstream of the *MAN2B1* gene.

Supplementary Figure 10. DNA methylation at the *ZNF791* locus in pig embryos downstream of the *MAN2B1* gene.

Supplementary Figure 11. Human lung exome from WES and RNA-seq.

Supplementary Figure 12. Expressed *ZNF791* transcripts in humans, primates, and mice.

Supplementary Figure 13. Expressed *ZNF791* transcripts in dogs, cattle, and pigs.

Supplementary Figure 14. Expressed transcripts at the *ZNF791* locus in pig oocytes.

Supplementary Figure 15. An expressed unannotated transcript at the *ZNF791* locus in pig oocytes.

Supplementary Figure 16. Dfam database search process.

Supplementary Figure 17. Expressed transcripts at the *ZNF791* locus in cow oocytes.

Supplementary Figure 18. LTR-initiated transcription and establishment of methylation imprint in mouse oocytes. Supplementary Figure 19. Expressed *ZNF791* transcripts in sheep, horses, and goats.

Supplementary Figure 20. Multiple sequence alignment of LTRs. Supplementary Figure 21. Motif discovery and comparison.

Supplementary Figure 22. QTL analysis for the upstream of the unannotated antisense transcript.

Supplementary Table 1. Summary of data metrics.

Supplementary Table 2. Identification of DMR-associated transcripts. Supplementary Table 3. Deposited data.

Supplementary Table 4. Analyses of partially methylated regions.

Supplementary Table 5. Individual-matched analyses of genomic DNA and RNA-seq read counts on informative SNPs.

## Declarations

### Ethics approval and consent to participate

All animal procedures were conducted in accordance with guidelines approved by the Institutional Animal Care and Use Committee (IACUC) of the National Institute of Animal Science, Rural Development Administration (RDA) of Korea (NIAS2015-670).

### Consent for publication

Not applicable

### Availability of data and materials

WGBS and RNA-seq data generated in this study have been submitted to the NCBI Gene Expression Omnibus (GEO; https://www.ncbi.nlm.nih.gov/geo/) under accession number GSE263495 and GSE263494, respectively. The publicly available datasets and VCF files used in this study are listed in Supplementary Table 3. Our published sheep RNA-seq data are available under accession number GSE253249.

## Authors’ Contributions

Conceptualization, KL, SH, and JA; methodology, JA, ISH, and MRP; processing sequencing data, JA; validation, JA and KL; formal analysis, JA; investigation, JA, ISH, MRP, MRV, and AER; resources, ISH, ICC, and AER; writing—original draft preparation, JA and KL; writing—review and editing, JA, SH, and KL; visualization, JA; supervision, SH and KL; funding acquisition, SH, and KL

## Acknowledgements

We are thankful to Ms. Michelle Milligan for proofreading this manuscript.

