## Supplementary Figure 1-15 and 17-22 for "Lineage-specific genomic imprinting in the *ZNF791* locus"

**Supplementary Figure 1. Measurement of porcine embryos.**
**Supplementary Figure 2. Positive controls of imprinted expression in pig embryos.**
**Supplementary Figure 3. Identification of expressed transcripts from RNA-seq data of porcine PA and CN embryos.**
**Supplementary Figure 4. Partial DNA methylation at the *ZNF791* locus in pigs downstream of the *MAN2B1* gene.**
**Supplementary Figure 5. Partial DNA methylation at the *ZNF791* locus in cattle downstream of the *MAN2B1* gene.**
**Supplementary Figure 6. Partial DNA methylation at the *ZNF791* locus in sheep downstream of the *MAN2B1* gene.**
**Supplementary Figure 7. Partial DNA methylation at the *ZNF791* locus in horses downstream of the *MAN2B1* gene.**
**Supplementary Figure 8. Partial DNA methylation at the *ZNF791* locus in goat downstream of the *MAN2B1* gene.**
**Supplementary Figure 9. Partial DNA methylation at the *ZNF791* locus in dogs downstream of the *MAN2B1* gene.**
**Supplementary Figure 10. DNA methylation at the *ZNF791* locus in pig embryos downstream of the *MAN2B1* gene.**
**Supplementary Figure 11. Human lung exome from WES and RNA-seq.**
**Supplementary Figure 12. Expressed *ZNF791* transcripts in humans, primates, and mice.**
**Supplementary Figure 13. Expressed *ZNF791* transcripts in dogs, cattle, and pigs.**
**Supplementary Figure 14. Expressed transcripts at the *ZNF791* locus in pig oocytes.**
**Supplementary Figure 15. An expressed unannotated transcript at the *ZNF791* locus in pig oocytes.**
**Supplementary Figure 16. Dfam database search process.**
**Supplementary Figure 17. Expressed transcripts at the *ZNF791* locus in cow oocytes.**
**Supplementary Figure 18. LTR-initiated transcription and establishment of methylation imprint in mouse oocytes.**
**Supplementary Figure 19. Expressed *ZNF791* transcripts in sheep, horses, and goats.**
**Supplementary Figure 20. Multiple sequence alignment of LTRs.**
**Supplementary Figure 21. Motif discovery and comparison.**
**Supplementary Figure 22. QTL analysis for the upstream of the unannotated antisense transcript.**

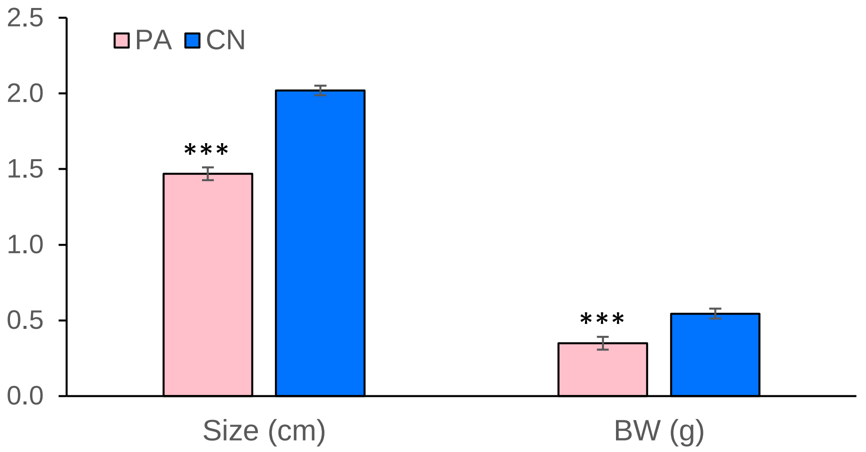


**Supplementary Figure 1. Measurement of porcine embryos.** Size, body weight (BW), and morphology of the control (CN, n = 10) and parthenote (PA, n = 10). Data are presented as mean ± SEM.***, $P < 0.001$.

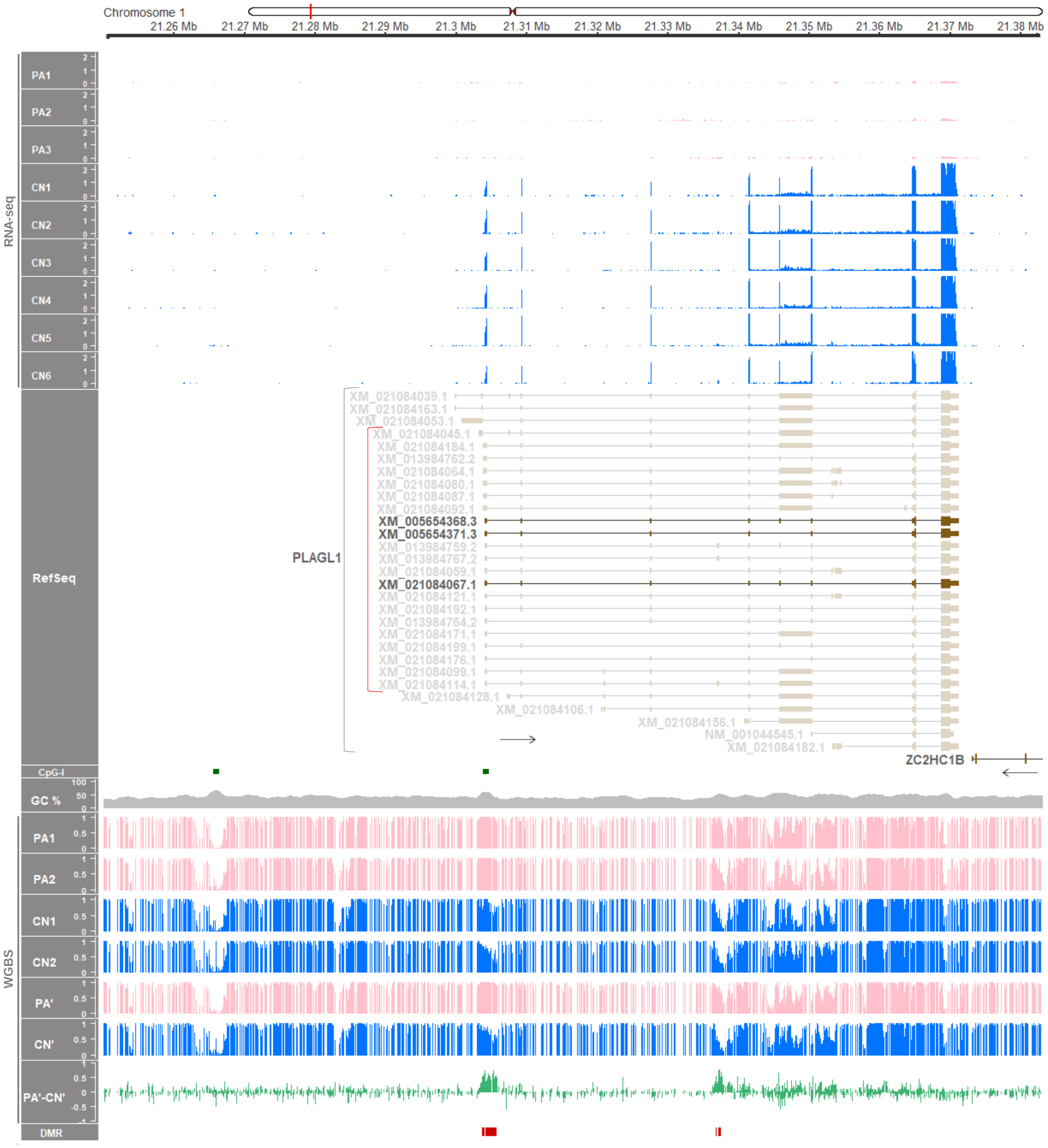


**Supplementary Figure 2. Positive controls of imprinted expression in pig embryos. (A)** The known imprinted gene, PLAG1 like zinc finger 1 (*PLAGL1*), was expressed exclusively in CN embryos indicating its paternal expression. Maternally methylated DMRs were identified within the transcribed region. The transcript IDs are from the NCBI RefSeq annotation. RNA-seq read coverages and WGBS methylation levels are presented as TPM values and DNA methylation ratios, respectively. The 21 transcripts associated with the DMR are marked with a red bracket. Non-expressed *PLAGL1* transcripts are covered with white shading for visual distinction.

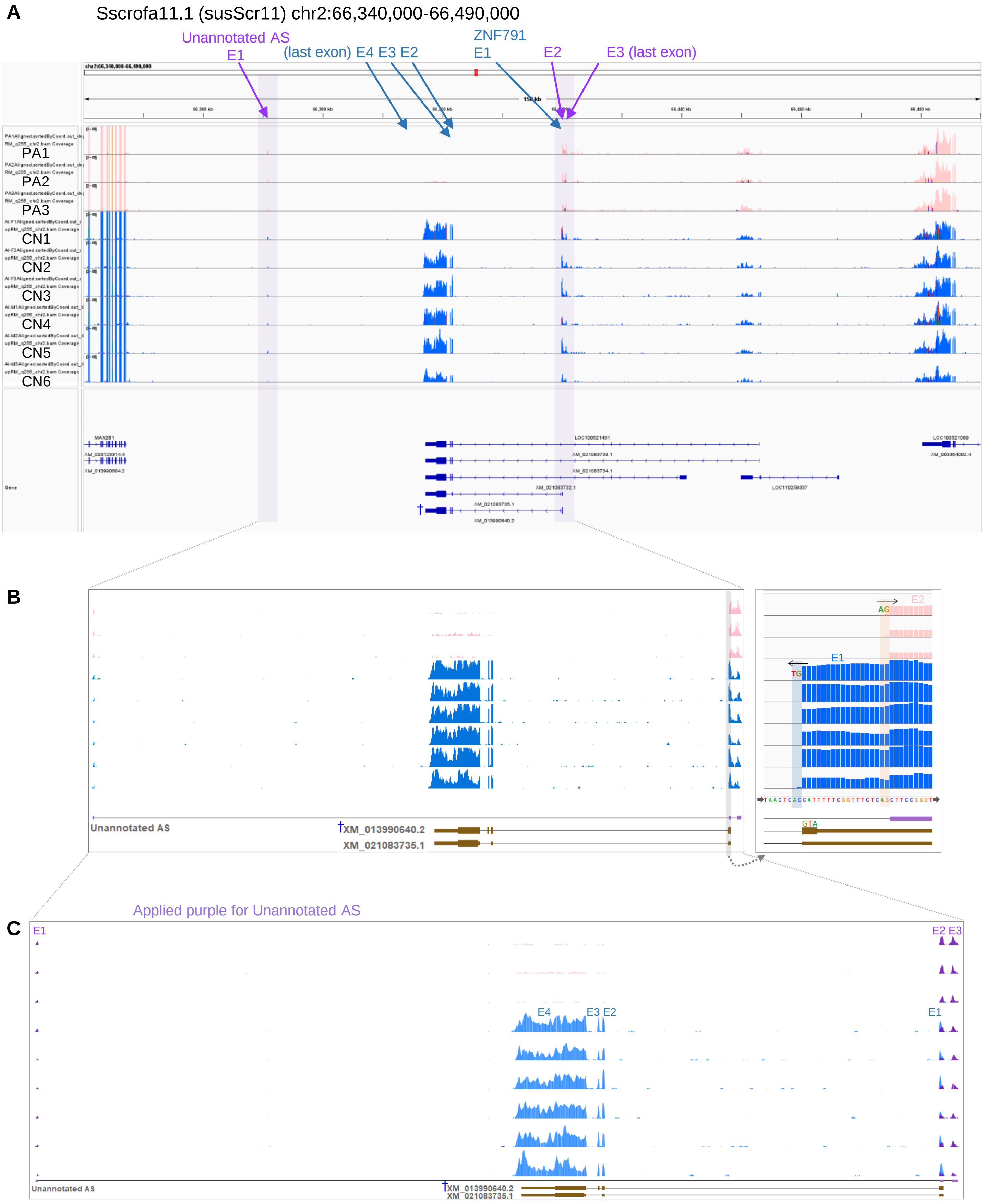


**Supplementary Figure 3. Identification of expressed transcripts from RNA-seq data of porcine PA and CN embryos. (A)** The pig *ZNF791-like* locus is shown with Integrative Genomics Viewer (IGV). **(B)** Close views of expressed transcripts. Splicing donor (GT) and acceptor (AG) are denoted with directional arrows. The start codon (ATG) is marked. **(C)** Visualization of expression and overlapping of the *ZNF791-like* transcript (blue) and Unannotated AS (purple). A predominant *ZNF791-like* transcript is marked with †.

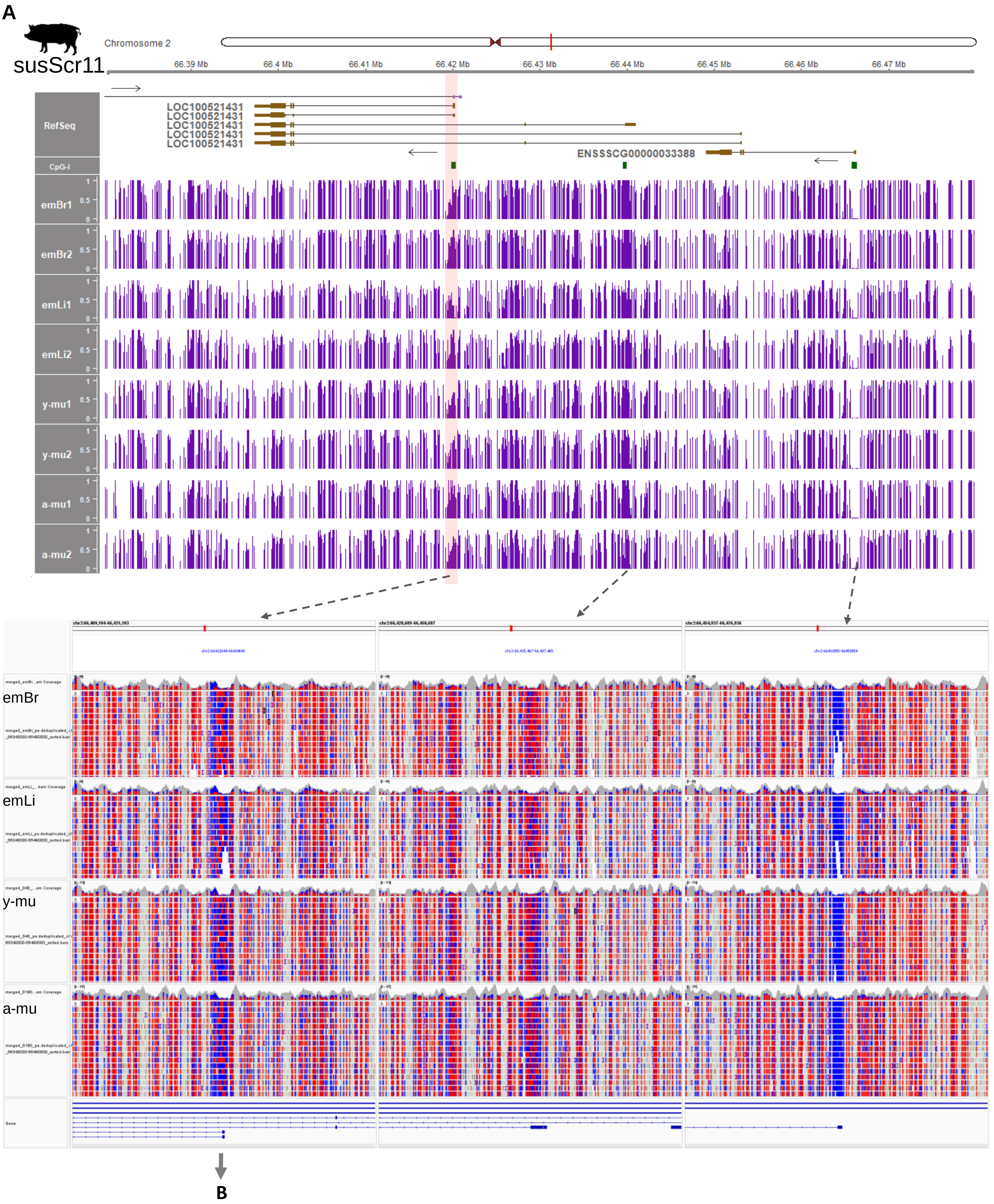
A
susScr11
Chromosome 2
66.39 Mb
66.4 Mb
66.41 Mb
66.42 Mb
66.43 Mb
66.44 Mb
66.45 Mb
66.46 Mb
66.47 Mb
RefSeq
LOC100521431
LOC100521431
LOC100521431
LOC100521431
LOC100521431
ENSSSCG00000033388
CpG-I
emBr1
emBr2
emLi1
emLi2
y-mu1
y-mu2
a-mu1
a-mu2
emBr
emLi
y-mu
a-mu
B

B

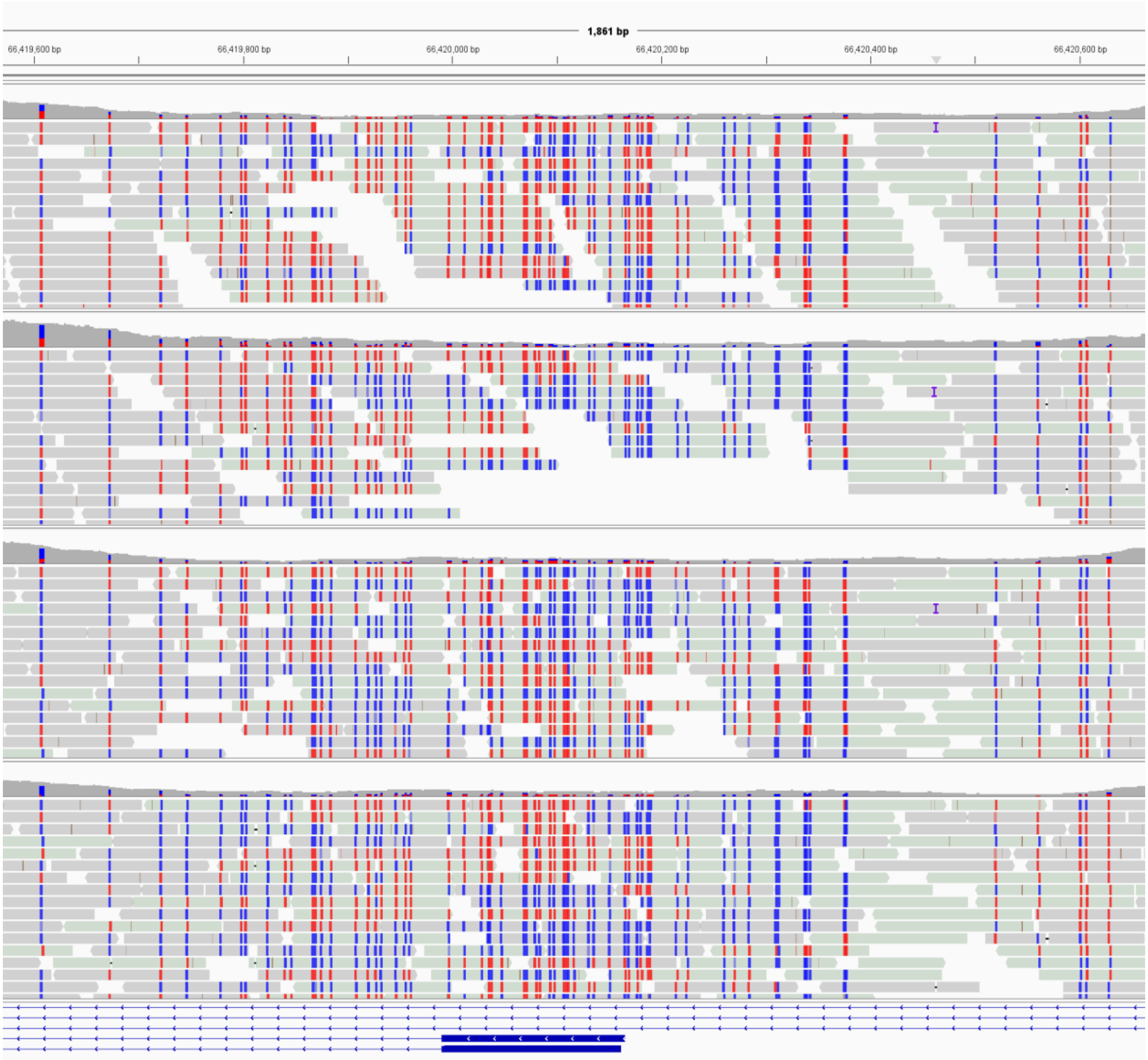


**Supplementary Figure 4. Partial DNA methylation at the *ZNF791* locus in pigs downstream of the *MAN2B1* gene. (A)** In the split-screen view of merged reads displayed at the bottom, the red color represents unconverted (methylated) cytosines, and the blue color represents bisulfite-converted (unmethylated) cytosines. The CpG sites are displayed in either red or blue. **(B)** A close view of the partially methylated region indicated in A around chr2:66,420,000 (66.42 Mb), which is highlighted with a red bar. This region was analyzed for hemi-methylation tendency and status as described in Figures 3C and 3D. The same approach was applied for other hemi-methylation analyses in the following Supplementary Figures 5~10.

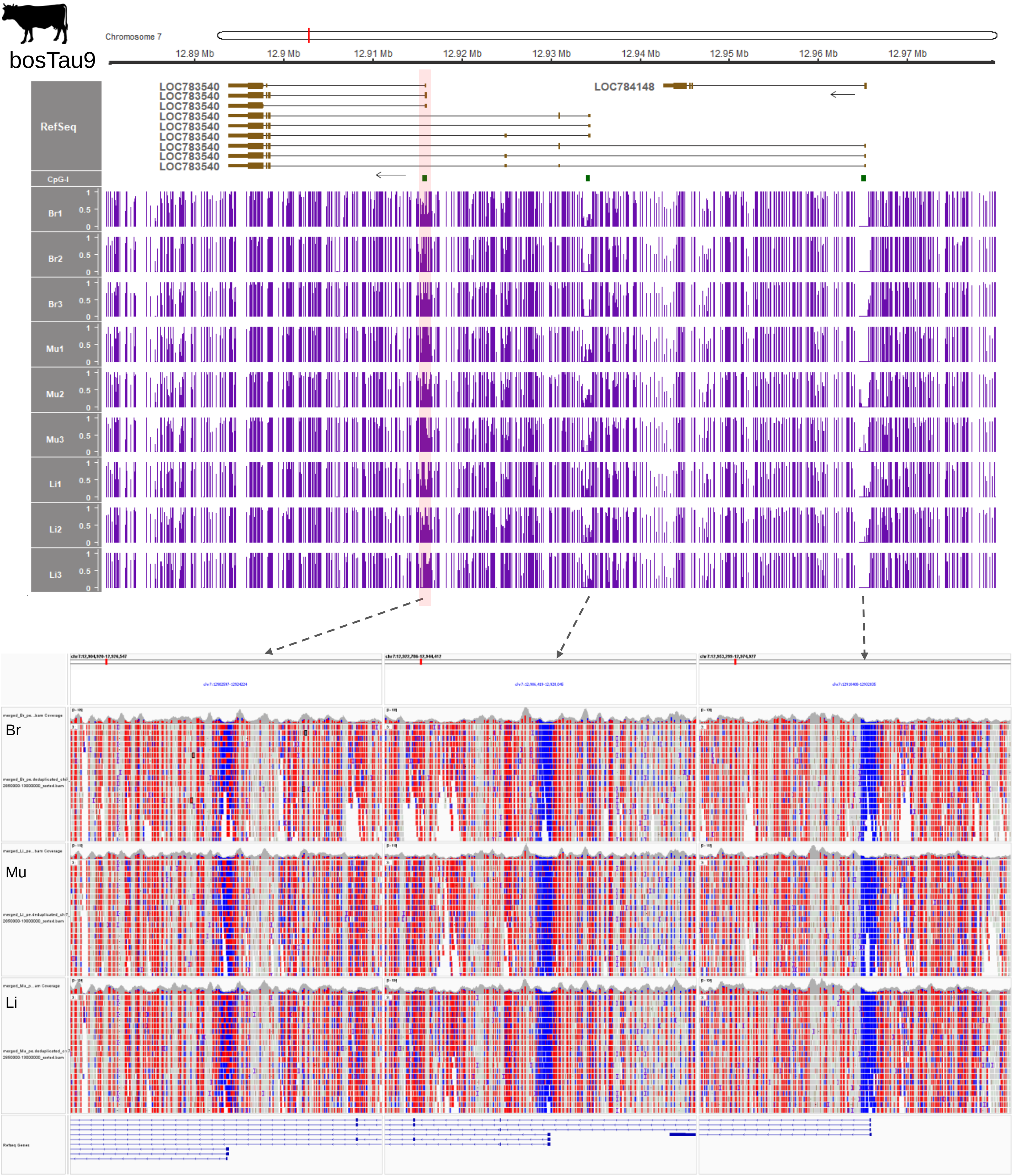


**Supplementary Figure 5. Partial DNA methylation at the *ZNF791* locus in cattle downstream of the *MAN2B1* gene.** In the split-screen view displayed at the bottom, the red color represents unconverted (methylated) cytosines, and the blue color represents bisulfite-converted (unmethylated) cytosines. The CpG sites are displayed in either red or blue.

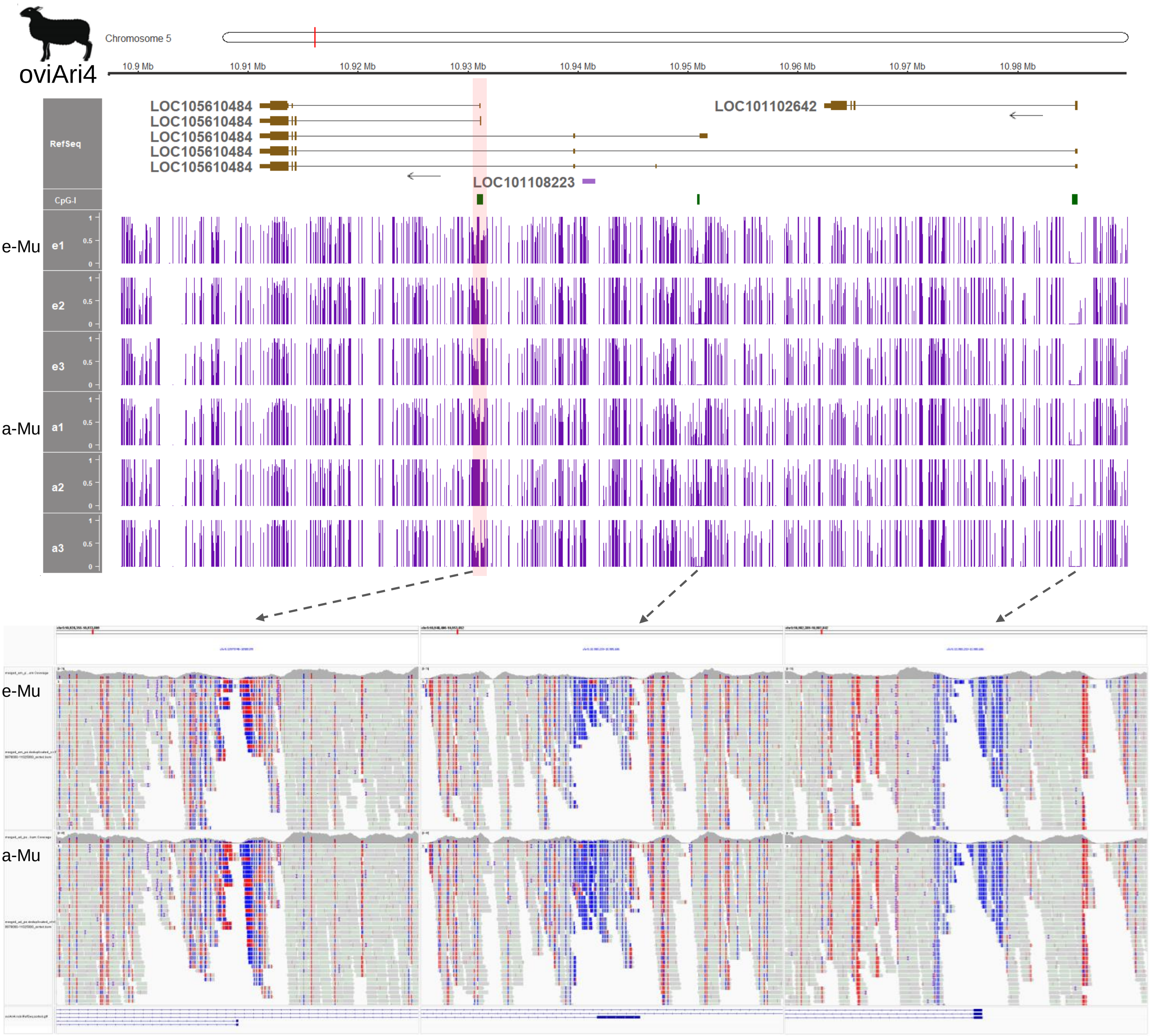


**Supplementary Figure 6. Partial DNA methylation at the *ZNF791* locus in sheep downstream of the *MAN2B1* gene.** Split screen view of merged reads are displayed at the bottom where red represents unconverted (methylated) and blue represents bisulfite-converted (unmethylated) cytosines. The CpG sites are displayed in either red or blue. e-Mu, embryonic muscle; a-Mu, adult muscle.

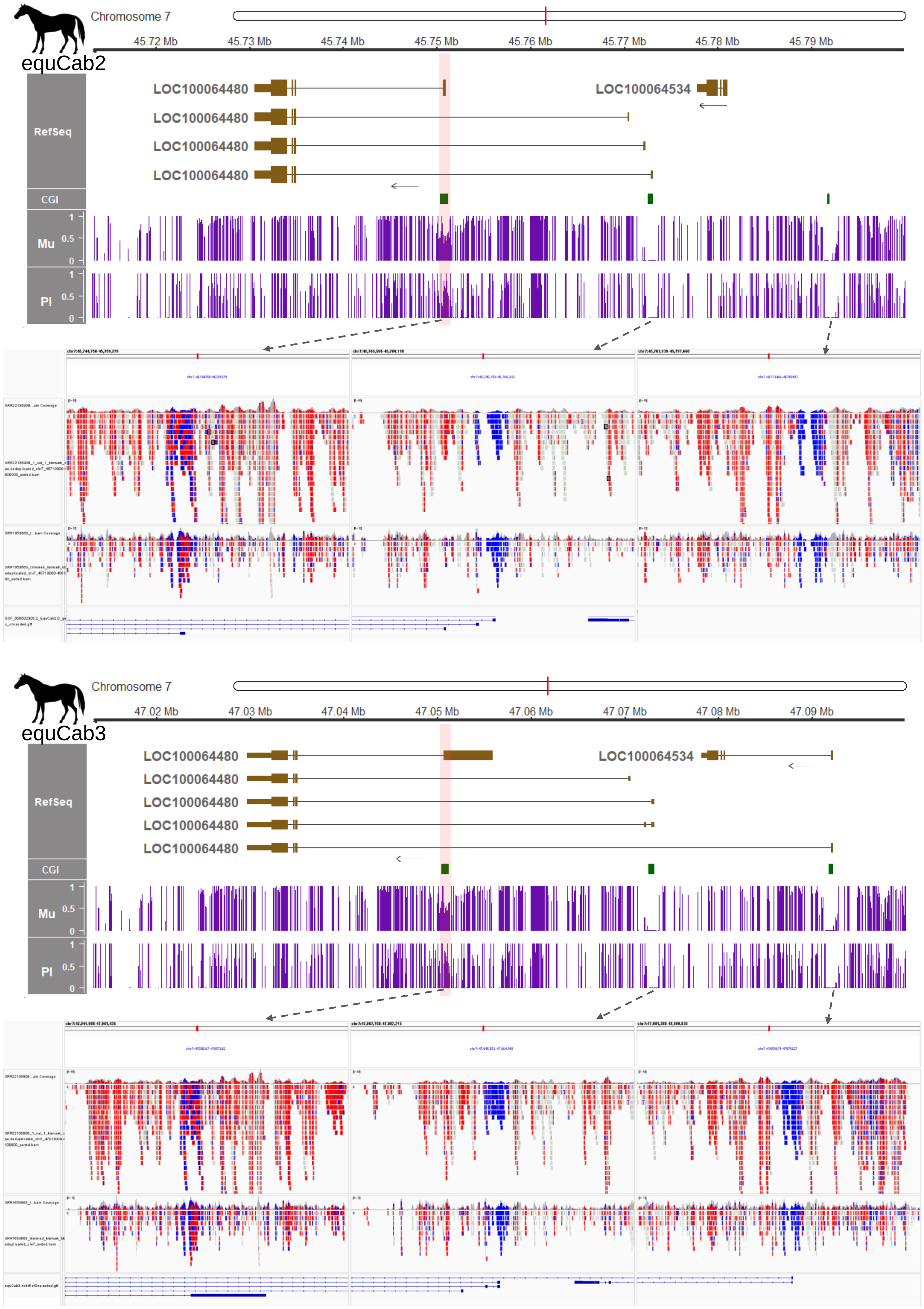
equCab2
Chromosome 7
45.72 Mb
45.73 Mb
45.74 Mb
45.75 Mb
45.76 Mb
45.77 Mb
45.78 Mb
45.79 Mb
RefSeq
LOC100064480
LOC100064480
LOC100064480
LOC100064480
LOC100064534
CGI
Mu
PI
1
0.5
0
equCab3
Chromosome 7
47.02 Mb
47.03 Mb
47.04 Mb
47.05 Mb
47.06 Mb
47.07 Mb
47.08 Mb
47.09 Mb
RefSeq
LOC100064480
LOC100064480
LOC100064480
LOC100064480
LOC100064480
LOC100064534
CGI
Mu
PI
1
0.5
0

**Supplementary Figure 7. Partial DNA methylation at the *ZNF791* locus in horses downstream of the *MAN2B1* gene.** Split screen view of merged reads are displayed at the bottom where red represents unconverted (methylated) and blue represents bisulfite-converted (unmethylated) cytosine. The CpG sites are displayed in either red or blue. The same data from horse skeletal muscle used in Fig. 3 were aligned to both the current reference genome (EquCab3.0/equCab3) and the previous (EquCab2.0/equCab2) genome.

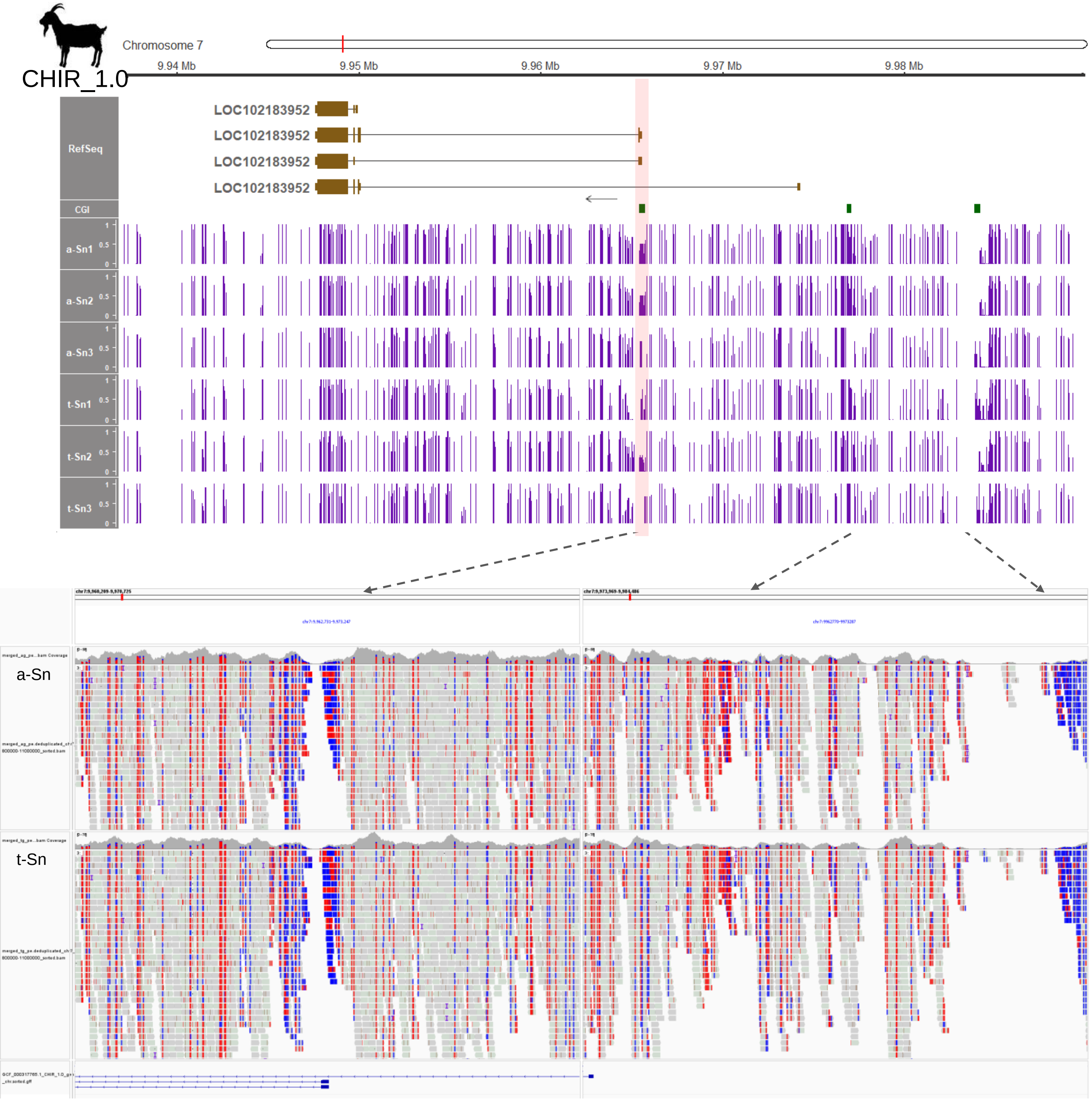


**Supplementary Figure 8. Partial DNA methylation at the *ZNF791* locus in goat downstream of the *MAN2B1* gene.** Split screen view of merged reads are displayed at the bottom (red represents unconverted (methylated) and blue represents bisulfite-converted (unmethylated). The CpG sites are displayed in either red or blue.

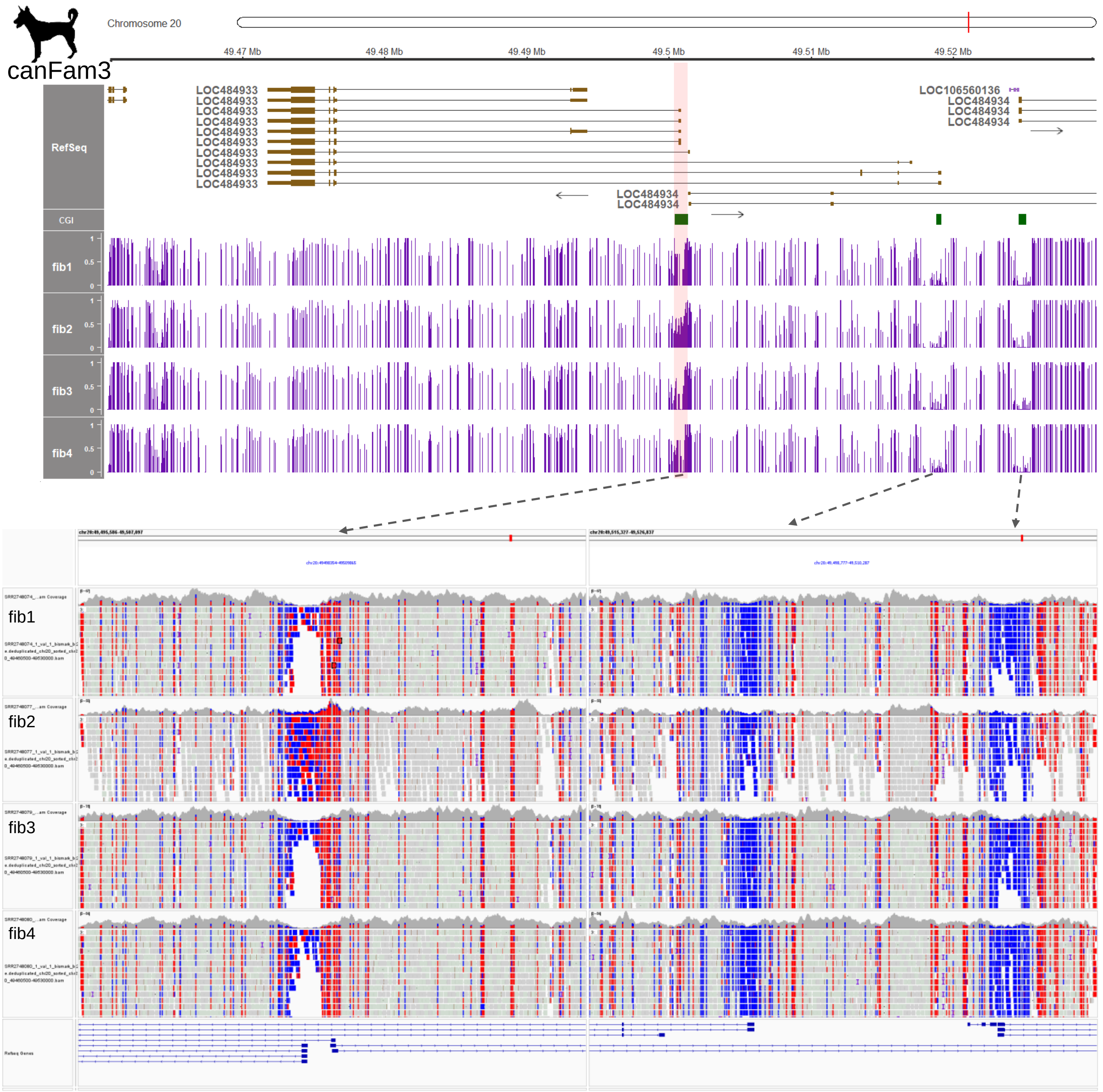


**Supplementary Figure 9. Partial DNA methylation at the *ZNF791* locus in dogs downstream of the *MAN2B1* gene.** Split screen view of sample reads are displayed, where red represents unconverted (methylated) and blue represents bisulfite-converted (unmethylated) cytosines. The CpG sites are displayed in either red or blue.

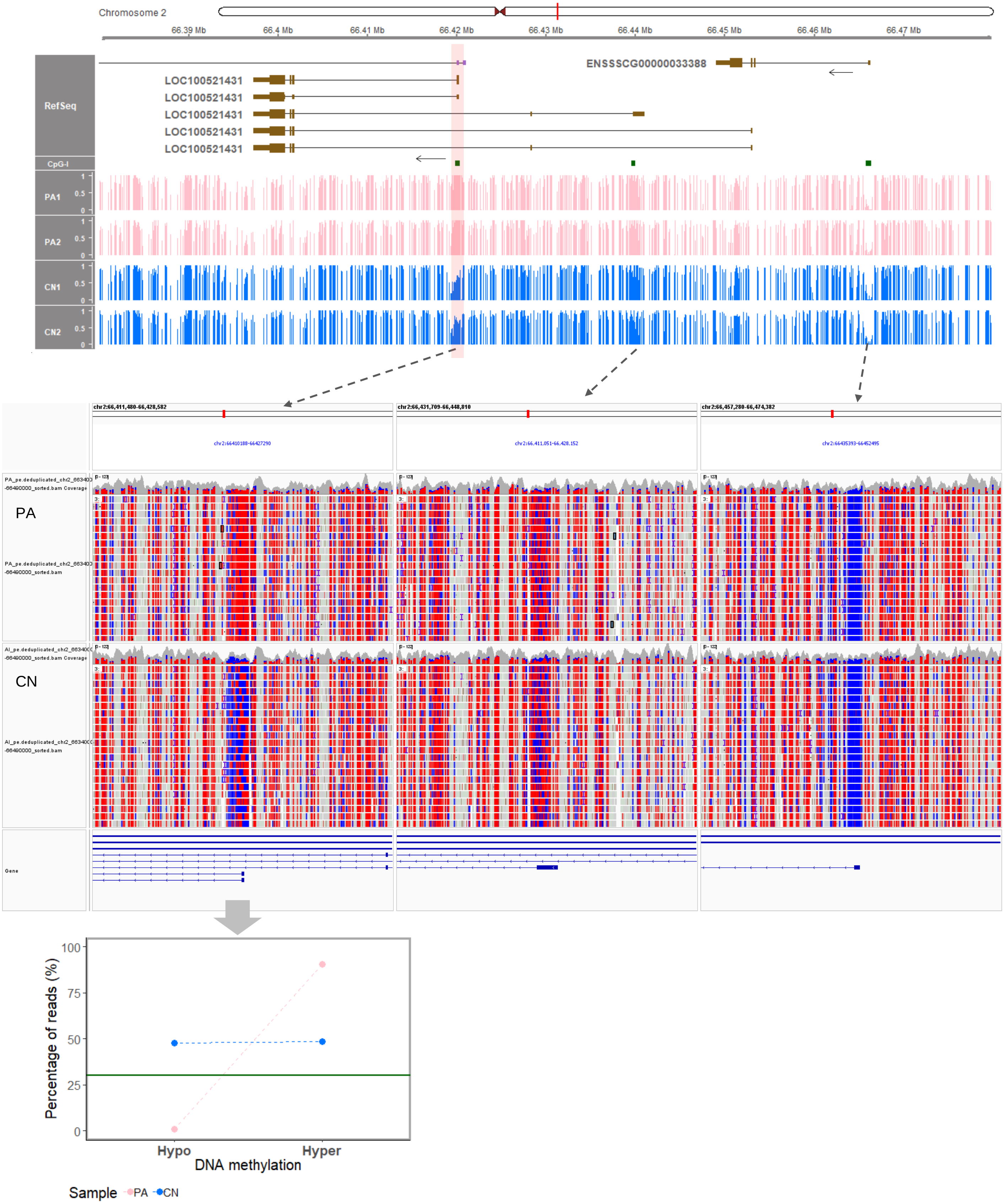


**Supplementary Figure 10. DNA methylation at the *ZNF791* locus in pig embryos downstream of the *MAN2B1* gene.** In the split-screen view of merged reads displayed in the middle, the red color represents unconverted (methylated) cytosines, and the blue color represents bisulfite-converted (unmethylated) cytosines. The CpG sites are displayed in either red or blue. At the bottom, the full methylated region in PA embryos exhibited hypermethylation only, while the partially methylated in CN embryos showed hemi-methylation tendency.

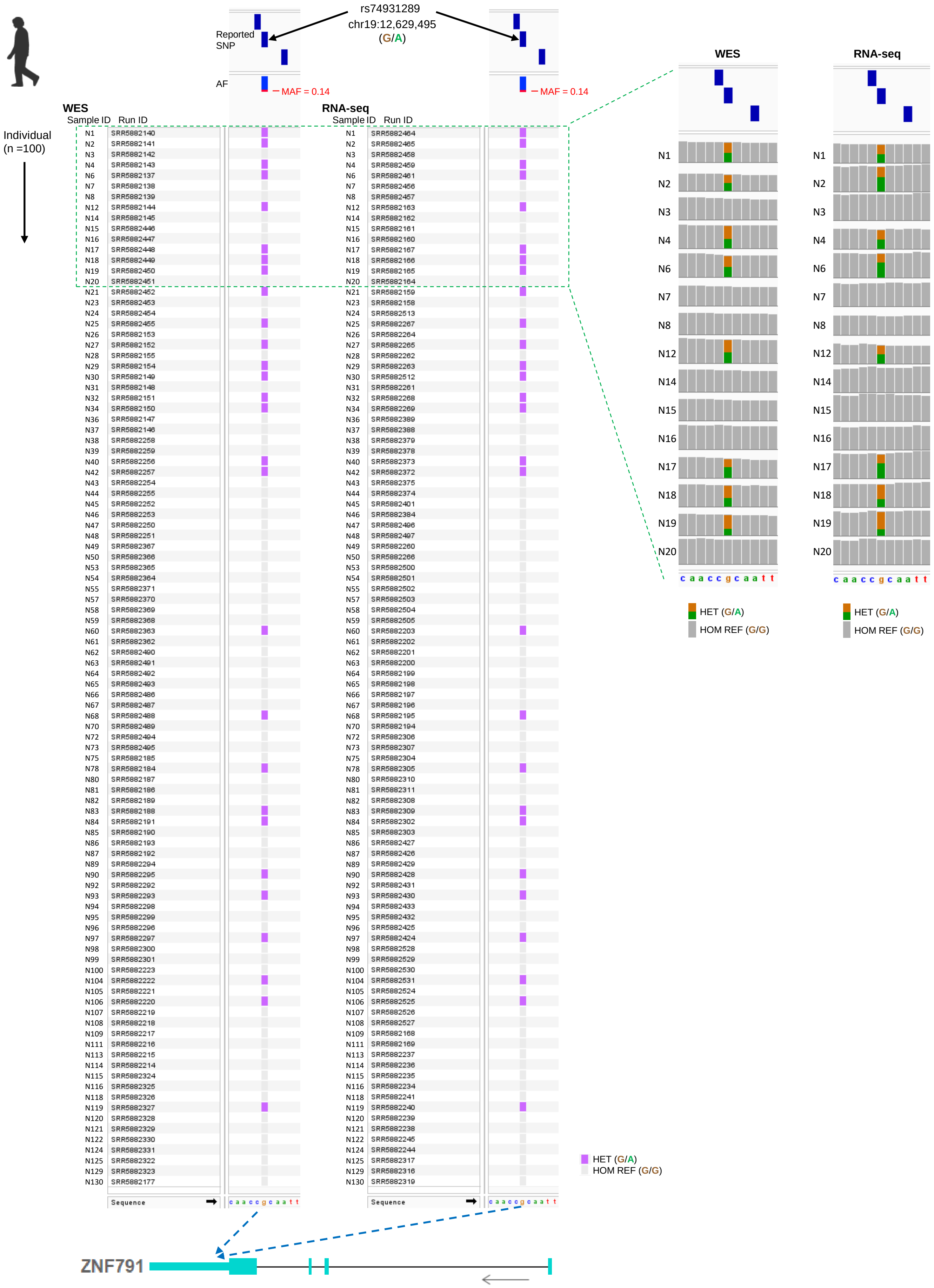


**Supplementary Figure 11. Human lung exome from WES and RNA-seq. (A)** SNP information for 100 individuals is shown using a VCF file. A SNP at the last exon (3′ UTR) of *ZNF791* [chr19:12,629,495; G/A (reference/alternate); rs74931289] was targeted. A minor allele frequency of this targeted SNP was 0.139 (MAF > 0.05). An informative, heterozygous SNPs (G/A) are indicated with blue. Grey indicates homozygous reference allele (G/G). The inset shows a close view of each SNP in lung tissues of each individual.

A

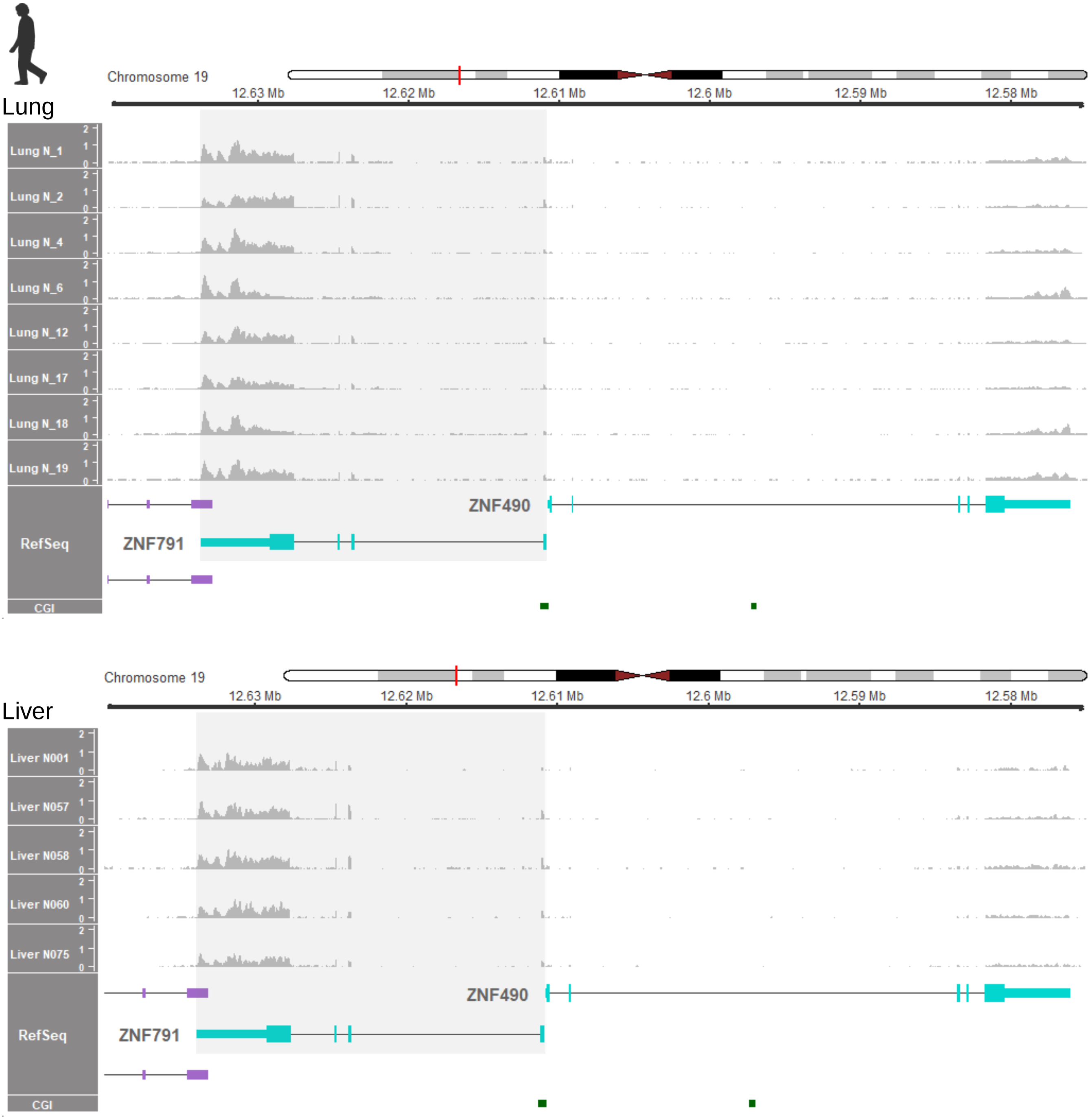
Lung
Chromosome 19
12.63 Mb
12.62 Mb
12.61 Mb
12.6 Mb
12.59 Mb
12.58 Mb
Lung N_1
Lung N_2
Lung N_4
Lung N_6
Lung N_12
Lung N_17
Lung N_18
Lung N_19
RefSeq
ZNF490
ZNF791
CGI
Liver
Chromosome 19
12.63 Mb
12.62 Mb
12.61 Mb
12.6 Mb
12.59 Mb
12.58 Mb
Liver N001
Liver N057
Liver N058
Liver N060
Liver N075
RefSeq
ZNF490
ZNF791
CGI

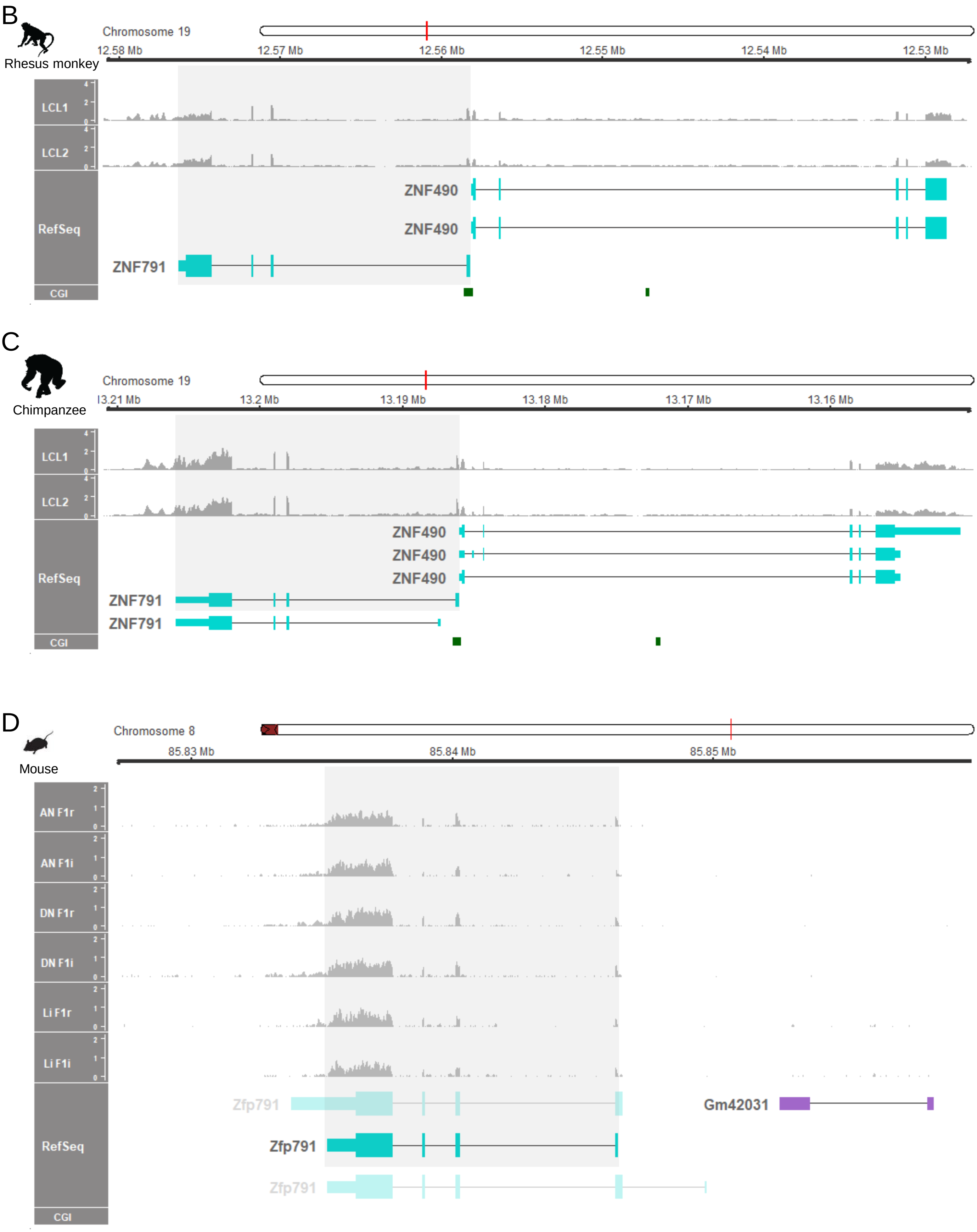


**Supplementary Figure 12. Expressed *ZNF791* transcripts in humans, primates, and mice. (A)** Human *ZNF791* mRNA expression. **(B, C)** Primate *ZNF791* mRNA expression. **(D)** Mouse *Zfp791* mRNA expression patterns. Transcripts that are expressed are indicated with grey highlights. RNA-seq read coverages were normalized to TPM. Datasets used in Figure 4 (PRJNA395106 for human lung, hum0158.v2 for human liver, PRJNA563344 for rhesus monkey and chimpanzee, and GSE70484 for mouse) are analyzed.

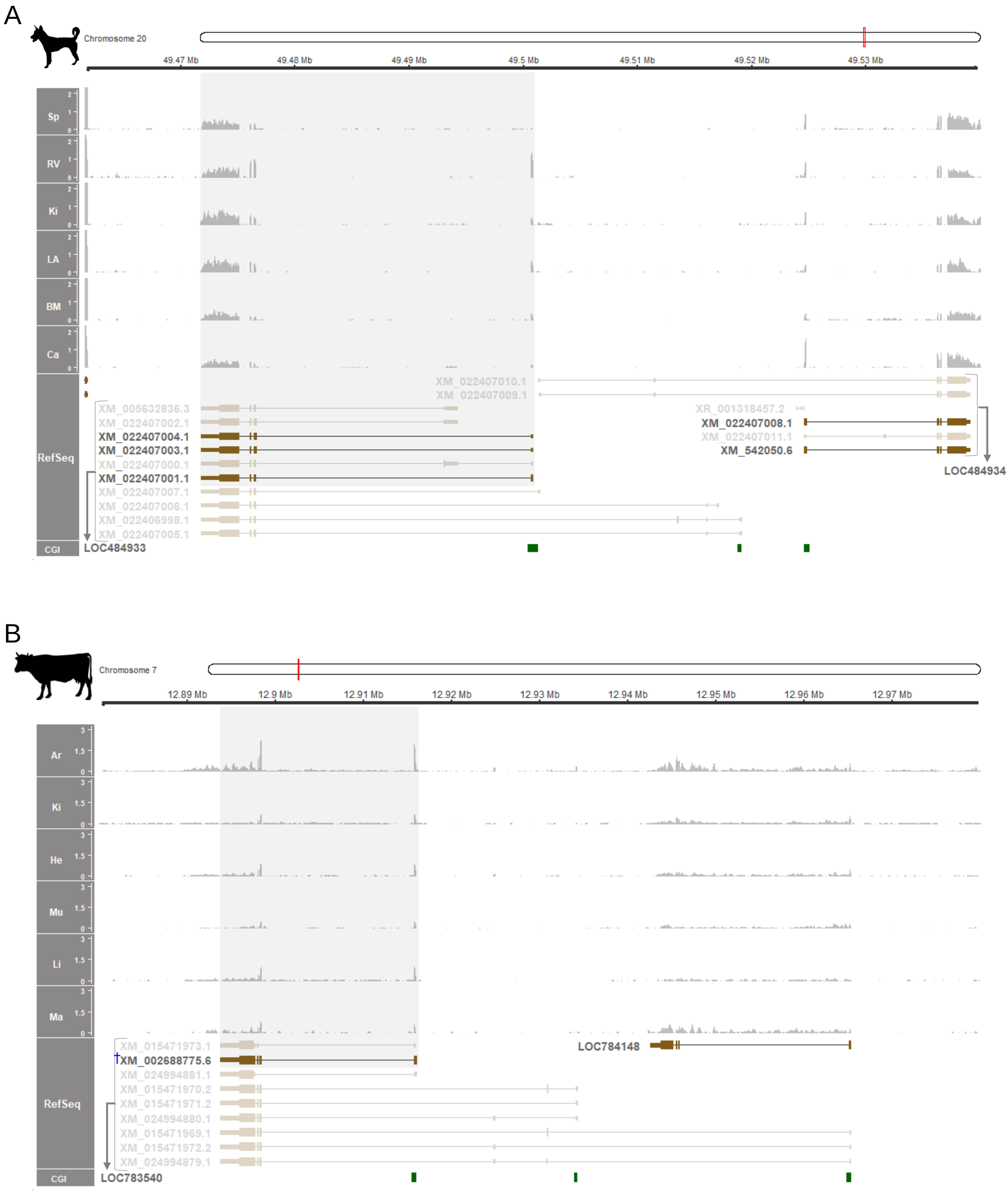
A
Chromosome 20
49.47 Mb
49.48 Mb
49.49 Mb
49.5 Mb
49.51 Mb
49.52 Mb
49.53 Mb
Sp
RV
Ki
LA
BM
Ca
RefSeq
CGI
XM_022407010.1
XM_022407009.1
XM_005632836.3
XM_022407002.1
XM_022407004.1
XM_022407003.1
XM_022407000.1
XM_022407001.1
XM_022407007.1
XM_022407006.1
XM_022406998.1
XM_022407005.1
LOC484933
XR_001318457.2
XM_022407008.1
XM_022407011.1
XM_542050.6
LOC484934
B
Chromosome 7
12.89 Mb
12.9 Mb
12.91 Mb
12.92 Mb
12.93 Mb
12.94 Mb
12.95 Mb
12.96 Mb
12.97 Mb
Ar
Ki
He
Mu
Li
Ma
RefSeq
CGI
XM_015471973.1
XM_002688775.6
XM_024994881.1
XM_015471970.2
XM_015471971.2
XM_024994880.1
XM_015471969.1
XM_015471972.2
XM_024994879.1
LOC783540
LOC784148

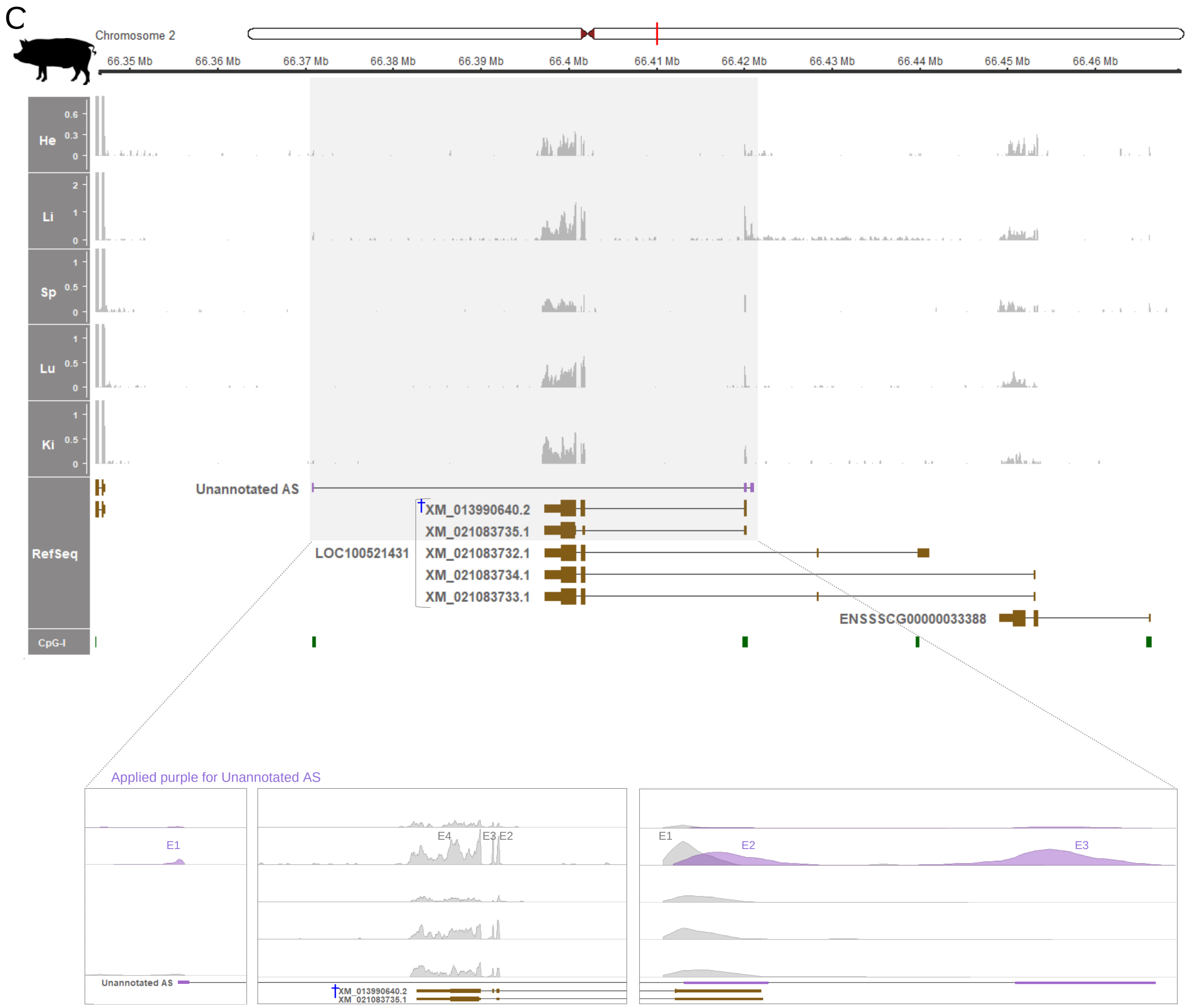


**Supplementary Figure 13. Expressed *ZNF791* transcripts in dogs, cattle, and pigs. (A)** Dog *ZNF791* mRNA expression. **(B)** Cattle *ZNF791* mRNA expression. **(C)** Pig *ZNF791* mRNA expression patterns. Transcripts that are expressed are indicated with grey highlights. RNA-seq read coverages were normalized to TPM. Datasets used in Figure 4 (PRJNA396033 for dog, ERP118133 for cow, and PRJNA309108 for pig) are analyzed. The major predominant *ZNF791* transcript is marked with a blue symbol (†).

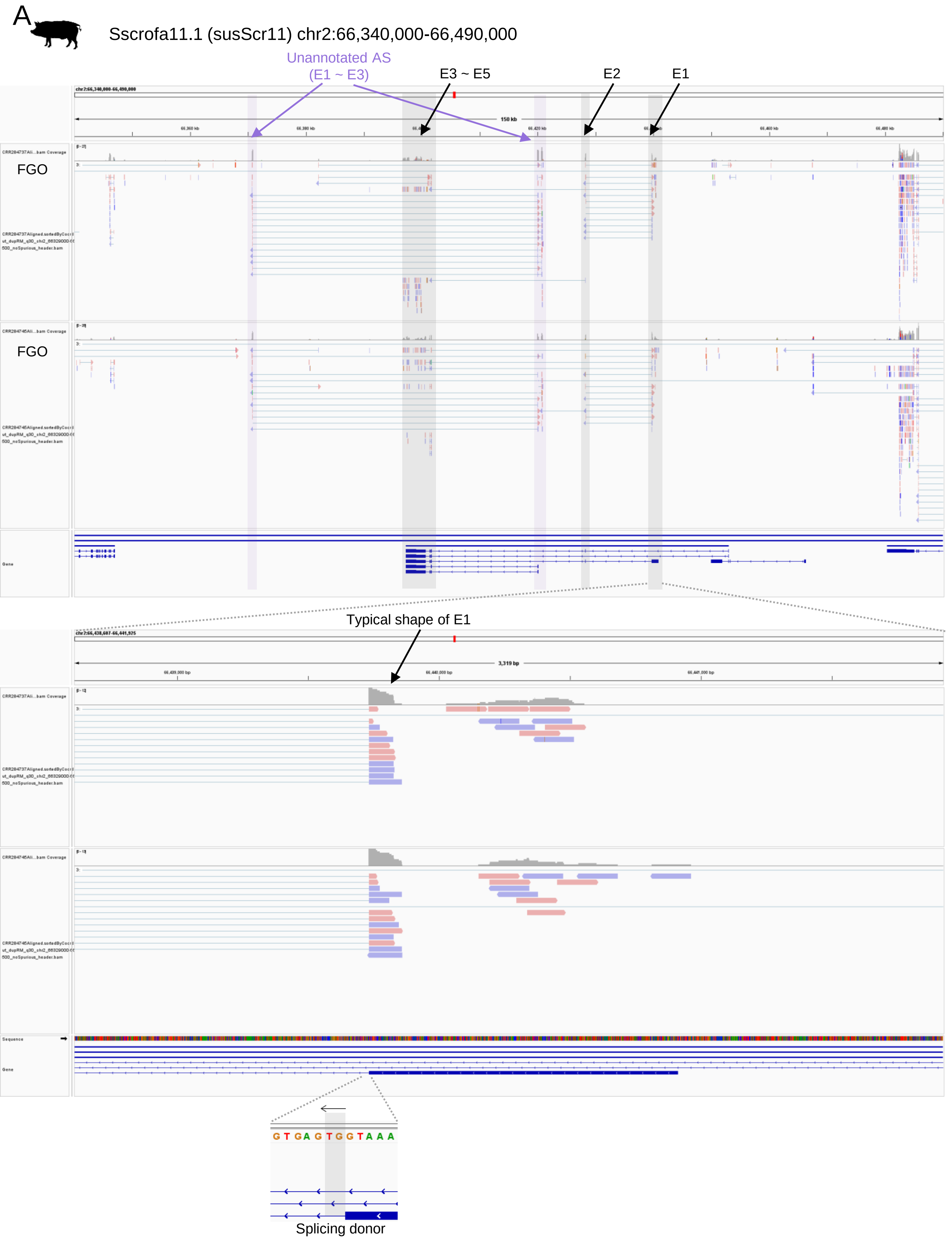


**Supplementary Figure 14. Expressed transcripts at the *ZNF791* locus in pig oocytes. (A)** The 1st, 2nd and last exons of expressed *ZNF791* transcript in pig oocytes are indicated with grey highlights. Zoomed view of the 1st exon region. The splicing donor (GT) is shown. The exon1 could be shorter than the annotated one because the typical shape of exon1 is shorter.

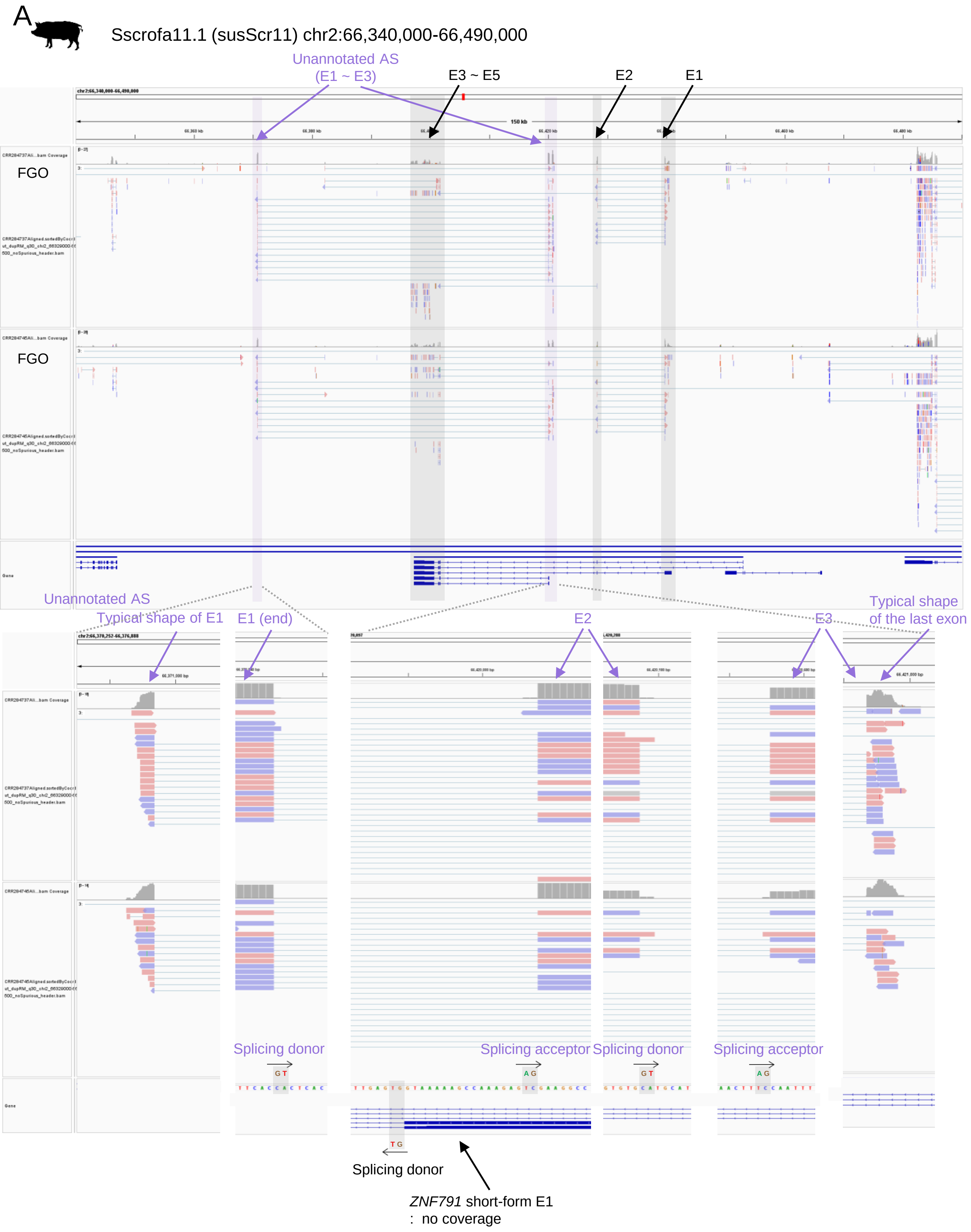


**Supplementary Figure 15. An expressed unannotated transcript at the *ZNF791* locus in pig oocytes. (A)** The 1st, 2nd and last exons of expressed *ZNF791* transcript in pig oocytes are indicated with grey highlights. Zoomed view of the 1st exon region. The splicing donor (GT) is shown. The exon1 could be shorter than the annotated one because of the typical shape of exon1 is shorter.

**Supplementary Figure 16. Dfam database search process.** The Dfam database was queried with input sequence from each species. These sequences were searched for transposable elements (TEs) specific to each organism using profile hidden Markov models (HMMs). This process was carried out in the following order:

1. Dfam database (https://www.dfam.org/home)
2. Pig (*Sus scrofa*) sequence
3. Cattle (*Bos taurus*) sequence
4. Sheep (*Ovis aries*) sequence
5. Horse (*Equus caballus*) sequence
6. Goat (*Capra hircus*) sequence
7. Dog (*Canis lupus familiaris*) sequence

*Data for Supplementary Figure 16 are presented in a supplementary Word file.*

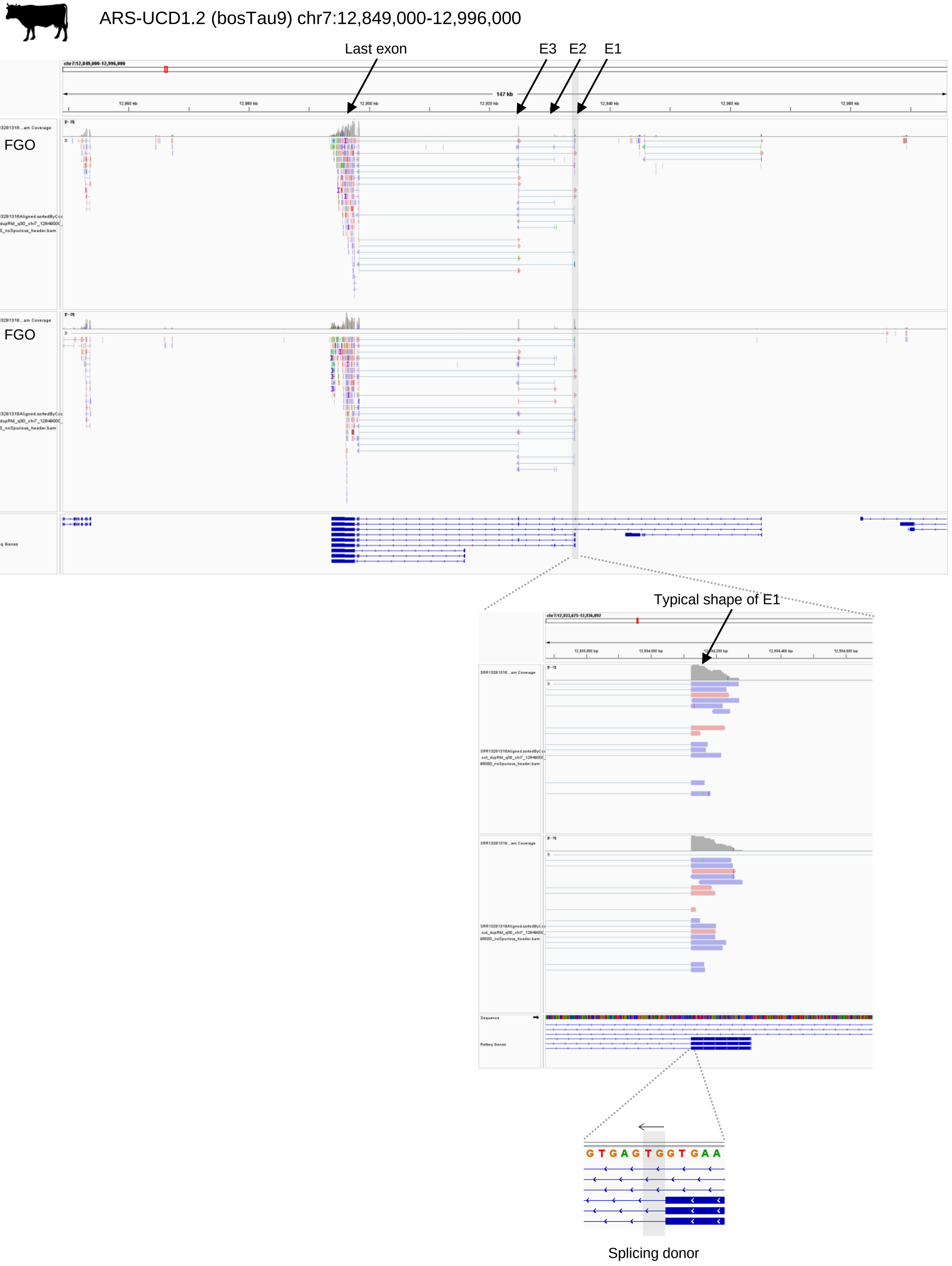


**Supplementary Figure 17. Expressed transcripts at the *ZNF791* locus in cow oocytes.** Detailed RNA-seq read coverages in full grown oocytes (FGOs) from cows (the same data used in Fig. 5) is shown using integrative genome viewer (IGV).

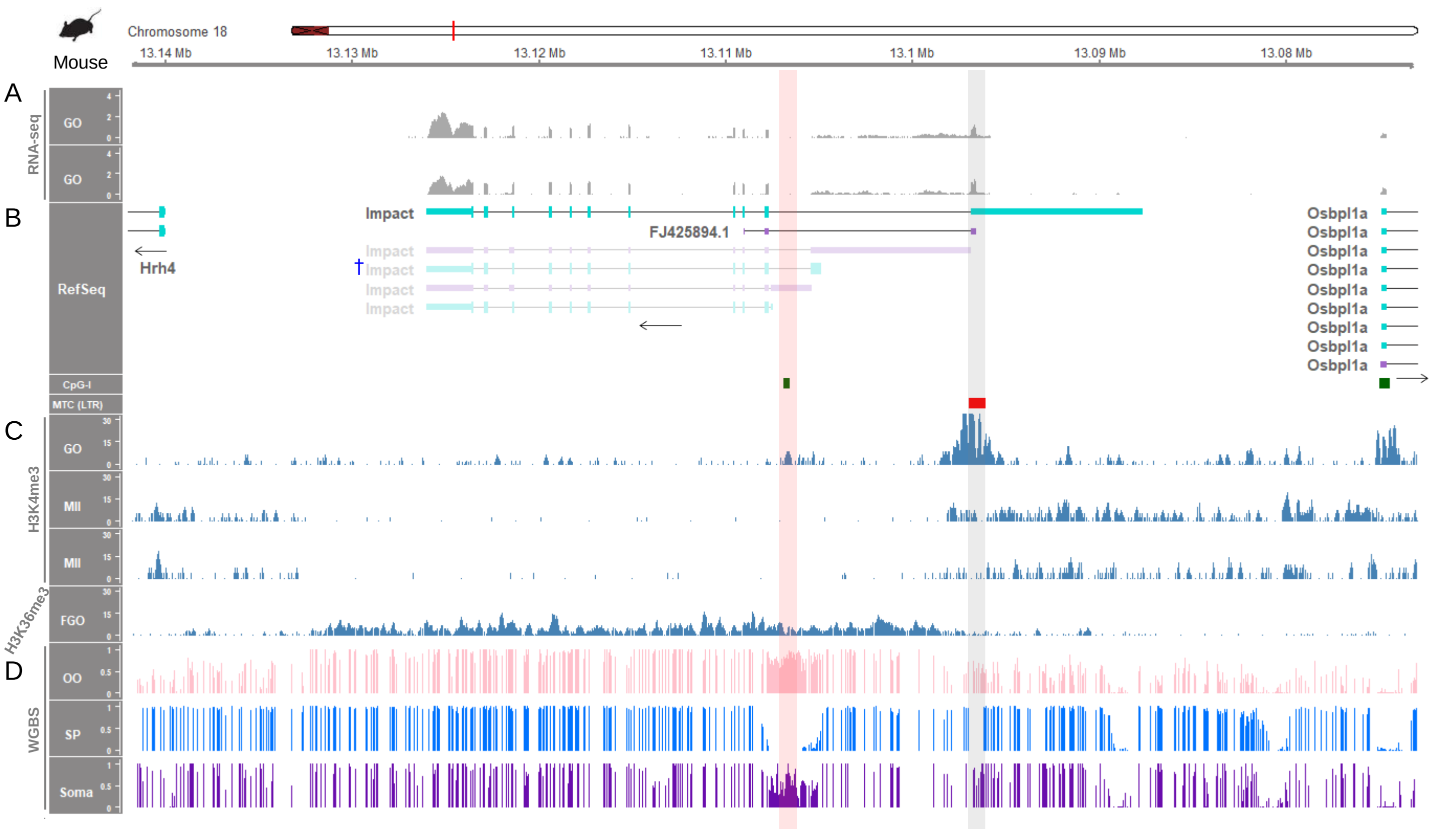


**Supplementary Figure 18. LTR-initiated transcription and establishment of methylation imprint in mouse oocytes.** This mechanism of imprinting was previously described for the known imprinted gene, mouse *Impact* (Bogutz, et al. 2019). **(A, B)** RNA-seq read coverages from growing oocytes (GOs) and the expressed *Impact* transcript. Besides the NCBI RefSeq annotation, the alternative short 1st exon was supported by an EST sequence (FJ425894.1) from the UCSC genome browser database. The LTR element (MTC) located around the TSS are marked with the red rectangle and highlighted with grey shading. Non-expressed transcripts are shown as faded. **(C)** Histone modifications, H3K4me3 and H3K36me3, in oocytes. **(D)** WGBS methylation ratios of mouse samples. Full methylation in oocytes (OO), unmethylation in sperm (SP), and partial methylation in somatic tissue (kidney) are highlighted with red shading. From the red-shaded region surrounding the intron 1 CpG island (CpG-I), the *Impact* transcript, indicated by a † in (B), is transcribed at later stages as an imprinted transcript. In addition, the rat *Impact* gene is also imprinted, but the *IMPACT* gene in humans and pigs are not imprinted (geneimprint.com).

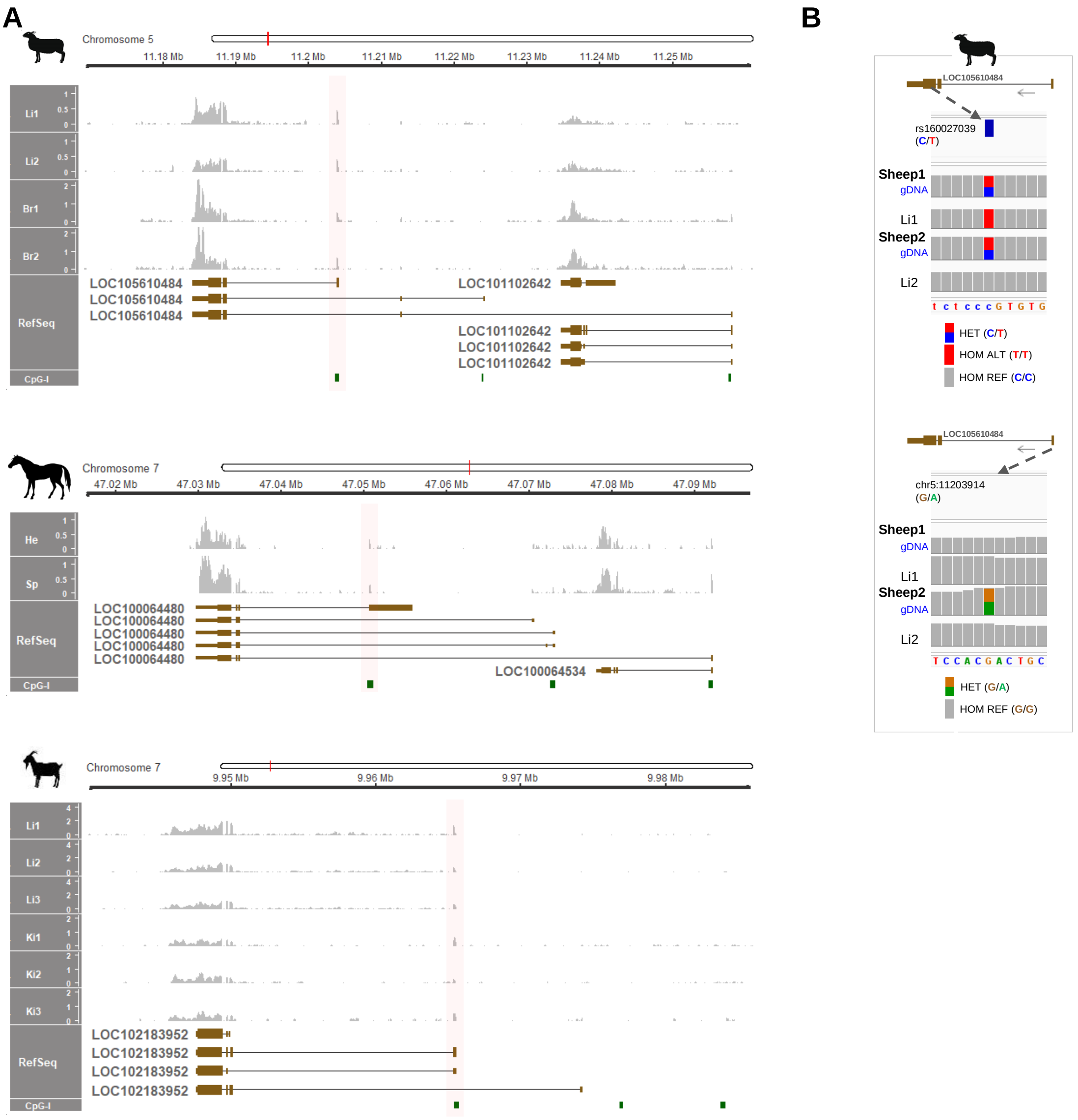


**Supplementary Figure 19. Expressed *ZNF791* transcripts in sheep, horses, and goats. (A)** Highlights in red are on the 1st exon of expressed transcripts from sheep, horses, and goats. For horses, reads were aligned to the equCab3 reference genome, displaying the short 1st exon as annotated in the equCab2 genome. RNA-seq data for sheep liver were derived from the GEO database under accession number PRJEB19199. RNA-seq data from brain (hypothalamus) of approximately 4-month-old sheep are from our published data (GSE253249). Adult horse heart (He) and spleen (Sp) RNA-seq data are from a dataset under accession number PRJEB26787. RNA-seq data for 3-year-old goats (Li, liver; Ki, kidney) are from a dataset under accession number GSE77020. **(B)** Both sheep liver RNA-seq and WGS data are from the same individuals under accession number PRJEB19199. Informative SNPs were as follows: sheep1 gDNA (C:T=5 reads: 6 reads; 45%:55%) and liver mRNA (C:T=0 read /12 reads, 0%:100%), Sheep2 gDNA (C:T=6 reads: 8 reads, 43%:57%) and liver mRNA (C:T=14 reads: 0 read, 100%:0%), Sheep2 gDNA (G:A=7 reads: 7 reads, 50%:50%) and liver mRNA (G:A=16 reads: 1 read, 94%:6%).

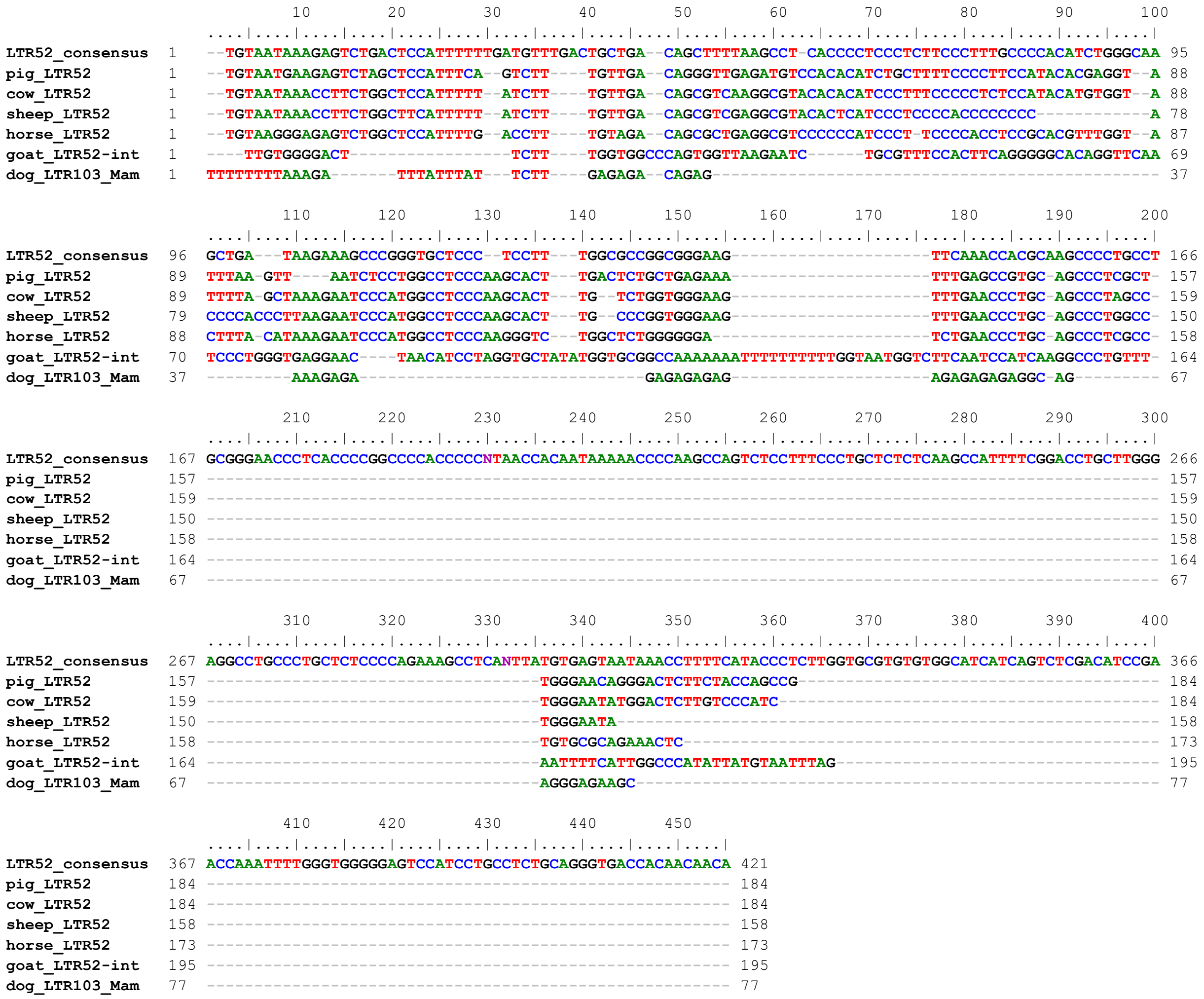


**Supplementary Figure 20. Multiple sequence alignment of LTRs.** The consensus LTR52 sequence and each matched LTR sequence were derived from the Dfam database and were aligned using the MUSCLE program.

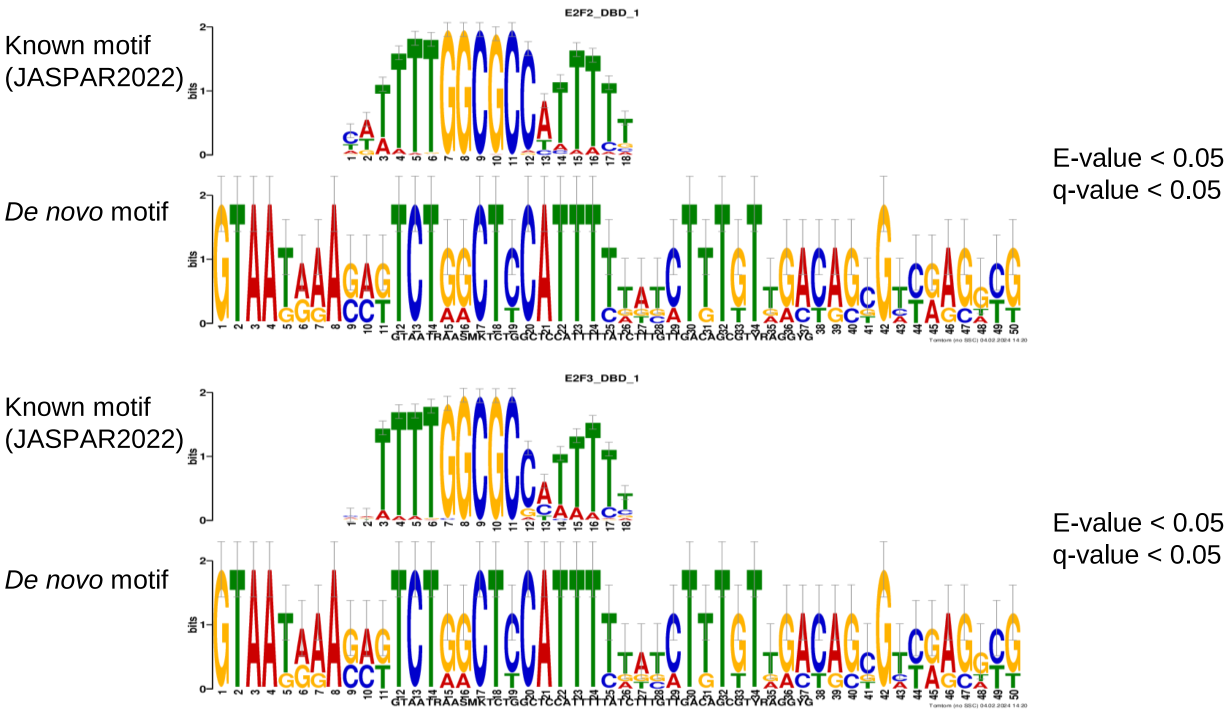


**Supplementary Figure 21. Motif discovery and comparison.** *De novo* motifs identified by the MEME tool that were common in five LTR52 sequences detected in this study (E-value < 0.05). They were found to be matched with known transcription factor binding motifs, E2F2 and E2F3 from the JASPAR2022 CORE vertebrates non-redundant database by the TOMTOM tool (E-value < 0.05, *q*-value < 0.05). The color for A and T were maintained as in the output, not converting to green and red, respectively, as in Fig. 7.

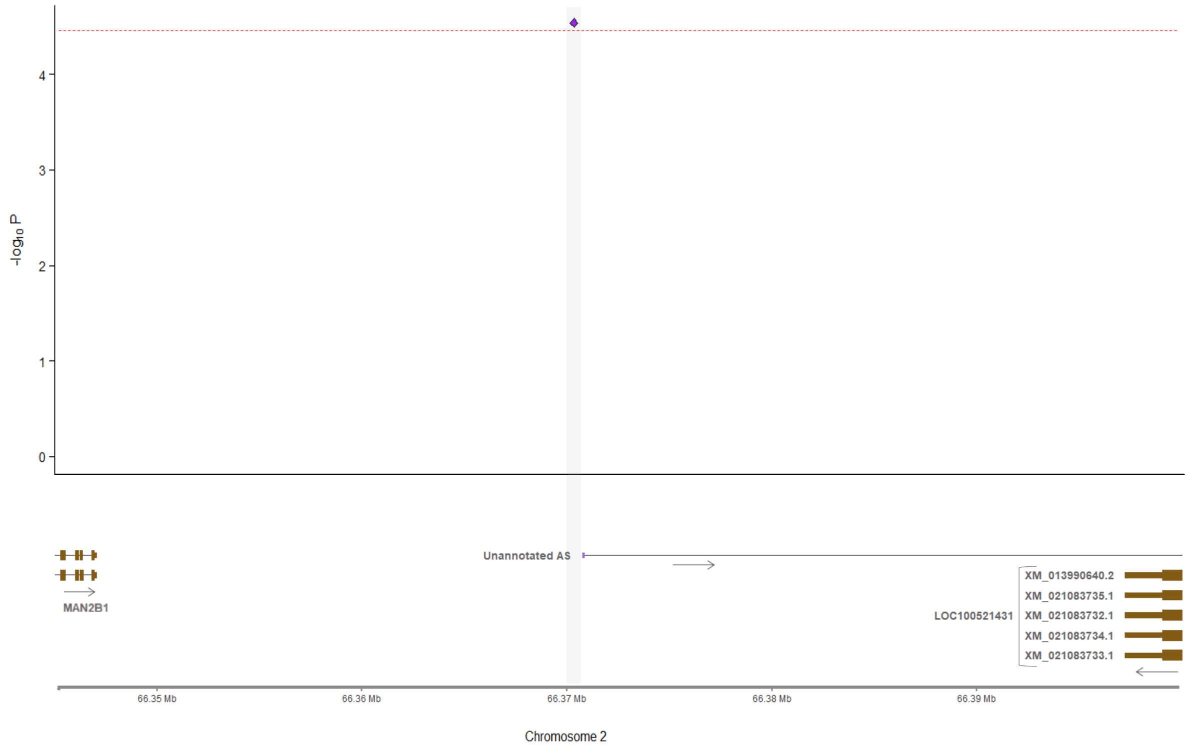


**Supplementary Figure 22. QTL analysis for the upstream of the unannotated antisense transcript.** Location (chr2:66370420) of a SNP (rs81272049) associated with residual feed intake (RFI) trait of pigs ($p$-value= $2.89 \times 10^{-5}$). The red dashed line shows suggestive significance with a $p$-value threshold of $3.09 \times 10^{-5}$. The suggestive significance and genome-wide significance were set as $p$ = 1/N and $p$ = 0.05/N, respectively, where N is the number of analyzed SNPs (Lander and Kruglyak, 1995). The genome-wide significance was $p$-value of $1.55 \times 10^{-6}$. The data were derived from pigQTLdb under http://www.animalgenome.org/QTLdb/ based on a recent GWAS (Li et al., 2022).
