## Supplementary Figure 16 for "Lineage-specific genomic imprinting in the *ZNF791* locus"

**Supplementary Figure 16. Dfam database search process.** The Dfam database was queried with input sequence from each species. These sequences were searched for transposable elements (TEs) specific to each organism using profile hidden Markov models (HMMs). This process was carried out in the following order:

1. Dfam database (<https://www.dfam.org/home>)

2. Pig (*Sus scrofa*) sequence

3. Cattle (*Bos taurus*) sequence

4. Sheep (*Ovis aries*) sequence

5. Horse (*Equus caballus*) sequence

6. Goat (*Capra hircus*) sequence

7. Dog (*Canis lupus familiaris*) sequence

**1. Dfam database (**[**https://www.dfam.org/home**](https://www.dfam.org/home)**)**

1.1. Sequence search screen

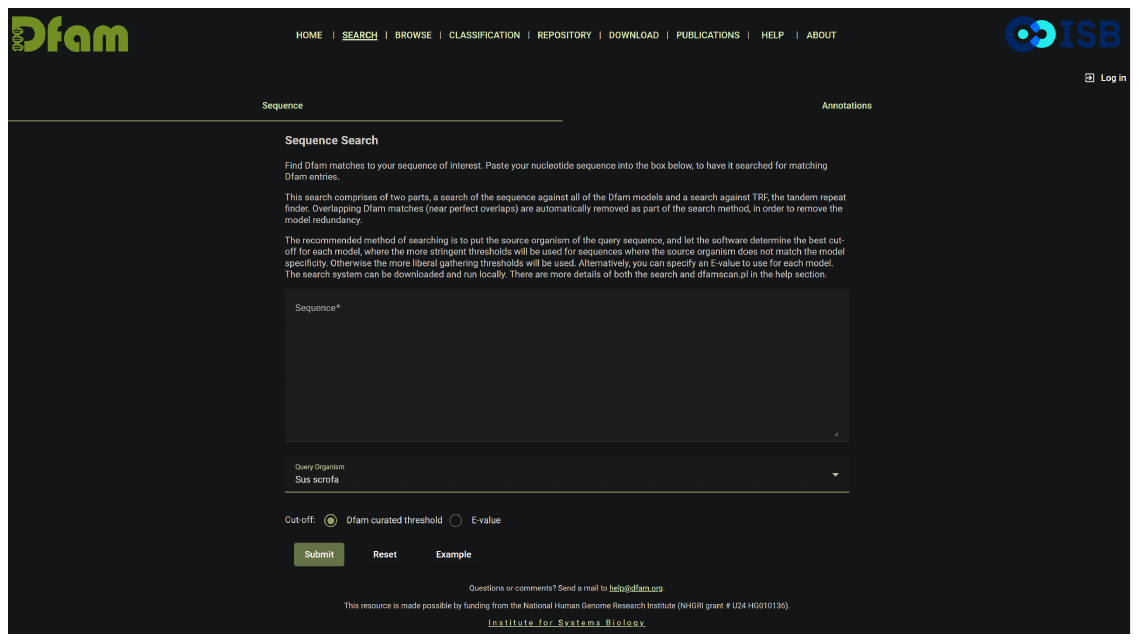

1.2. FASTA-formatted consensus LTR52 sequence, deposited in the Dfam database

>DF000000543.4 LTR52

TGTAATAAAGAGTCTGACTCCATTTTTTGATGTTTGACTGCTGACAGCTTTTAAGCCTCA

CCCCTCCCTCTTCCCTTTGCCCCACATCTGGGCAAGCTGATAAGAAAGCCCGGGTGCTCC

CTCCTTTGGCGCCGGCGGGAAGTTCAAACCACGCAAGCCCCTGCCTGCGGGAACCCTCAC

CCCGGCCCCACCCCCNTAACCACAATAAAAACCCCAAGCCAGTCTCCTTTCCCTGCTCTC

TCAAGCCATTTTCGGACCTGCTTGGGAGGCCTGCCCTGCTCTCCCCAGAAAGCCTCANTT

ATGTGAGTAATAAACCTTTTCATACCCTCTTGGTGCGTGTGTGGCATCATCAGTCTCGAC

ATCCGAACCAAATTTTGGGTGGGGGAGTCCATCCTGCCTCTGCAGGGTGACCACAACAAC

A

**2. Pig (*Sus scrofa*) sequence**

2.1. Sequence search input (*ZNF791* E1 + 1kb upstream)

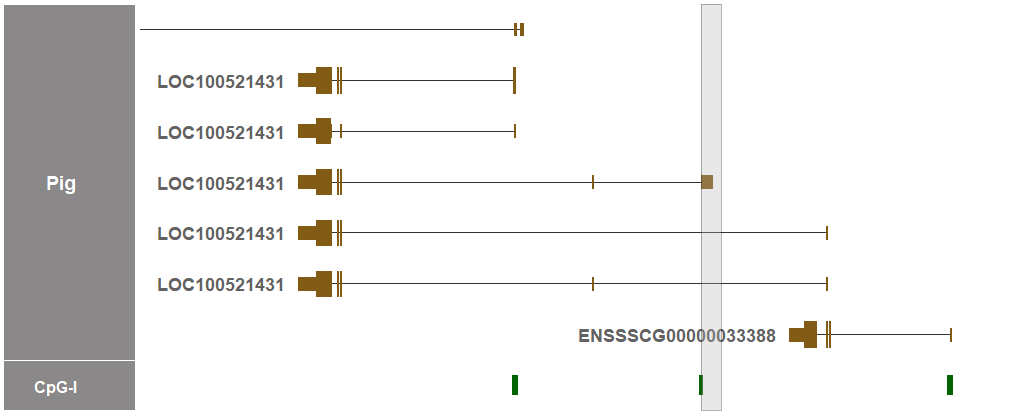

>chr2:66439732-66441912

CCATTTCCCCGCTTCTGGGATGCTCCGGCGTCACACCTTAAACTTCTACGACCCAAACAG

CTCACGGCAGGAATAACGCTGCAGCTGACAAACCTGGGCCGCTTAAGGTCCGTTGCGACT

TGACGCCAAATCTGGCCAGGAGGGTCCACGTGGTTTGTGGAGCATCGCCCAACCCACCTA

CCCCTGCTGCGTCCGTGATTGGACAATTCGCAAGCACCCGCCCCCTTTTCCTGAGTGAAG

TAGGTCTGGAGGTCACGTGATTGATCCCAccagcttctttccttaaacgTGATTGTGTAT

CTTAGAGAGCGACGTTTGCTTTTAAAACCACACTTGTTCCTGGCCCTAGAAGATGTGGCC

GTCTTCCTCAGATTTTTCAGAACGCGGGTACAGTGGTGTGCGCAGGGAATTCCAGAGTCA

AATTCTAGAAGAACCAACATACTTGTCCAAAGCTAGAGGGTCGCCTGGGTCTGGCCAACT

GGGACGATAGTTGGCATAGGCTCCGAAGCCACTCGTGGTACTGTGACCCTTGTTTACAAA

CACACTGAAAAATAACCTCACCACGTGTAATGAAGAGTCTAGCTCCATTTCAGTCTTTGT

TGACAGGGTTGAGATGTCCACACATCTGCTTTTCCCCTTCCATACACGAGGTATTTAAGT

TAATCTCCTGGCCTCCCAAGCACTTGACTCTGCTGAGAAATTTGAGCCGTGCAGCCCTCG

CTTGGGAACAGGGACTCTTCTACCAGCCGTAAGTTGTAAGATGTGTTCAGTAATACTCAG

TATACATTTTATCCTacaaggaaatttattttaattttgtagtgTGTGCCTAGAGAGTTT

GAAATGTAAATGGCTATGATATATTTAGGTCAATCCAGTTACTATTTTCACCAGTATAAC

AAAAACCAGCCAAAAGTTATGGAAATGTCGGATACCTGCAAACTTTTAATATTCTTAACA

CATTGGTTCAAAACTTCCAGtgcatttatttatcttaataGTTTTACTGAGGTAAAATAC

ACATACCATACACTGCATCCCTTAAATGattacaattcagtggttttcagtGTATTCACA

GGTATGCGCAATCAGCACCACAATTTTTGAACATTACTCTCACTTCTAAAAGAAACCCCT

GAAGGAGAGAGACAGTctcatcgggttaaggacctgacattgtctccctgaggatgcagg

ttcaatccctggttttgctcagtgggttaaggatccagcgttgccacaagctttggtgta

ggtaacagatgatacttggatccagtgtggctgtggctgtggcatagcctcagctgcagc

tcagattcaacccccaccccaccccacccccagtaagggaacttccatatgatgcaggtg

ggccataaaaaggaaaaagaaaaaaagagaaaccccTGCCCCCTTTAGAAGTTCTTCTCT

ATCTTCCTATCTTCCTCCAAGCTACAATTAGACAACCTCTAATCTTTGTTTCTGTAGATT

TCCCTATTCTTCATTTACATATGAATGGACTCACATAATGTATTGTCTTTGTAATTGACT

CTGGTTGCTTAGCATAACTTTTTCAAGAAACATGTATgctgagttccctgatggcctagt

ggctaaaaaaaatttcaaaattcaaaccaaaataaaccataaaaaaagaaacatctatgC

TGTAGCATGTATGGATACCTATTTGtctttatggccaaataatattctactgtatggata

aaacatatttattcacCCCTTAATGGATTTACTTTGTTCCTacgttttggctattatgaa

taatgttgcaataaaCTTTCATGGAGAAGTTTTTGTTTAGAcgtattttcatttctattt

ggcaGATTCCTAGGAATGAATTGATGAGTCTTAAGGTAATTCTAGGTTTAATTGCTAAGG

AACTTTTCCCCaaaagactgttttccaaagtggctgcaccattttatattcccaccagca

gtgggtGAGTGTTCCAGTTTTTCCAAATCATCCACAAAACTtattatctcactttttttt

tttttgtattttttagggcaacacctggaatatggaagttcccagtctactCAGAaccga

actgcagctgccagcctaagccacagccacagcaatgccagatctaagcaatgctggata

cttacccactgagtgaggaca

2.2. Dfam results from profile HMMs

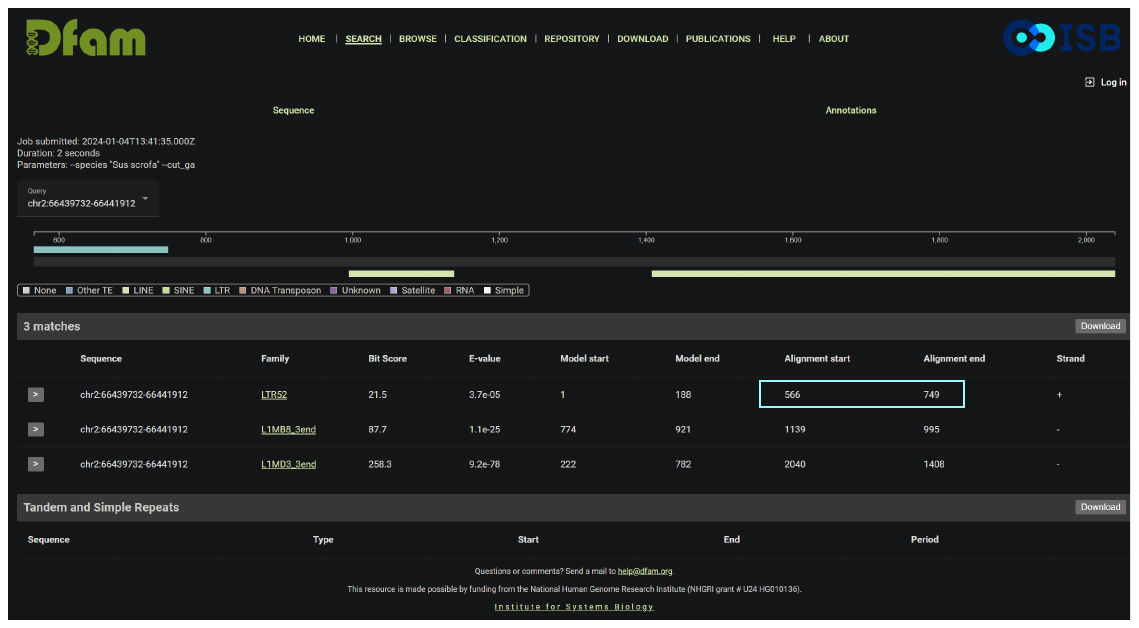

Using *Sus scrofa* as a query organism and the Dfam curated threshold, the LTR52 (Model) was

detected within the query input (Alignment) as shown in cyan.

>chr2:66439732-66441912

CCATTTCCCCGCTTCTGGGATGCTCCGGCGTCACACCTTAAACTTCTACGACCCAAACAG

CTCACGGCAGGAATAACGCTGCAGCTGACAAACCTGGGCCGCTTAAGGTCCGTTGCGACT

TGACGCCAAATCTGGCCAGGAGGGTCCACGTGGTTTGTGGAGCATCGCCCAACCCACCTA

CCCCTGCTGCGTCCGTGATTGGACAATTCGCAAGCACCCGCCCCCTTTTCCTGAGTGAAG

TAGGTCTGGAGGTCACGTGATTGATCCCAccagcttctttccttaaacgTGATTGTGTAT

CTTAGAGAGCGACGTTTGCTTTTAAAACCACACTTGTTCCTGGCCCTAGAAGATGTGGCC

GTCTTCCTCAGATTTTTCAGAACGCGGGTACAGTGGTGTGCGCAGGGAATTCCAGAGTCA

AATTCTAGAAGAACCAACATACTTGTCCAAAGCTAGAGGGTCGCCTGGGTCTGGCCAACT

GGGACGATAGTTGGCATAGGCTCCGAAGCCACTCGTGGTACTGTGACCCTTGTTTACAAA

CACACTGAAAAATAACCTCACCACGTGTAATGAAGAGTCTAGCTCCATTTCAGTCTTTGT

TGACAGGGTTGAGATGTCCACACATCTGCTTTTCCCCTTCCATACACGAGGTATTTAAGT

TAATCTCCTGGCCTCCCAAGCACTTGACTCTGCTGAGAAATTTGAGCCGTGCAGCCCTCG

CTTGGGAACAGGGACTCTTCTACCAGCCGTAAGTTGTAAGATGTGTTCAGTAATACTCAG

TATACATTTTATCCTacaaggaaatttattttaattttgtagtgTGTGCCTAGAGAGTTT

GAAATGTAAATGGCTATGATATATTTAGGTCAATCCAGTTACTATTTTCACCAGTATAAC

AAAAACCAGCCAAAAGTTATGGAAATGTCGGATACCTGCAAACTTTTAATATTCTTAACA

CATTGGTTCAAAACTTCCAGtgcatttatttatcttaataGTTTTACTGAGGTAAAATAC

ACATACCATACACTGCATCCCTTAAATGattacaattcagtggttttcagtGTATTCACA

GGTATGCGCAATCAGCACCACAATTTTTGAACATTACTCTCACTTCTAAAAGAAACCCCT

GAAGGAGAGAGACAGTctcatcgggttaaggacctgacattgtctccctgaggatgcagg

ttcaatccctggttttgctcagtgggttaaggatccagcgttgccacaagctttggtgta

ggtaacagatgatacttggatccagtgtggctgtggctgtggcatagcctcagctgcagc

tcagattcaacccccaccccaccccacccccagtaagggaacttccatatgatgcaggtg

ggccataaaaaggaaaaagaaaaaaagagaaaccccTGCCCCCTTTAGAAGTTCTTCTCT

ATCTTCCTATCTTCCTCCAAGCTACAATTAGACAACCTCTAATCTTTGTTTCTGTAGATT

TCCCTATTCTTCATTTACATATGAATGGACTCACATAATGTATTGTCTTTGTAATTGACT

CTGGTTGCTTAGCATAACTTTTTCAAGAAACATGTATgctgagttccctgatggcctagt

ggctaaaaaaaatttcaaaattcaaaccaaaataaaccataaaaaaagaaacatctatgC

TGTAGCATGTATGGATACCTATTTGtctttatggccaaataatattctactgtatggata

aaacatatttattcacCCCTTAATGGATTTACTTTGTTCCTacgttttggctattatgaa

taatgttgcaataaaCTTTCATGGAGAAGTTTTTGTTTAGAcgtattttcatttctattt

ggcaGATTCCTAGGAATGAATTGATGAGTCTTAAGGTAATTCTAGGTTTAATTGCTAAGG

AACTTTTCCCCaaaagactgttttccaaagtggctgcaccattttatattcccaccagca

gtgggtGAGTGTTCCAGTTTTTCCAAATCATCCACAAAACTtattatctcactttttttt

tttttgtattttttagggcaacacctggaatatggaagttcccagtctactCAGAaccga

actgcagctgccagcctaagccacagccacagcaatgccagatctaagcaatgctggata

cttacccactgagtgaggaca

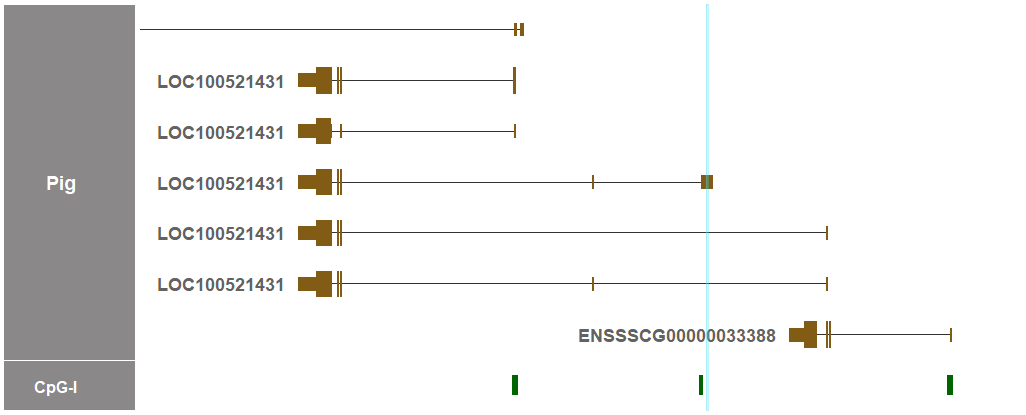

The detected region of LTR52 is indicated in cyan in both the sequence above and the genome visualizer.

2.3. Location of the detected region of LTR52 in the UCSC Genome Browser track

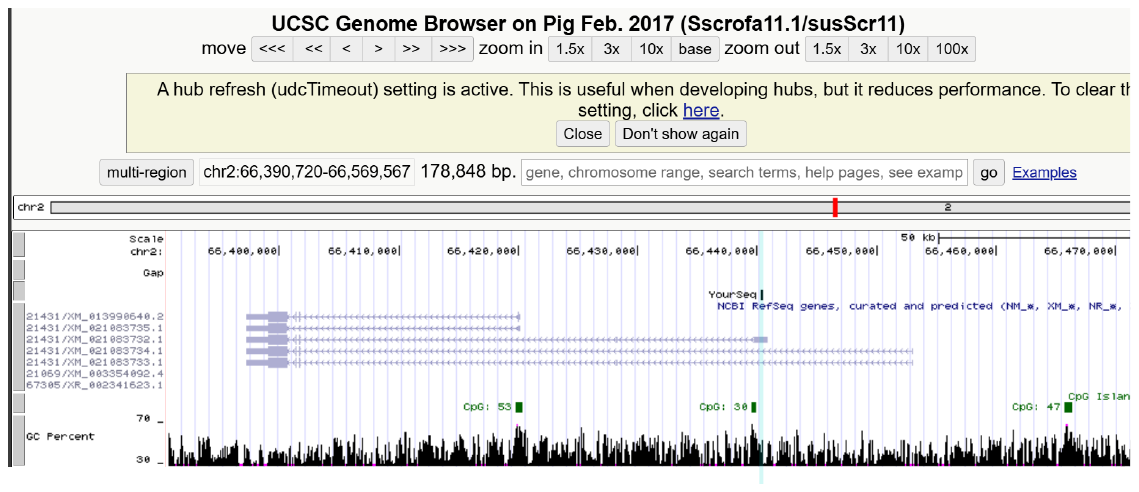

The above sequence in cyan was used an input (YourSeq) for the UCSC BLAT search.

**3. Cattle (*Bos taurus*) sequence**

3.1. Sequence search input (*ZNF791* E1 + 1kb upstream)

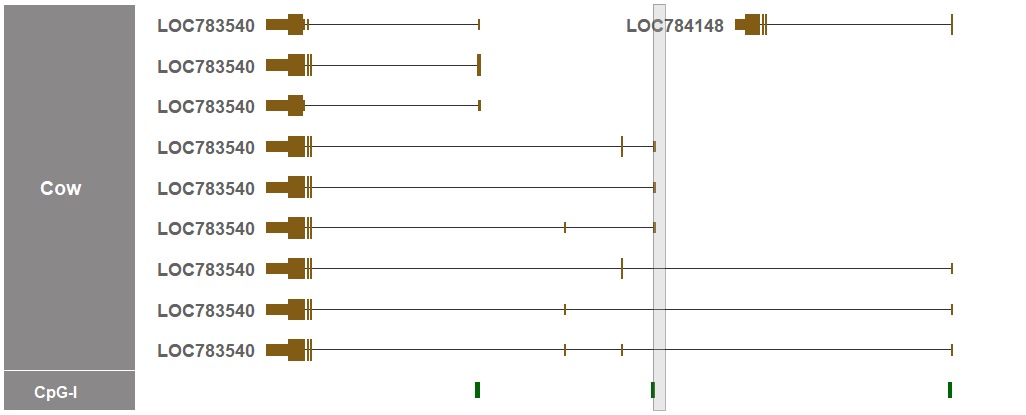

>chr7:12934124-12935308

CACTTCACGCCTTCCGGGGTTCTCCGGCGTTATTCCTTAGGCTCACGCGGCCTCTGCAGC

TCATAGCGGTAATGAAGCTGCAGCGAACTAACCTGAGCTGCTTCAACACGGCGACACTCC

ACGCAGAGCGTGCCGGGAGAGGCCGACCACACGGTTTGCGGGACGTGGCCCCACCCACCT

TCCCCTTCTGCGGTCCTGATTGGACAGTTCGCAAGCAACGCCCCCTTTATCCTGAGTAAT

AATGGATCCAGAGGCCACGTGCCTGAGCAGAATCAGCTTCTGCCGGTGGGTAGAAACCTA

CCCCTCTCCAGggacttttgtttttaaaccactCTTGTTACTTGTCCTAGAGGGACAGTG

GCAACTTCCTCAGATACTTCAGAACGTGGGGACACAGGTGTGCACAAGGAATTCCAGAGT

CAGATCCCAGAAGAGCCGACGTTCTTGTCCAAAGCAAGAGCAGGCCAACTGGAAAAACAG

GGTCTGAGGGTCCTCCATCGGGAAAATAGTTCGCACAGACTGCGGAGCTACTGATGATAA

CTATGACACTCGATTTCAAACACACTGAAAAATATCCTCACCATGTGTAATAAACCTTCT

GGCtccatttttatctttgttgACAGCGTCAAGGCGTACACACATCCCTTTCCCCCTCTC

CATACATGTGGTATTTTAGctaaagaatcccatggcctCCCAAGCACTTGTCTGGTGGGA

AGTTTGAACCCTGCAGCCCTAGCCTGGGAATATGGACTCTTGTCCCATCAACCAGAAGTT

GTGCGATGTGTTCAATAATACTCTCAAGTATACATGTTACGTGAGAAGGAAATTTTAGTT

TCGTAGTATGTGCCTAGAGATTTTGAAATGTAAACGGCTATGATATCCCAAAGGCATTTC

TGTTACTATTGTCACCAATATaacaagaaacaataaaaagtttTGGAAATGAACCATACA

CTTGAAAACTTTTGATATTCTTAACATACTGGTTCAAAACTTCTAATGAATTTATCTGAA

TAGTTTTATGaggtaaaatacacacacatttcctgcaagactccgatgctgggaaagatt

gagaacaaaaggataagagggtgacagaggatgagatggatggcatcactgactcaaagg

atgtgagtttgagcaaactctgggacatagtgaaggacagggaag

3.2. Dfam results from profile HMMs

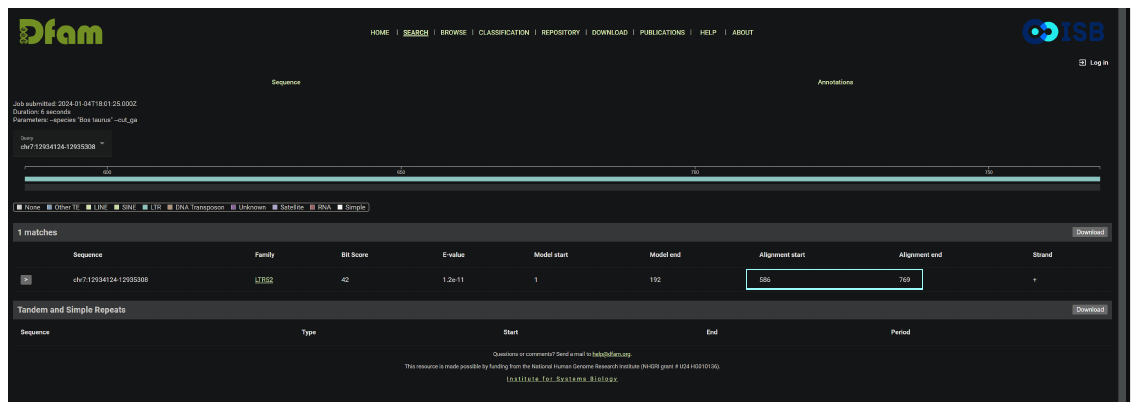

Using *Bos taurus* as a query organism and the Dfam curated threshold, only the LTR52 (Model) was

detected within the query input (Alignment) as shown in cyan.

>chr7:12934124-12935308

CACTTCACGCCTTCCGGGGTTCTCCGGCGTTATTCCTTAGGCTCACGCGGCCTCTGCAGC

TCATAGCGGTAATGAAGCTGCAGCGAACTAACCTGAGCTGCTTCAACACGGCGACACTCC

ACGCAGAGCGTGCCGGGAGAGGCCGACCACACGGTTTGCGGGACGTGGCCCCACCCACCT

TCCCCTTCTGCGGTCCTGATTGGACAGTTCGCAAGCAACGCCCCCTTTATCCTGAGTAAT

AATGGATCCAGAGGCCACGTGCCTGAGCAGAATCAGCTTCTGCCGGTGGGTAGAAACCTA

CCCCTCTCCAGggacttttgtttttaaaccactCTTGTTACTTGTCCTAGAGGGACAGTG

GCAACTTCCTCAGATACTTCAGAACGTGGGGACACAGGTGTGCACAAGGAATTCCAGAGT

CAGATCCCAGAAGAGCCGACGTTCTTGTCCAAAGCAAGAGCAGGCCAACTGGAAAAACAG

GGTCTGAGGGTCCTCCATCGGGAAAATAGTTCGCACAGACTGCGGAGCTACTGATGATAA

CTATGACACTCGATTTCAAACACACTGAAAAATATCCTCACCATGTGTAATAAACCTTCT

GGCtccatttttatctttgttgACAGCGTCAAGGCGTACACACATCCCTTTCCCCCTCTC

CATACATGTGGTATTTTAGctaaagaatcccatggcctCCCAAGCACTTGTCTGGTGGGA

AGTTTGAACCCTGCAGCCCTAGCCTGGGAATATGGACTCTTGTCCCATCAACCAGAAGTT

GTGCGATGTGTTCAATAATACTCTCAAGTATACATGTTACGTGAGAAGGAAATTTTAGTT

TCGTAGTATGTGCCTAGAGATTTTGAAATGTAAACGGCTATGATATCCCAAAGGCATTTC

TGTTACTATTGTCACCAATATaacaagaaacaataaaaagtttTGGAAATGAACCATACA

CTTGAAAACTTTTGATATTCTTAACATACTGGTTCAAAACTTCTAATGAATTTATCTGAA

TAGTTTTATGaggtaaaatacacacacatttcctgcaagactccgatgctgggaaagatt

gagaacaaaaggataagagggtgacagaggatgagatggatggcatcactgactcaaagg

atgtgagtttgagcaaactctgggacatagtgaaggacagggaag

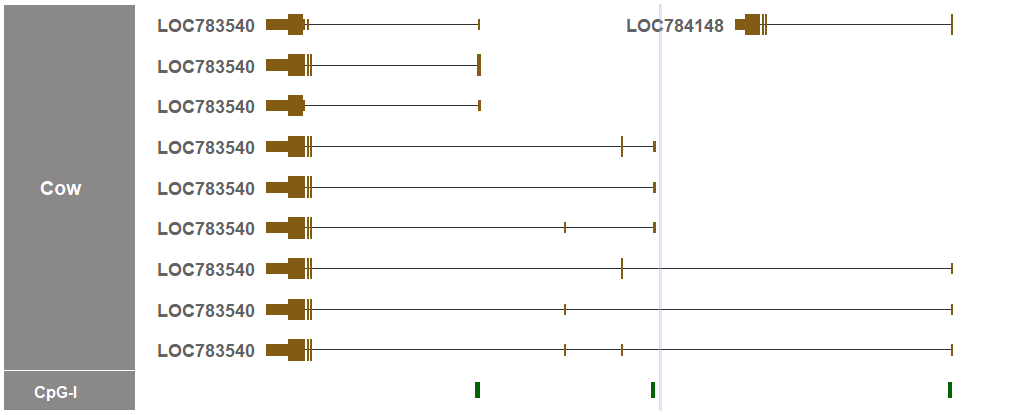

The detected region of LTR52 is indicated in cyan in both the sequence above and the genome visualizer.

3.3. Location of the detected region of LTR52 in the UCSC Genome Browser track

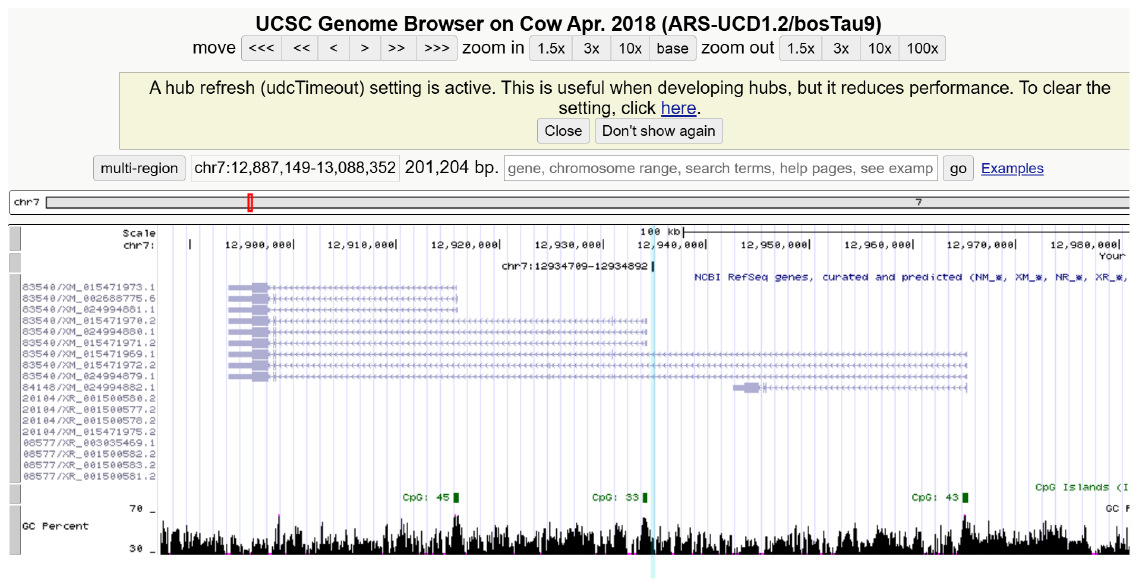

The above sequence in cyan was used an input for the UCSC BLAT search.

**4. Sheep (*Ovis aries*) sequence**

4.1. Sequence search input (*ZNF791* E1 + 1kb upstream)

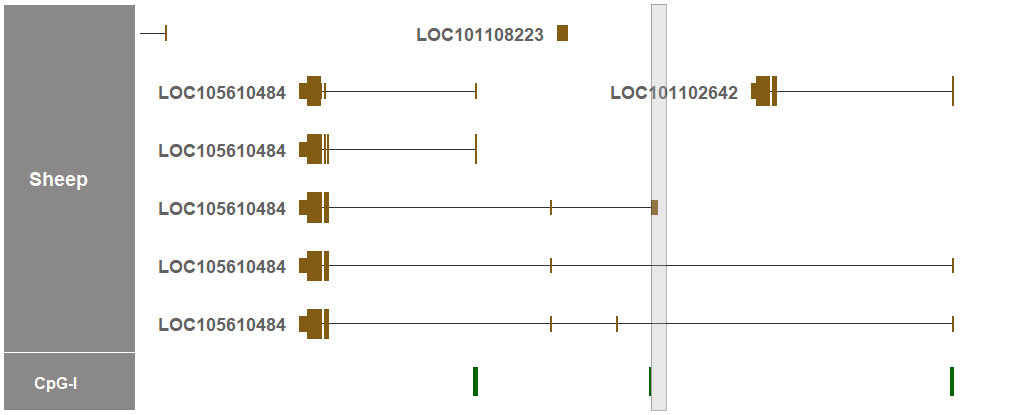

>chr5:10951059-10952708

CACTTCACACCTTCTGGGGTTCTCCAGAGACATTCCTTAGACTCACGCGGCCTCTGCAGC

TCACAGCGGTGATGAAGCTGCAGCGAACTAACCTGAGCTGCTTCAACGCTGGTGAAACTC

CACACAGAGCGTGCTAGGAGCCATCACAGGGTCTGCGGGACGTGGCCCCACCCACCTTCC

CCTGCTGCACTTCTGATTGGACAGTTTGCAAGCACAGCTCCCTTTTTCCTGAGTAATAAT

TGGTCCAGAGGCCACGTGGCTGAGCAGAATCAGCTTCTGCAGGTGGGTGGAAACCTACCC

GCCTCCAGGGACTTTTGTTTTTAAACCACTCTTGTTACTTGTCCTAGAGGGACAGTGGCA

ACTTCCTCAGATACTTCAGAACGTGGGGACACAGGTGTGCACAAGGAATTCCAGAGTCAG

ATTCCAGAAGAACCGACGTTCTTGTCCAAAAAAAAGAGCAGGCCAACTGGAAAAAGAGGG

TCTGAGGTGCTCCATCGGGAAAATAGTTCGCACAGATTGCGGAGCCACTGATGATACTAG

GACACTCGATTTCAAACACACTGAAAAATATCCTCACTATGTGTAATAAACCTTCTGGCT

TCATTTTTATCTTTGTTGACAGCGTCGAGGCGTACACTcatccctccccaccccccccac

cccacccTTAAGAATCCCATGGCCTCCCAAGCACTTGCCCGGTGGGAAGTTTGAACCCTG

CAGCCCTGGCCTGGGAATATGGACTCTTCTCCCATCAATCAGAACTTGTGCGACGTGTTA

AATAATATTCTCAAGTATACATGTTACGTGAGAAGGAAATTTTAGTTTCGtatgatccat

ggggtcgccaagagtcagacacaactgagcaacttcactttcacttttcactttcatgca

ttggagaaggaaatggcaacccactccagtgttcttgcctggagaatcccagggaggagg

gagcctggtgcgctgccatctatggggtcgcacagagtcagacacgactgaagcgactta

gcagcagcagcAGCAGCAGTATGTGCCTAGAGATTTTGAAATGTAAACGGCTATGATATC

CCAAAGGCAATTCAGTTACTATTGTCACCAATATAACAAGAAACAATAAAAAGTTTTGGA

AATGTACCATACACTTGAAAACTTTTGATATACTTAACATACTGGTTCAAAACTTCTAAT

GAATTTATCTGAATAGCTTTATGAGGTAAAATACACATACATTTCTTGCaagactctgat

gctgggaaagattgagcacaaaaggataagagggtgacagaggatgagatggttggatgg

cattaccgactcaaagggacatgagtttgagcaagctccaggagttggtgatggacaggg

aagcctggtgtgctgcagtcaaagaatcagacaccactgagggacGAAATAACAATAACC

ACTAAAGGGATGTTGCCTACAAATATAATATATACACAAGGGCCCATCTTTGGGAACCCT

GCCTTCACCTGTAAGGAGCATTAAACTAAAATGCCTTGTTTAGCTACAGAAAACATCCTG

ATCAGGACAAGTTAATCACTAAAGGGATGTTGCCTATAAAGCTTAAATTATATGTAATGG

CTCATCGCTGGTAACCCTGGGTTCTGTAAT

4.2. Dfam results from profile HMMs

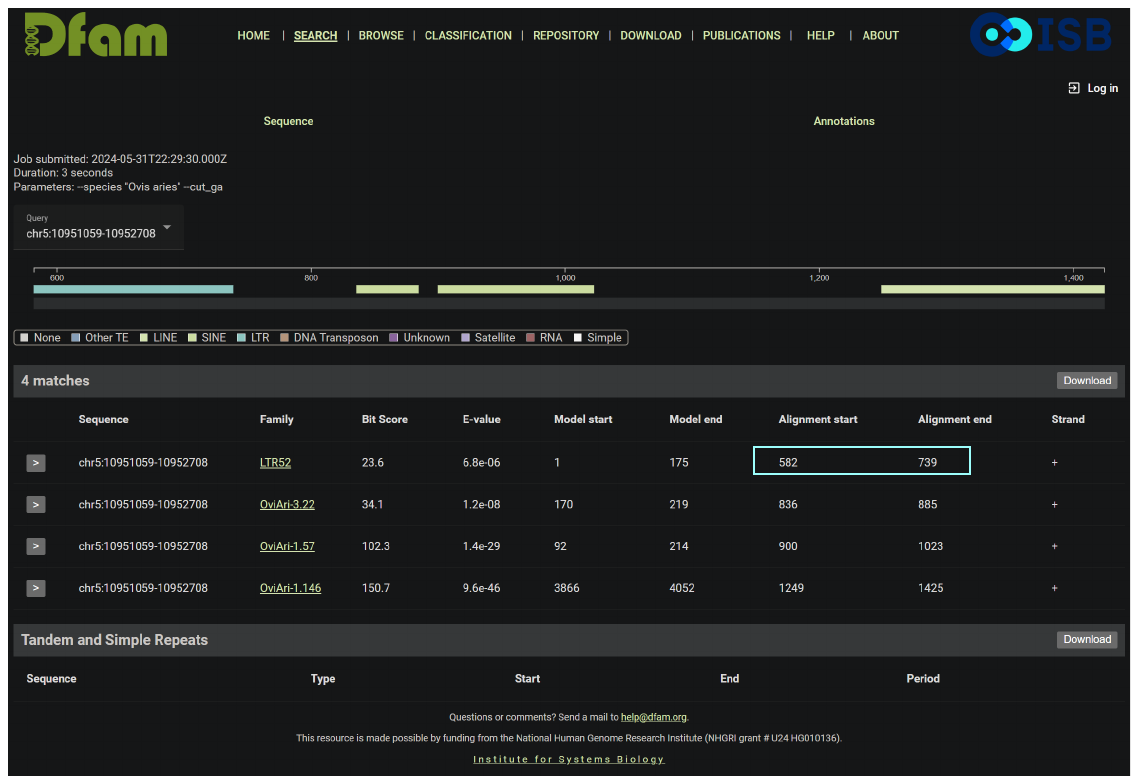

Using *Ovis aries* as a query organism and the Dfam curated threshold, the LTR52 (Model) was

detected within the query input (Alignment) as shown in cyan.

>chr5:10951059-10952708

CACTTCACACCTTCTGGGGTTCTCCAGAGACATTCCTTAGACTCACGCGGCCTCTGCAGC

TCACAGCGGTGATGAAGCTGCAGCGAACTAACCTGAGCTGCTTCAACGCTGGTGAAACTC

CACACAGAGCGTGCTAGGAGCCATCACAGGGTCTGCGGGACGTGGCCCCACCCACCTTCC

CCTGCTGCACTTCTGATTGGACAGTTTGCAAGCACAGCTCCCTTTTTCCTGAGTAATAAT

TGGTCCAGAGGCCACGTGGCTGAGCAGAATCAGCTTCTGCAGGTGGGTGGAAACCTACCC

GCCTCCAGGGACTTTTGTTTTTAAACCACTCTTGTTACTTGTCCTAGAGGGACAGTGGCA

ACTTCCTCAGATACTTCAGAACGTGGGGACACAGGTGTGCACAAGGAATTCCAGAGTCAG

ATTCCAGAAGAACCGACGTTCTTGTCCAAAAAAAAGAGCAGGCCAACTGGAAAAAGAGGG

TCTGAGGTGCTCCATCGGGAAAATAGTTCGCACAGATTGCGGAGCCACTGATGATACTAG

GACACTCGATTTCAAACACACTGAAAAATATCCTCACTATGTGTAATAAACCTTCTGGCT

TCATTTTTATCTTTGTTGACAGCGTCGAGGCGTACACTcatccctccccaccccccccac

cccacccTTAAGAATCCCATGGCCTCCCAAGCACTTGCCCGGTGGGAAGTTTGAACCCTG

CAGCCCTGGCCTGGGAATATGGACTCTTCTCCCATCAATCAGAACTTGTGCGACGTGTTA

AATAATATTCTCAAGTATACATGTTACGTGAGAAGGAAATTTTAGTTTCGtatgatccat

ggggtcgccaagagtcagacacaactgagcaacttcactttcacttttcactttcatgca

ttggagaaggaaatggcaacccactccagtgttcttgcctggagaatcccagggaggagg

gagcctggtgcgctgccatctatggggtcgcacagagtcagacacgactgaagcgactta

gcagcagcagcAGCAGCAGTATGTGCCTAGAGATTTTGAAATGTAAACGGCTATGATATC

CCAAAGGCAATTCAGTTACTATTGTCACCAATATAACAAGAAACAATAAAAAGTTTTGGA

AATGTACCATACACTTGAAAACTTTTGATATACTTAACATACTGGTTCAAAACTTCTAAT

GAATTTATCTGAATAGCTTTATGAGGTAAAATACACATACATTTCTTGCaagactctgat

gctgggaaagattgagcacaaaaggataagagggtgacagaggatgagatggttggatgg

cattaccgactcaaagggacatgagtttgagcaagctccaggagttggtgatggacaggg

aagcctggtgtgctgcagtcaaagaatcagacaccactgagggacGAAATAACAATAACC

ACTAAAGGGATGTTGCCTACAAATATAATATATACACAAGGGCCCATCTTTGGGAACCCT

GCCTTCACCTGTAAGGAGCATTAAACTAAAATGCCTTGTTTAGCTACAGAAAACATCCTG

ATCAGGACAAGTTAATCACTAAAGGGATGTTGCCTATAAAGCTTAAATTATATGTAATGG

CTCATCGCTGGTAACCCTGGGTTCTGTAAT

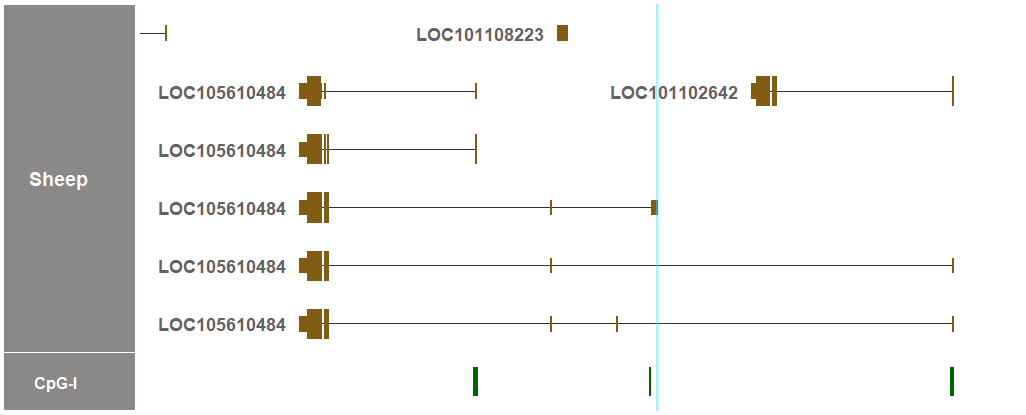

The detected region of LTR52 is indicated in cyan in both the sequence above and the genome visualizer.

4.3. Location of the detected region of LTR52 in the UCSC Genome Browser track

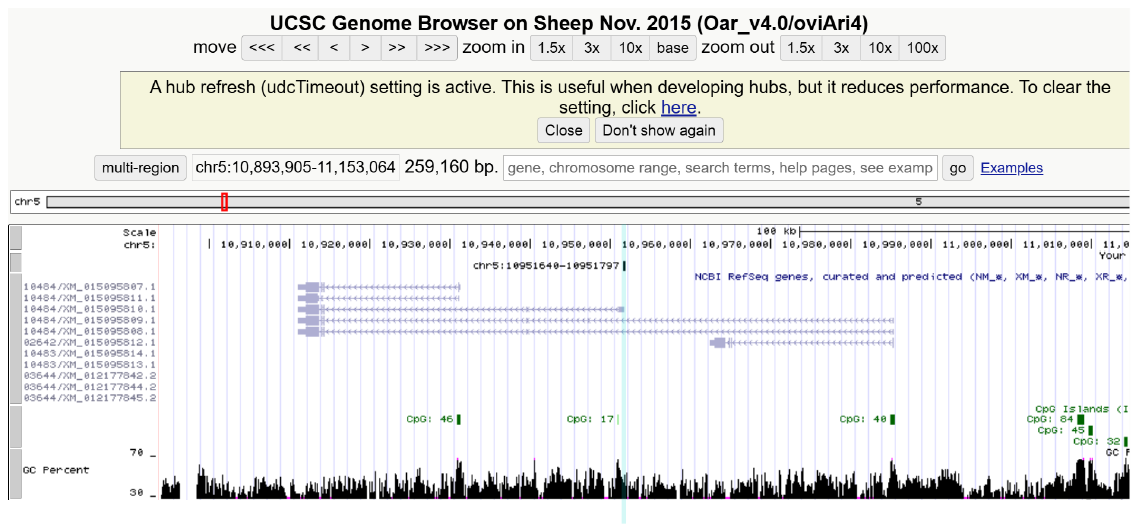

The above sequence in cyan was used an input for the UCSC BLAT search.

**5. Horse (*Equus caballus*) sequence**

5.1. Sequence search input (*ZNF791* E1 + 1kb upstream)

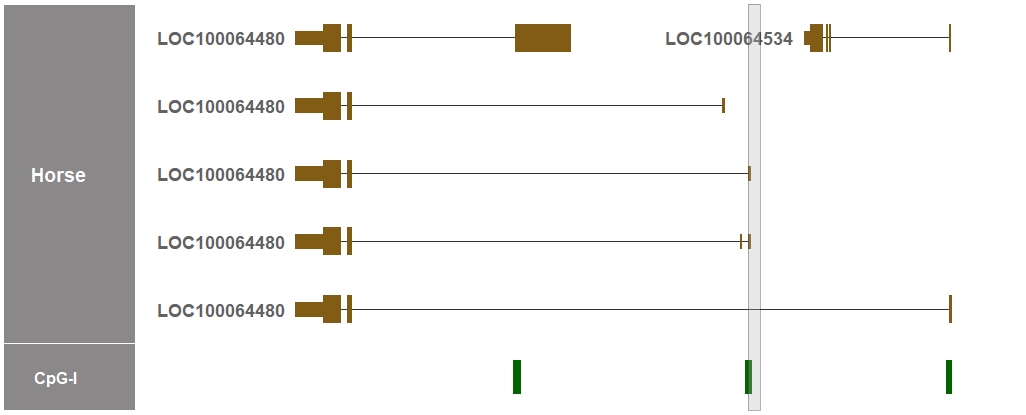

>chr7:47072841-47074025

CGGTTTCCCGGCTTCTGGGTACCCCGGCGTCGCCCCCACTCCGCCGGCGGCCGCGGCAGG

TCCCAGGGTACCGACGCCGCAGCAGAAACGCCGGAGCCCGTTAACTGCAGGTGAGACGCT

GTCTGAGCTCGGCCAAGAGCCTCCGCGCGGTCTGCGGGGCGTCGCCCCACCCACCTGCCC

CTGCTGCGCCTCTGATTGGACTGTTCGCAAGGGCCCGCCCCCTCGCTTCTTGAGTGACAG

TGGGGCAGGAGGTCAGACGCCTGAGCCGAATCAGCGTTCAGCGGTGAGCGGGAAACTGTG

TCGTCCGATGAGTGAGGTTTGTTTTTAAACAAAAGTTGTTCTTGTCCCTAAAGGGACAGT

GGCGTCTTCCTGCAGCGCCTTCTGAACGCGGAGACACAGGTGTGCAGGGAATTCCAGAGT

CAGTTGCCAGATGGAGCGATGCCCTTATCCAGAGCAAGAGGGCGCCCTAGCCAGGCTAGA

CAGGGACCAGAGGGTCTGAGGTccccgcccctccccccctccccccccagcccccGGGTC

AGGGGTTGGCATAAGGCTGGGAAGCCACTCATGGAACCGTGACTCATTTACAAACACGCT

TAAATATGACATCACCACATGTAAGGGAGAGTCTGGCTCCATTTTGACCTTTGTAGACAG

CGCTGAGGCGTCCCCCCATCCCTTCCCCACCTCCGCACGTTTGGTACTTTACATAAAGAA

TCCCATGGCCTCCCAAGGGTCTGGCTCTGGGGGGATCTGAACCCTGCAGCCCTCGCCTGT

GCGCAGAAACTCGTCTCTGACCAGCTGTGAGCTTTGAGATGGGTTCAATAAAACTCAGGT

GTACATTTTACACTAGAAGGAAATTTATTTTACTTTCTGAGCATGACCTTAAGAGATTTT

GAAATGTAAACAGCTATGATATCCCTAAGGCAACTGAATTCTGATTTCCACCAATATACA

TATAAAAACAATGTTATGTAAATGTGAGCTGTACCTGAAAACTCTTTAAGATACTTAACA

CATTGGTTCAAAAAATTCCTAATTCATTTAtttgaatagctttactgagatataatacac

atattatacaaccctttaatgtgtataattcaatggttttcagtacattccgatatgtgc

aataatcatcacaacttttgaacatttttgtggcttcaaaaagaa

5.2. Dfam results from profile HMMs

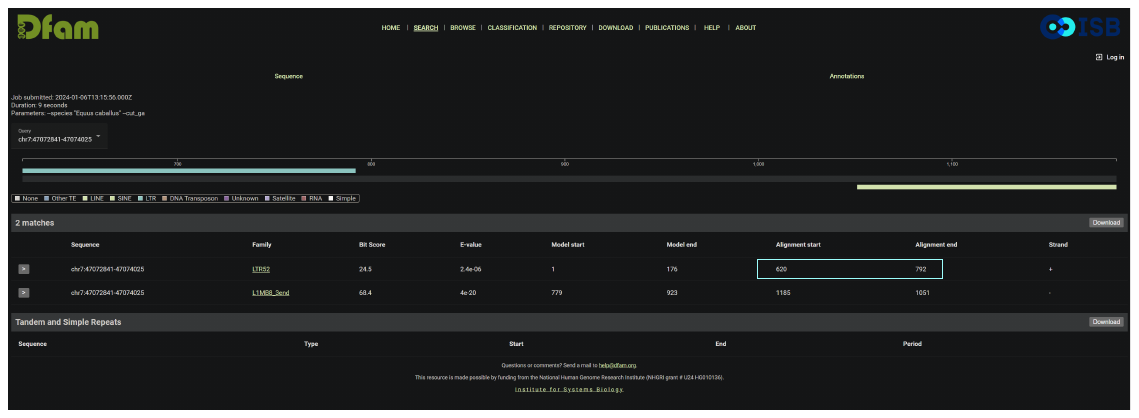

Using *Equus caballus* as a query organism and the Dfam curated threshold, the LTR52 (Model) was

detected within the query input (Alignment) as shown in cyan.

>chr7:47072841-47074025

CGGTTTCCCGGCTTCTGGGTACCCCGGCGTCGCCCCCACTCCGCCGGCGGCCGCGGCAGG

TCCCAGGGTACCGACGCCGCAGCAGAAACGCCGGAGCCCGTTAACTGCAGGTGAGACGCT

GTCTGAGCTCGGCCAAGAGCCTCCGCGCGGTCTGCGGGGCGTCGCCCCACCCACCTGCCC

CTGCTGCGCCTCTGATTGGACTGTTCGCAAGGGCCCGCCCCCTCGCTTCTTGAGTGACAG

TGGGGCAGGAGGTCAGACGCCTGAGCCGAATCAGCGTTCAGCGGTGAGCGGGAAACTGTG

TCGTCCGATGAGTGAGGTTTGTTTTTAAACAAAAGTTGTTCTTGTCCCTAAAGGGACAGT

GGCGTCTTCCTGCAGCGCCTTCTGAACGCGGAGACACAGGTGTGCAGGGAATTCCAGAGT

CAGTTGCCAGATGGAGCGATGCCCTTATCCAGAGCAAGAGGGCGCCCTAGCCAGGCTAGA

CAGGGACCAGAGGGTCTGAGGTccccgcccctccccccctccccccccagcccccGGGTC

AGGGGTTGGCATAAGGCTGGGAAGCCACTCATGGAACCGTGACTCATTTACAAACACGCT

TAAATATGACATCACCACATGTAAGGGAGAGTCTGGCTCCATTTTGACCTTTGTAGACAG

CGCTGAGGCGTCCCCCCATCCCTTCCCCACCTCCGCACGTTTGGTACTTTACATAAAGAA

TCCCATGGCCTCCCAAGGGTCTGGCTCTGGGGGGATCTGAACCCTGCAGCCCTCGCCTGT

GCGCAGAAACTCGTCTCTGACCAGCTGTGAGCTTTGAGATGGGTTCAATAAAACTCAGGT

GTACATTTTACACTAGAAGGAAATTTATTTTACTTTCTGAGCATGACCTTAAGAGATTTT

GAAATGTAAACAGCTATGATATCCCTAAGGCAACTGAATTCTGATTTCCACCAATATACA

TATAAAAACAATGTTATGTAAATGTGAGCTGTACCTGAAAACTCTTTAAGATACTTAACA

CATTGGTTCAAAAAATTCCTAATTCATTTAtttgaatagctttactgagatataatacac

atattatacaaccctttaatgtgtataattcaatggttttcagtacattccgatatgtgc

aataatcatcacaacttttgaacatttttgtggcttcaaaaagaa

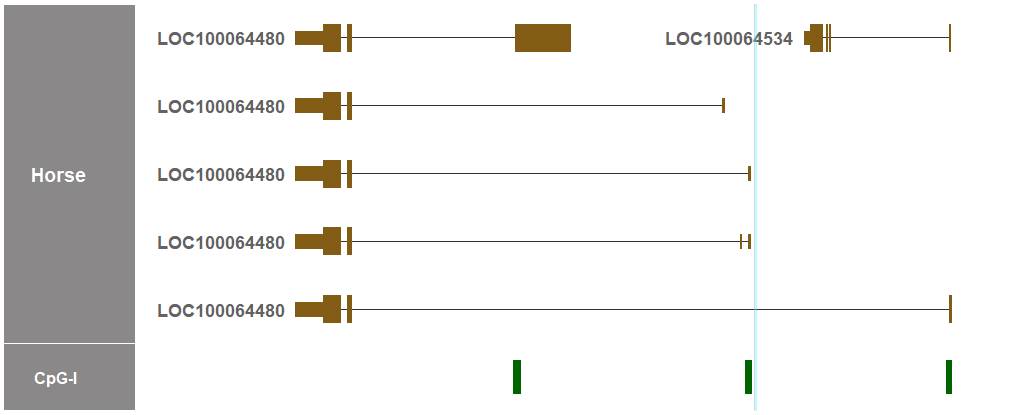

The detected region of LTR52 is indicated in cyan in both the sequence above and the genome visualizer.

5.3. Location of the detected region of LTR52 in the UCSC Genome Browser track

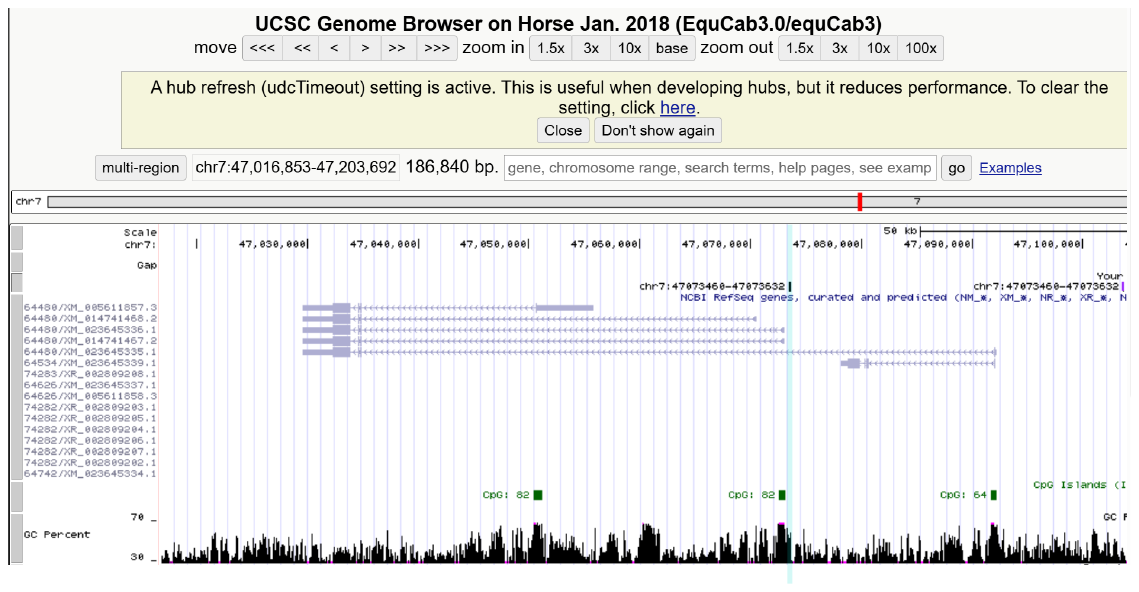

The above sequence in cyan was used an input for the UCSC BLAT search.

**6. Goat (*Capra hircus*) sequence**

6.1. Sequence search input (*ZNF791* E1 + 1kb upstream)

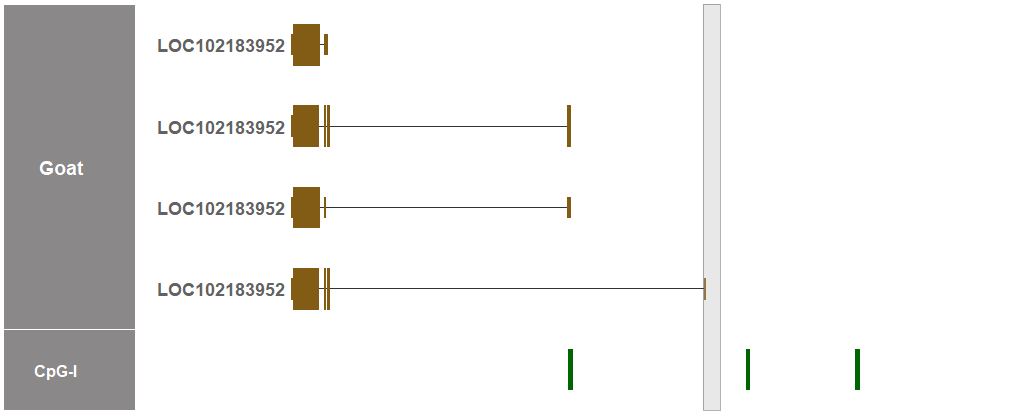

>chr7:9974100-9975196

CCTGTGCTCTTGGAGCCAAGAGGATGCAGCTCACATCCTGACTGCAGAGACCGTGTGCAG

GGGCAGCGGTGCTGAGGCTCCCAAAGCTAACCAGGTCAAGGTGTATCACGTCCCTTCAGA

GGGGAGGGCCAAGTGCAGCAAAGAAGCTGTTGAAATAGGCATCAAACACAACAAATTTAT

CCGAATATGTGAGAGCAGGGTATCTGCCAGCTGTTGACGGGATGGAGTCGGTAATTTCCA

CAATGTTGTGGACTGAGTACAGAGGTTCATTTGGTGTGACCACAACCTCAAGTGTTCAGT

AATCCACTGTCAGGTGTCATTCAGTCTTTTTAGATTTAAGAGCAGGCCAAattcagctgt

gaaaagaagtacAGGGAGTAATCACCTCTTTGCTAAATAATTTTTGtggggacttctttg

gtggcccagtggttaagaatctgcgtttccacttcagggggcacaggttcaatccctggg

tgaggaactAACATCCTAGGTGCTAtatggtgcggccaaaaaaattttttttttggtaat

ggtCTTCAATCCATCAAGGCCCTGTTTAATTTTCATTGGCCCATATTATGTAATTTAGAC

TTCCTCAAtggctcagtggggaaagaatctccctgcaaagGAGGAGAtaaaggagactca

ggtttgatccctgggttgggaagatcccctgcaaaaagaaatggcaacgcactctagtat

tcttgcctgaaaaatcccgtggacagaagagcctggcaggctacagtccatggggtcaca

aagagtcggatacaactgagcatgagcacaatAATGCAATTTACCTAGGGGGGTAGACTG

TGGGGTCCTATTTTGTCAAGCCAATTTGGAATTAGCAAAAAGTATTTAACTTTAATTTGT

GAATCATTGGTCCAGAGCTTTCCTGTTCACTATAGAAAACTTTAGGAAAATTGATCCTAT

GACCGTGGGAGAATTTAGGCAAGAAAAGAGTTGGCTGACAATCATTAAAGAAAGATATGC

CTATTCAGTAATTCCTCTAAGATTATGGGAAGTTCCTTGCTTAAGTTTGttaatgttgtt

tagtcgctcagttgtgt

6.2. Dfam results from profile HMMs

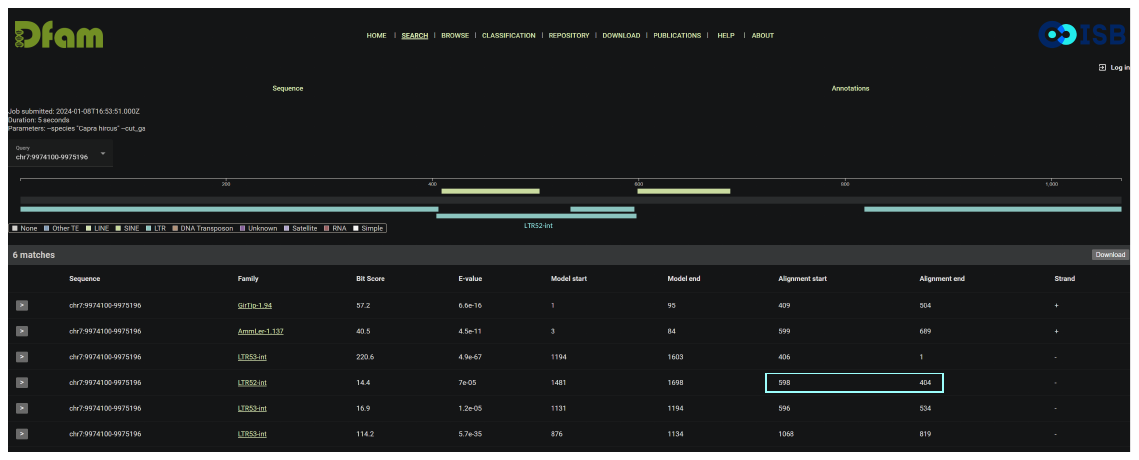

Using *Capra hircus* as a query organism and the Dfam curated threshold, the LTR52-int (Model) was

detected within the query input (Alignment) as shown in cyan.

>chr7:9974100-9975196

CCTGTGCTCTTGGAGCCAAGAGGATGCAGCTCACATCCTGACTGCAGAGACCGTGTGCAG

GGGCAGCGGTGCTGAGGCTCCCAAAGCTAACCAGGTCAAGGTGTATCACGTCCCTTCAGA

GGGGAGGGCCAAGTGCAGCAAAGAAGCTGTTGAAATAGGCATCAAACACAACAAATTTAT

CCGAATATGTGAGAGCAGGGTATCTGCCAGCTGTTGACGGGATGGAGTCGGTAATTTCCA

CAATGTTGTGGACTGAGTACAGAGGTTCATTTGGTGTGACCACAACCTCAAGTGTTCAGT

AATCCACTGTCAGGTGTCATTCAGTCTTTTTAGATTTAAGAGCAGGCCAAattcagctgt

gaaaagaagtacAGGGAGTAATCACCTCTTTGCTAAATAATTTTTGtggggacttctttg

gtggcccagtggttaagaatctgcgtttccacttcagggggcacaggttcaatccctggg

tgaggaactAACATCCTAGGTGCTAtatggtgcggccaaaaaaattttttttttggtaat

ggtCTTCAATCCATCAAGGCCCTGTTTAATTTTCATTGGCCCATATTATGTAATTTAGAC

TTCCTCAAtggctcagtggggaaagaatctccctgcaaagGAGGAGAtaaaggagactca

ggtttgatccctgggttgggaagatcccctgcaaaaagaaatggcaacgcactctagtat

tcttgcctgaaaaatcccgtggacagaagagcctggcaggctacagtccatggggtcaca

aagagtcggatacaactgagcatgagcacaatAATGCAATTTACCTAGGGGGGTAGACTG

TGGGGTCCTATTTTGTCAAGCCAATTTGGAATTAGCAAAAAGTATTTAACTTTAATTTGT

GAATCATTGGTCCAGAGCTTTCCTGTTCACTATAGAAAACTTTAGGAAAATTGATCCTAT

GACCGTGGGAGAATTTAGGCAAGAAAAGAGTTGGCTGACAATCATTAAAGAAAGATATGC

CTATTCAGTAATTCCTCTAAGATTATGGGAAGTTCCTTGCTTAAGTTTGttaatgttgtt

tagtcgctcagttgtgt

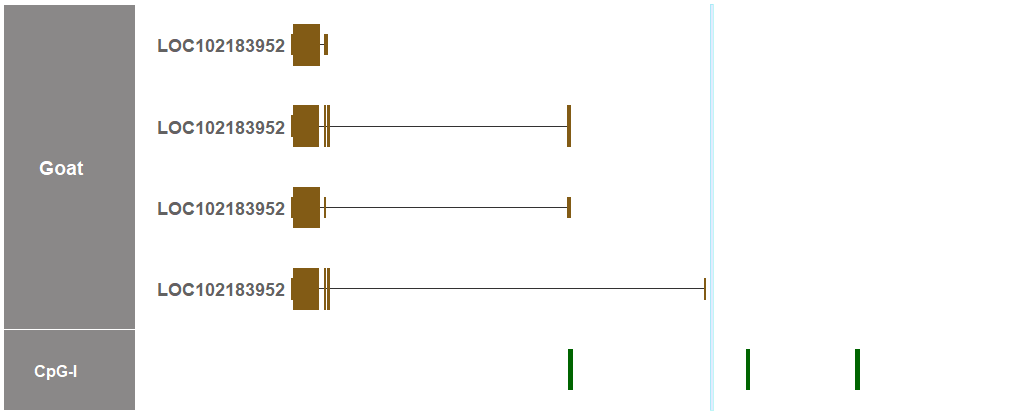

The detected region of LTR52-int is indicated in cyan in both the sequence above and the genome visualizer.

6.3. Location of the detected region of LTR52-int in the NCBI genome track

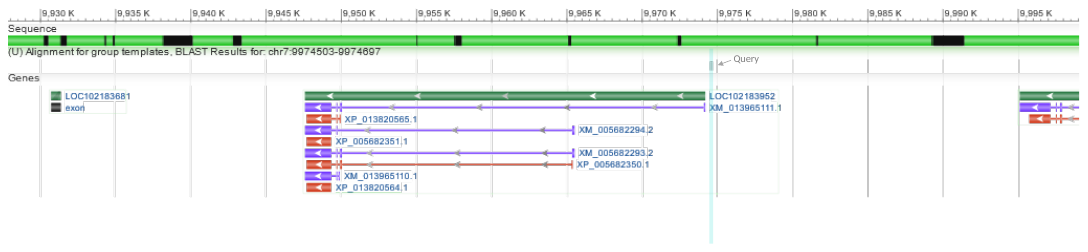

Capra hircus breed Yunnan black goat chromosome 7, CHIR_1.0, whole genome shotgun sequence

(NCBI Reference Sequence: NC_022299.1). The above sequence in cyan was used an input (Query)

for the NCBI BLAST search. The NCBI genome track was used because the CHIR_1.0 goat genome

was unavailable in the UCSC BLAT Search.

**7. Dog (*Canis lupus familiaris*) sequence**

7.1. Sequence search input (*ZNF791* E1 + 1kb upstream)

>chr20:49516920-49518052

CTCGAAGGGCCTCATCATCCCTTTGTGCATATGCATCCCCAGCGTTCCAGGCTCTGTAGC

TTCTCTTCATACAATGGCTCCTTCTGTGCTGGGTATGAGCAGGAAGGACTTCTGCAGAGG

CTCCTCAGGAAGAACCagctgggttcttttttttttttttttttttaaagattttattta

tttacatatgagagaccagagagagagaggcagagacacaggcagagggagaagcaggct

ccatgcagggagcccgggactccaggaccacaccctgggctgaaggcaggtgctaaaccg

ctgagccacccgggctgccccccccccttttttttaaagatttatttattcttgagagac

agagaaagagagagagagagagagagagaggcagagggagaagcaggctccacgcaggga

gcccgatatgggactggatcctgggactccaggatcatgccctgggccgaaggcaagcgc

tcaaccgctgagccacccagggatccctgaaccaGCTAGGTCTTGAAGGTATTTCCCTAT

GCAGGCTCAATGAGAAACGGGGTGGGGAGCCTACCTTGGGTCTAGTCATAGTCTTCTGCT

CCTAGTCCCATAGCTTTGGactatattgatatatttaatgACACAAACCCAGTATTAAAT

AATCTAGAAAAACCGCACAAATCCAATGTTTATGTagaaaagatggaagaagattgtaag

gcttttttttttccagctcaacAAGAAAATCCACAAGTATTTATTACTTTACGAAACAAT

GAGAGTAGATTATTATTTCCATGGCAAGAAGCTGTCCTATAAACAAACAtacttgaatat

tattttttgaagattttatttatttatttatgagagaccaagagagagacagagaggcag

agacacacaggcagagagggaagcaggctccatacagggagccggatgtgggactcaatc

ccgggtctccaggatcactccatgagctgaaggtggcgctaaaccgctgagccacccagg

ctagtACTTGAATATTCTAATTGCTGGAATTATATTGGAAACTGCTCCCCTTCGCcctca

cattaaaaaacaaaacaatacagacCTTAGAATATTATTACACCTTCTCAGGg

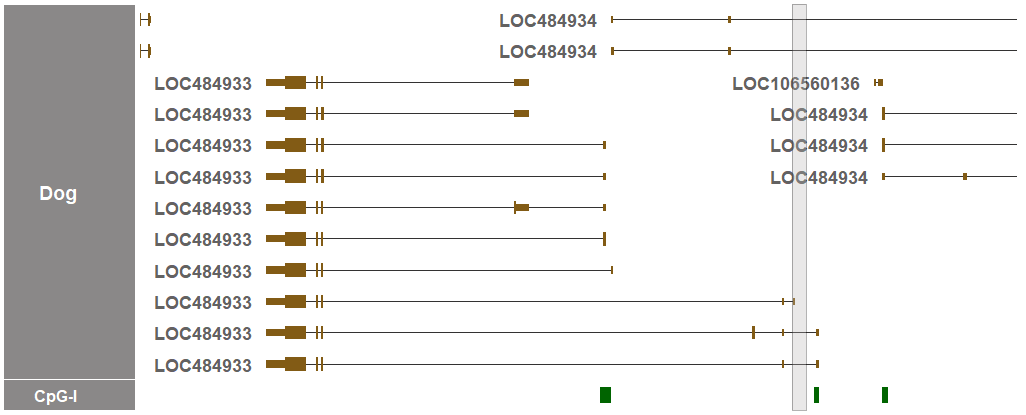

7.2. Dfam results from profile HMMs

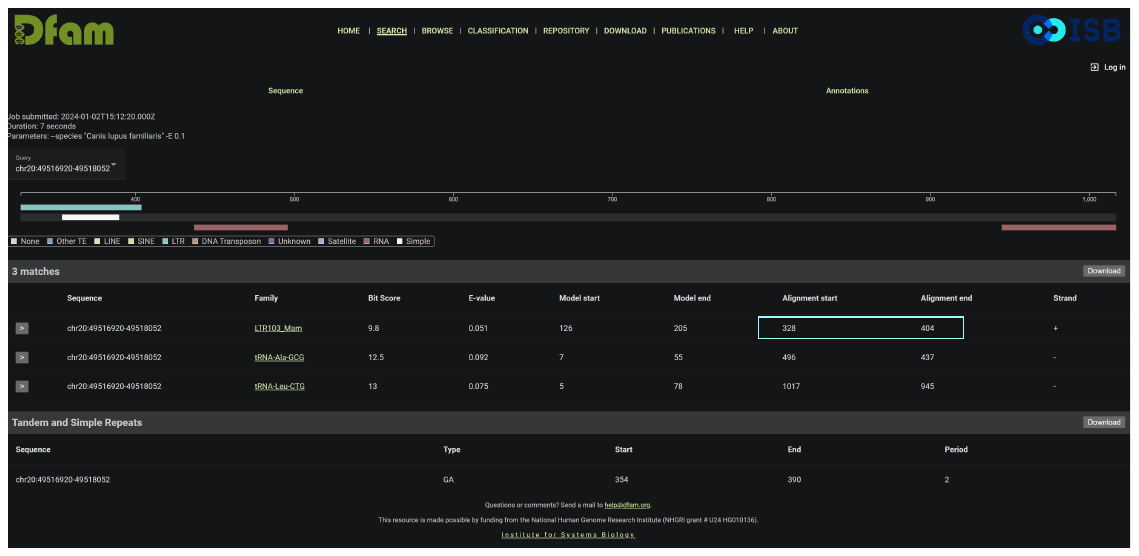

Using *Canis lupus familiaris* as a query organism and the E-value threshold, the LTR103-Mam (Model) was

detected within the query input (Alignment) as shown in cyan.

>chr20:49516920-49518052

CTCGAAGGGCCTCATCATCCCTTTGTGCATATGCATCCCCAGCGTTCCAGGCTCTGTAGC

TTCTCTTCATACAATGGCTCCTTCTGTGCTGGGTATGAGCAGGAAGGACTTCTGCAGAGG

CTCCTCAGGAAGAACCagctgggttcttttttttttttttttttttaaagattttattta

tttacatatgagagaccagagagagagaggcagagacacaggcagagggagaagcaggct

ccatgcagggagcccgggactccaggaccacaccctgggctgaaggcaggtgctaaaccg

ctgagccacccgggctgccccccccccttttttttaaagatttatttattcttgagagac

agagaaagagagagagagagagagagagaggcagagggagaagcaggctccacgcaggga

gcccgatatgggactggatcctgggactccaggatcatgccctgggccgaaggcaagcgc

tcaaccgctgagccacccagggatccctgaaccaGCTAGGTCTTGAAGGTATTTCCCTAT

GCAGGCTCAATGAGAAACGGGGTGGGGAGCCTACCTTGGGTCTAGTCATAGTCTTCTGCT

CCTAGTCCCATAGCTTTGGactatattgatatatttaatgACACAAACCCAGTATTAAAT

AATCTAGAAAAACCGCACAAATCCAATGTTTATGTagaaaagatggaagaagattgtaag

gcttttttttttccagctcaacAAGAAAATCCACAAGTATTTATTACTTTACGAAACAAT

GAGAGTAGATTATTATTTCCATGGCAAGAAGCTGTCCTATAAACAAACAtacttgaatat

tattttttgaagattttatttatttatttatgagagaccaagagagagacagagaggcag

agacacacaggcagagagggaagcaggctccatacagggagccggatgtgggactcaatc

ccgggtctccaggatcactccatgagctgaaggtggcgctaaaccgctgagccacccagg

ctagtACTTGAATATTCTAATTGCTGGAATTATATTGGAAACTGCTCCCCTTCGCcctca

cattaaaaaacaaaacaatacagacCTTAGAATATTATTACACCTTCTCAGGg

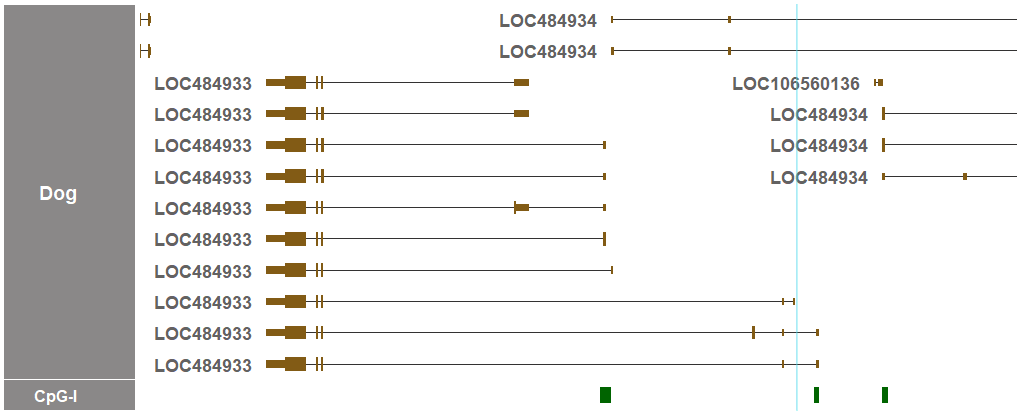

The detected region of LTR103-Mam is indicated in cyan in both the sequence above and the genome visualizer.

7.3. Location of the detected region of LTR103-Mam in the UCSC Genome Browser track

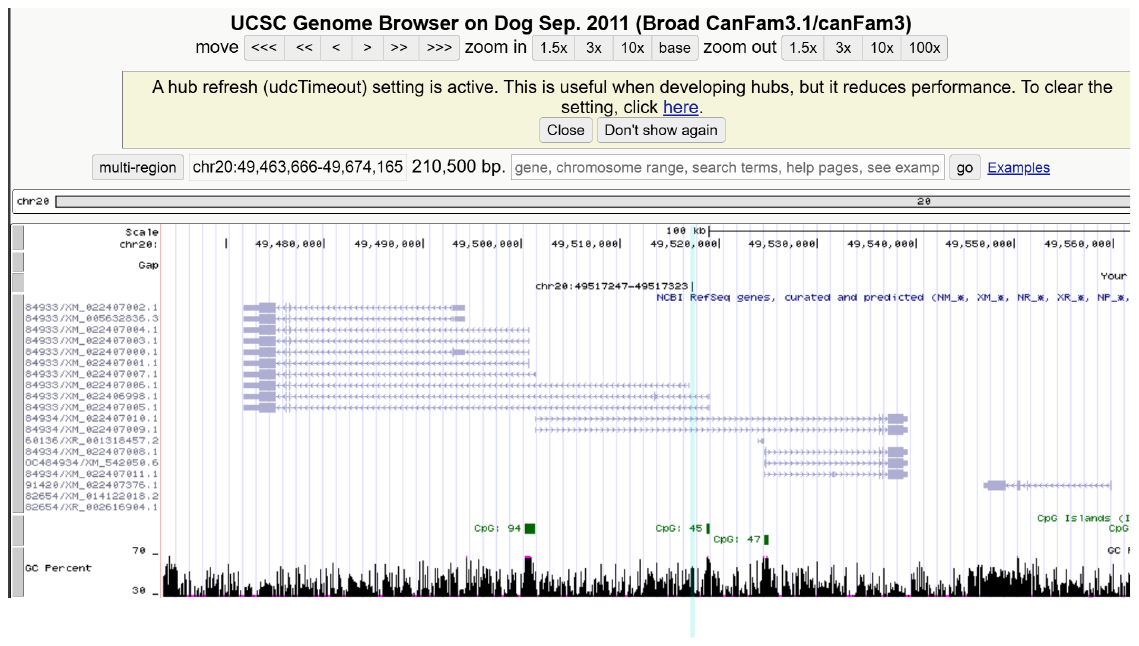

The above sequence in cyan was used an input for the UCSC BLAT search.
